# Decoding the mechanisms of cooperative DNA binding by the Paired-like homeodomain family

**DOI:** 10.1101/2025.10.29.685385

**Authors:** Brittany Cain, Connor Wasmund, Fiona C. Rowan, Brian Gebelein

**Affiliations:** Division of Developmental Biology, Cincinnati Children’s Hospital Medical Center, 3333 Burnet Ave, MLC 7007, Cincinnati, OH 45229, USA; Cincinnati Children’s Hospital Research Foundation, Graduate Program in Development, Stem Cells and Regenerative Medicine, Cincinnati, OH 45229, USA; Department of Pediatrics, University of Cincinnati College of Medicine, Cincinnati, OH 45229, USA

**Author notes:** denotes co-corresponding authors.

## Abstract

The 36 Paired-like homeodomain transcription factors are required for the development of many cell types, tissues, and organs as missense variants in 24 genes are associated with a variety of diseases and developmental disorders. How these factors identify distinct genomic targets using highly similar DNA binding domains is not fully understood. Here, we focus on determining how the Paired-like homeodomain factors gain DNA binding specificity by cooperatively binding palindromic sites spaced three base pairs apart (P3 site). Through structural, biochemical, and bioinformatic approaches, we define 11 rules that describe homeodomain residues that are critical, permissive, and inhibitory to cooperativity on the P3 site. Applying these rules, we successfully altered the cooperative behavior of Paired-like factors, identified residues that prevent the related Antennapedia class of homeodomains from binding cooperatively, and predict that thirty-eight disease-associated missense variants across ten Paired-like proteins alter cooperativity. Using quantitative DNA binding assays, we confirmed eleven of twelve of these disease-associated variants impact cooperativity but not DNA binding affinity. These findings reveal the importance of cooperativity in defining DNA binding specificity and highlight how missense variants associated with as many as fifteen different diseases can selectively disrupt cooperative DNA binding.

## 2. Introduction

The homeodomain (HD) family is one of the largest families of transcription factors (TFs) with over 200 members in the human genome (Lambert et al., 2018). HD proteins are classified by their highly conserved 60 amino acid helix-turn-helix domains, most of which bind highly similar AT-rich DNA sites (Badis et al., 2009; Berger et al., 2008; Jolma et al., 2013; Lambert et al., 2018). However, HD TFs typically bind distinct genomic targets and regulate different developmental pathways. Moreover, HD TFs have the most disease associated alleles reported in the Human Gene Mutation Database (HGMD) among all TF families, with numerous HD missense variants associated with a variety of diseases and birth defects (Kock et al., 2024). The way in which the human HDs acquire sufficient DNA binding specificity to regulate distinct biological outcomes, and how many of the pathogenic missense variants impact HD function is not well understood. Although computational tools have been developed to predict the consequences of single missense variants, a comprehensive analysis of 42 such prediction tools found that they largely fail to accurately identify variants that altered HD specificity (Kock et al., 2024), emphasizing the importance of elucidating the specificity-gaining mechanisms.

HD TFs can be categorized into distinct subfamilies based on sequence homology and the presence of additional conserved domains and motifs (Bürglin & Affolter, 2016). For example, the Paired/PAX subfamily of HD TFs are defined by having a second DNA binding domain, known as the Paired (Prd) domain and can thereby bind three types of DNA motifs using either the Prd domain, the HD, or both domains (Jun & Desplan, 1996). In contrast, the Paired-like factors, which constitute a family of 36 human genes that closely share a HD with the PAX factors but lack an associated Paired domain, encode only a single DNA binding domain. Thus, the Paired-like factors, much like the even larger Antennapedia (ANTP) class of HD factors that similarly lack a second DNA binding domain, must increase specificity by other mechanisms (Bürglin & Affolter, 2016).

One mechanism employed by TFs to increase their DNA binding specificity is the formation of complexes that bind longer recognition sequences. A well characterized example is the HOX factors within the ANTP subclass that specify distinct anterior-posterior fates along the body plan. HOX factors have relatively low DNA sequence specificity when bound alone but have distinct, increased specificity when complexed with PBX factors (Slattery et al., 2011). Previous work also revealed that some members of the Paired-like HDs form homo and heterodimers on a TAAT – NNN – ATTA P3 site (Cai, 1998; Cain et al., 2023; Jolma et al., 2013; Qu et al., 1999; Tucker & Wisdom, 1999; D. Wilson et al., 1993). To better understand which Paired-like HDs cooperatively bind DNA, we developed the Cooperativity Predictor pipeline to analyze HT-SELEX data to provide evidence that 8 of 23 Paired-like factors bind cooperatively to the P3 site (Cain et al., 2023). Sequence analysis and biochemical testing further revealed that several of the predicted non-cooperative Paired-like factors had residues at two HD positions (28 and 43) that were not conducive to P3 site cooperativity (Cain et al., 2023). However, these two rules were insufficient to accurately classify all the Paired-like HDs as either cooperative or non-cooperative. Moreover, a further limitation of the computational pipeline was that it treated cooperativity as a binary and did not capture the wide range of cooperativities seen in our biochemical validations (Cain et al., 2023).

In this study, we used comparative quantitative DNA binding assays to assess the degree of cooperativity of 15 of the 36 Paired-like HDs on P3 sites. By combining sequence conservation, biochemical analysis, and structural modeling, we identified key residues within the HD that are critical, permissive, and inhibitory to cooperativity and validated these residues via amino acid swaps to convert non-cooperative Paired-like HDs into cooperative proteins and vice versa. We then expanded our understanding of these mechanisms by examining the sequence constraint differences between the Paired-like factors and the related ANTP factors that are unable to bind cooperatively to the P3 site. This analysis revealed several additional residue changes that selectively disrupt cooperativity, including in residues that do not make direct protein-protein contacts but influence cooperativity through indirect mechanisms. In total, we defined 11 rules that either are critical for or contribute to cooperative DNA binding and apply these rules to both explain why the ANTP class of HDs fail to cooperatively bind P3 sites and predict which Paired-like HDs can cooperatively bind P3 sites. Lastly, we predicted 38 missense variants across 10 Paired-like proteins from HGMD would impact cooperativity. We tested 12 of these variants and found that all but one selectively impact cooperativity but maintain DNA binding. These findings reveal the importance of cooperativity in increasing DNA binding specificity and provide insights into how residues that do not contact DNA impact specificity. Moreover, these studies highlight how missense variants associated with at least 15 disease states can selectively disrupt cooperative DNA binding, providing new mechanistic understanding into how disease variants impact HD function.

## 3. Results

### Broad variation in cooperativity levels across the Paired-like class

While the Paired-like and ANTP subfamilies of HDs bind highly similar TAAT monomer DNA sites (Berger et al., 2008; Jolma et al., 2013; Lambert et al., 2018), HT-SELEX and biochemical studies found that only some, but not all, Paired-like HDs cooperatively bind TAAT – 3N – ATTA P3 sites (Cai, 1998; Cain et al., 2023; Jolma et al., 2013; Qu et al., 1999; Tucker & Wisdom, 1999; D. Wilson et al., 1993). Moreover, the prevalence and degree of cooperative binding to the P3 site across the 36 member Paired-like class is unknown. To better define DNA binding cooperativity across the Paired-like class, we sampled 15 human Paired-like factors (Figure 1A; highlighted in purple text) and performed quantitative electrophoretic mobility shift assays (EMSAs) using purified proteins containing their HDs and a P3 DNA probe. To select Paired-like factors for this analysis, we used two criteria: First, we created a dendrogram based on residue similarity across the entire 60 amino acid HD to identify factors from different subgroups. A key feature of Paired-like factors is that they can be classified based solely on the identity of the 50^th^ amino acid in the HD. The K50 HDs include members of the CRX/OTX, PITX, and GSC subfamilies, whereas the other Paired-like factors and nearly all ANTP class factors encode a Q50 HD. Since K50 and Q50 HDs are known to interact with DNA differently due to their preference for distinct nucleotides flanking the TAAT site (Treisman et al., 1989), we selected representatives of both the K50 and Q50 subfamilies for comparative studies. Second, we prioritized factors from each subgroup based on the prevalence of reported disease-associated HD missense variants on the HGMD (Stenson et al., 2017, 2020) (Figure 1A).

**Figure 1.**
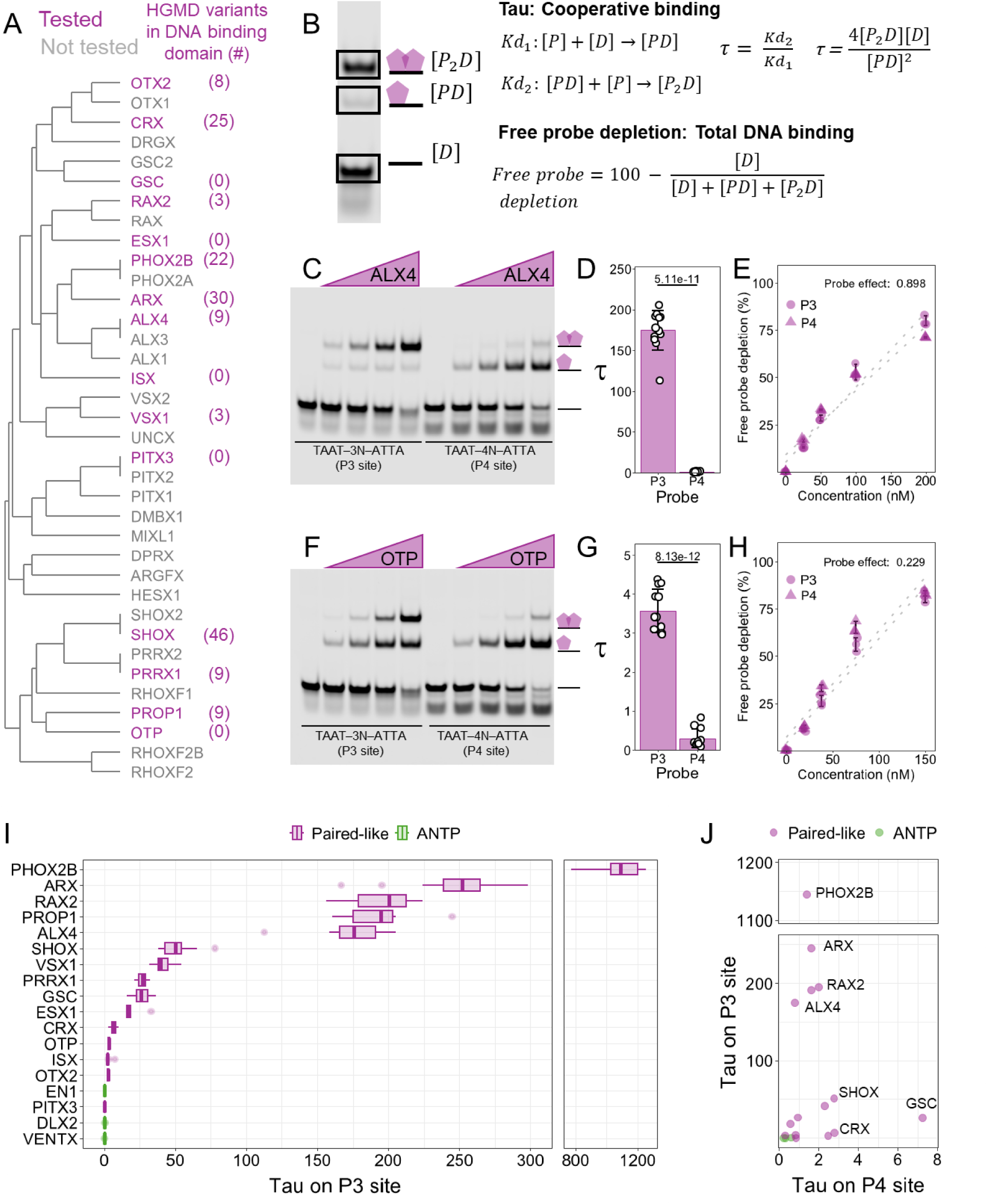
Paired-like factors vary in their ability to cooperatively bind the P3 site. **(A)** Dendrogram of the Paired-like class based on sequence similarity across the 60 amino acid HD. Proteins selected for cooperativity measurements are shown in purple text with the number of disease associated missense variants within each HD denoted in parentheses (Stenson et al., 2017, 2020). **(B)** Schematic describing calculation of cooperative binding (Tau) and total DNA binding (free probe depletion) using a portion of the EMSA for ALX4 shown in panel C. **(C)** ALX4 (amino acids = 209-274) was tested at 25, 50, 100, and 200 nM. **(D, G)** Tau cooperativity factors were calculated for every lane in which more than 5% of the probe was bound. Bars represent the average Tau for each protein and probe, and each dot represents a Tau from an independent binding reaction. Error bars represent standard deviation. Tau was compared via a Welch’s two-sided t-test (n = 12). **(D)** ALX4 binds with high cooperativity to the P3 site but not the P4 site. **(E, H)** The percentage of free probe depletion was calculated for each independent binding reaction. P3 probe depletion (circles) and P4 probe depletion (triangles) were compared via a two-way ANOVA and the effect of the protein variable is reported (5 concentrations; n = 3). **(E)** ALX4 had similar free probe depletion across the P3 and P4 site. **(F)** OTP (amino acids = 99-164) was tested at 18.75, 37.5, 75, and 150 nM. **(G)** OTP binds with low cooperativity to the P3 site but does not bind cooperatively to the P4 site. **(H)** OTP had similar free probe depletion across the P3 and P4 site. **(I)** Paired-like HDs bind to the P3 site with a large range of cooperativities. The Tau factors of 15 Paired-like (purple) and 3 ANTP HDs (green) were calculated from triplicate EMSAs (uncropped gels are shown in Supplementary Figure 1). Tau cooperativity factors were calculated for every lane in which more than 5% of the probe was bound. Lower and upper hinges denote the first and third quartiles, respectively. The lower and upper whiskers denote the most extreme value if there are no outliers or 1.5 times the interquartile range. **(J)** Paired-like factor cooperativity on the P3 site is spacer length dependent as HDs are unable to bind with high cooperativity to the P4 site. Each dot represents an average Tau for a protein across all concentrations and replicates.

For comparative DNA binding analysis across the Paired-like HD family, we used P3 probes with preferred spacer sequences for the Q50 versus the K50 HDs. For Q50 proteins, we used the most frequently bound 20mer in ALX4 HT-SELEX assays which contains a TAAT-TAG-ATTA P3 site (Cain et al., 2023; Jolma et al., 2013). For K50 proteins, we used a P3 site (TAAT-CCG-ATTA) that was preferably bound by the Prd K50 HD in SELEX assays (D. Wilson et al., 1993). To assess if cooperative binding was spacer length dependent, we also tested each factor on a P4 site in which a single G was added to the P3 spacer of each respective probe. As protein controls, we tested 3 ANTP factors (EN1, DLX2, and VENTX) that bind the ‘TAAT’ HD monomer site but were not predicted to cooperatively bind P3 sites (Cain et al., 2023). For each protein and binding site, we tested four protein concentrations in triplicate in EMSAs, and quantified band intensities to calculate both cooperativity and the total amount of bound probe (Figure 1B). To quantify cooperativity, we used a previously derived metric (D. Wilson et al., 1993), Tau, which measures the multiplier by which the binding of the first protein facilitates the binding of a second protein. Tau is the ratio of two equilibrium equations: Kd_2_ in which a dimer complex *[P_2_D]* forms from a monomer complex *[PD]* and Kd_1_ in which a monomer complex is formed from unbound DNA *[D]* and unbound protein *[P]* (Figure 1B). A tau greater than 1 is indicative of cooperative binding, a tau equal to 1 is independent binding, and a tau less than 1 indicates that the binding of the first protein inhibits the binding of the second. To quantify the relative overall binding affinity of probe/protein interactions, we measured free probe depletion which denotes the percentage of total DNA bound, independent of whether the DNA is bound as a monomeric or dimeric complex (Figure 1B). For example, ALX4 bound in a highly cooperative manner on the P3 site with a tau of 175 ± 24.2, while OTP bound with little cooperativity with a tau of 3.57 ± 0.56, but neither protein bound cooperatively to the P4 site (ALX4 tau: 0.792 ± 0.445; OTP tau: 0.290 ± 0.242) (Figure 1C-D, F-G). However, both proteins had similar free probe depletions on the P3 and P4 sites (Figure E, H), demonstrating that PROP1 and OTP can bind both DNA site types, but their ability to bind as a cooperative dimer on the P3 site varies.

Through this survey of 15 Paired-like HDs, we found a wide range of P3 site cooperativities across the Paired-like class with OTX2, ISX, and PITX3 showing little to no cooperativity, whereas ALX4, PROP1, RAX2, ARX, and PHOX2B have a Tau value greater than 100 (Figure 1I). Apart from OTX2 and PITX3, which were not highly cooperative on either site, we found that all Paired-like factors bound with a higher cooperativity to the P3 site compared to the P4 site (Figure 1J; Supplementary Figure 1). As expected, the ANTP factors, EN1, DLX2, and VENTX, did not bind cooperatively to the P3 site (Figure 1I) (Cain et al., 2023), and none of the ANTP factors preferred the P3 over the P4 site (Figure 1J; Supplementary Figure 1). Thus, while Paired-like factors share many highly conserved HD residues, they vary greatly in their ability to form cooperative DNA complexes on the P3 site.

### Residue changes across the Paired-like class contributes to cooperativity variation

To identify residue changes that contribute to this variation in cooperative DNA binding, we first used all 36 Paired-like sequences to generate an information content logo of conserved residues across the HD (Figure 2A, top). We then summarized the protein-protein contacts (red) and protein-DNA contacts (blue) using high resolution structural data of ALX4 (Cain et al., 2025) and Prd S50Q (D. S. Wilson et al., 1995) bound to a P3 site (Figure 2A, middle; structural alignment shown in Supplementary Figure 2). We next aligned the amino acid sequences of the tested Paired-like HDs in order of descending tau values (Figure 2A; bottom). Sequence analysis of HDs with low cooperativity (CRX, OTP, ISX, OTX2, and PITX3) revealed changes at five positions that make protein-protein contacts in at least one of the two available Paired-like structures: positions 2, 4, 28, 32, and 46 (bolded in red text, Figure 2A). Here, we interrogate the impact of these residues on cooperative DNA binding through molecular modeling, amino acid substitutions, and quantitative EMSAs. Our strategy is to swap key residues between cooperative and non-cooperative factors to identify those required to confer cooperativity and when possible, amino acid substitutions were tested in Paired-like factors that have disease associated variants at these positions. It is important to note that the cooperativity of Paired-like factors is highly variable (Figure 1E). Because of this, axes are set to best visualize the respective tested proteins and should not be compared across subpanels.

**Figure 2.**
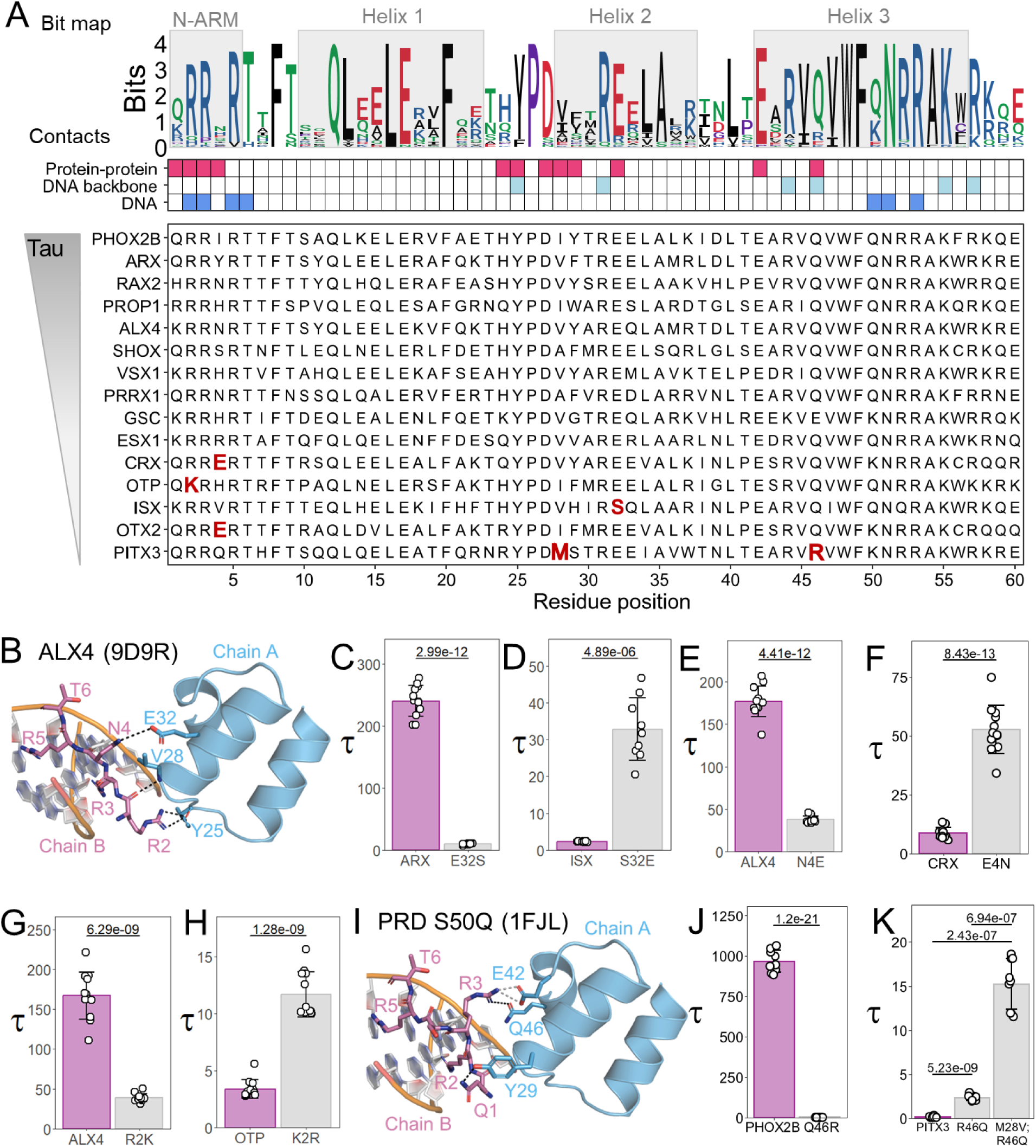
Paired-like HD sequence analysis, structural modeling, and quantitative DNA binding assays identify five HD residues that contribute to low cooperativity. **(A)** Sequence conservation and schematic depicting protein-DNA and protein-protein contacting residues that mediate Paired-like HD cooperativity on P3 sites. (Top) Information bit map across the 36 Paired-like factors reveals HD amino acid conservation. Residues are colored based on their biochemical properties (basic = blue; acidic = red; polar = green; special = purple; hydrophobic = black). (Middle) Contact grid highlights HD positions involved in protein-protein contacts (red), DNA backbone contacts (light blue), and DNA base contacts (dark blue). (Bottom) Sequence alignment of tested Paired-like factors in Figure 1 organized from high to low Tau values. Residues of interest at key positions are noted in red for the Paired-like HDs with low cooperativity. **(B,I)** PYMOL schematics of key protein-protein contacts that mediate cooperative DNA binding by the ALX4 and Prd S50Q HDs on a P3 site. Chain A and B are shown in blue and purple respectively. Hydrogen bonds are denoted with black dashed lines. Salt bridges are denoted with gray dashed lines. **(C-H; J-K)** Tau values were calculated from triplicate EMSAs using each lane with more than 5% of the probe bound. Bars represent the average Tau for each protein across all tested concentrations, and each dot represents a Tau from an independent binding reaction. Error bars represent standard deviation. Tau values were compared with a Welch’s or standard t-test depending on the result of the Levene’s test for variance. **(K)** Tau was compared with a one-way ANOVA (3 proteins; n = 9; 4.07e-16). Pairwise two-sided Welch’s t-tests were performed with a Holm correction for multiple comparisons. Gels are shown in Supplementary Figure 3.

Of the tested Paired-like factors, all have a glutamic acid at position 32 except ISX, which has a S32. In the ALX4 structure, Chain A E32 interacts with the N4 residue of Chain B (Figure 2B), whereas in Prd S50Q, Chain B E32 and Chain A R3 form a salt bridge. Given the importance of E32 in both structures, we tested an E32S mutation within the highly cooperative ARX protein. EMSAs on the P3 site revealed that while wildtype ARX had a Tau of 240 ± 25, ARX E32S had a 24-fold decrease in Tau (10 ± 0.97; Figure 2C). Inversely, the cooperativity of ISX increased ∼16-fold with an S32E mutation (ISX wildtype: 2.34 ± 0.15; ISX S32E: 32.8 ± 8.5; Figure 2D).

As mentioned, ALX4 E32 of Chain A interacts with N4 of Chain B, which is a residue that is not highly conserved across the Paired-like class (Figure 2A). However, very few Paired-like factors have a glutamic acid at position 4 except for the weakly cooperative CRX and OTX2 factors from those surveyed (Figure 2A, bold red text). In contrast to the ALX4 structure, S4 in the Prd S50Q structure forms no polar interactions and minimal non-polar interactions with the second protein (D. S. Wilson et al., 1995). Despite this difference, a large, negatively charged E4 residue is predicted to clash with E32 of the opposing chain and thereby inhibit cooperativity. To test this prediction, we created an ALX4 N4E mutation and found that it reduced cooperativity ∼4.5-fold (ALX4 wildtype: 176 ± 18.0; ALX4 N4E: 37.8 ± 4.33; Figure 2E). Inversely, a CRX E4N mutation increased cooperativity 6-fold (CRX wildtype: 8.70 ± 2.34; CRX E4N: 52.7 ± 10.2; Figure 2F). Thus, residue differences at positions 4 and 32 explain the low cooperativity levels of ISX, OTX2, and CRX compared to the other Paired-like factors. Note, a summary of the role of each tested residue on cooperative DNA binding and the impact of sequence variation at that HD position is found in Table 1.

**Table 1.** Requirements of P3 site cooperativity identified from Figure 2-Figure 5 and past studies.

| HD position | Rule | Importance | Source | Structural Rational |
| --- | --- | --- | --- | --- |
| 2 | Must be an R | Contributory | This study (Cain et al., 2025) | R2 is required to either facilitate the protein-protein interface or insert into the minor groove. |
| 3 | Must be an R | Critical | (Cain et al., 2025) | R3 is required to either facilitate the protein-protein interface or insert into the minor groove. |
| 4 | Cannot be an E | Contributory | This study | E4 clashes with E32 of the partner protein |
| 26 | Must be a P | Critical | This study | Branched residues cannot fit in the narrow hydrophobic core between helix 1 and 2 |
| 28 | Cannot be an R | Critical | (Cain et al., 2025; D. S. Wilson et al., 1995) | R clashes with the basic residues in the N-terminal ARM |
| 32 | Must be an E | Critical | This study | E32 facilitates hydrogen bonds or salt bridges with the N-terminal ARM |
| 42 | Must be an E | Critical | (Cain et al., 2025; Zheng et al., 2024) | E42 forms a salt bridge with R3 |
| 43 | Cannot be an R | Critical | (D. S. Wilson et al., 1995) | R43 clashes where the recognition helices meet |
| 44 | Cannot be a Q | Critical | This study | Q44 would be unable to form the water mediated hydrogen bond with E42 |
| 46 | Cannot be an R or K | Critical | This study | R46 or K46 clashes with R3 in the N-terminal ARM |
| 50 | Cannot be a K | Contributory | This study (D. Wilson et al., 1993; Zheng et al., 2024) | Unknown; See discussion |

While R2 is largely conserved across the Paired-like class, OTP has a lysine in this position. In the Prd S50Q structure, the R2 residue of both chains inserts into the minor groove of DNA, and R3 mediates protein-protein interactions (D. S. Wilson et al., 1995). Conversely, in ALX4, the R2 residue of Chain A inserts into the minor groove of DNA, while Chain B R2 contacts Y25 and V28 of Chain A, revealing that this residue can directly contribute to cooperativity (Figure 2B) (Cain et al., 2025). While lysine is similarly sized and charged as arginine, lysine residues typically make fewer minor groove contacts (Rohs et al., 2009) and are less frequent at protein-protein interfaces (Tsai et al., 1997). To determine if K2 impacts cooperativity, we tested an ALX4 R2K mutation and found it decreased cooperativity more than four-fold (ALX4 wildtype: 167 ± 29.4; ALX4 R2K: 39.0 ± 5.08; Figure 2G). Note, the cooperativity of ALX4 R2K was still two-fold higher than a published ALX4 R2A mutation (Cain et al., 2025), which is consistent with lysine being better able to fulfill protein-DNA and protein-protein contacts compared to alanine. To assess if K2 inhibits cooperativity in OTP, we compared OTP K2R to wildtype and found that this mutation increased cooperativity more than three-fold (OTP wildtype: 3.39 ± 0.804; OTP K2R: 11.7 ± 1.98; Figure 2H).

While most Paired-like HDs have a Q46 residue, the non-cooperative PITX3 factor has an R46. In the Prd S50Q structure, Chain A Q46 forms a hydrogen bond with Chain B R3 (Figure 2I) (D. S. Wilson et al., 1995), while this residue makes DNA backbone contacts in the ALX4 structure (Cain et al., 2025). In the PITX factors, R46 is predicted to clash with R3 due to size and electrostatic repulsion. Intriguingly, a Q46R missense variant in the highly cooperative PHOX2B factor has been associated with congenital central hypoventilation syndrome (CCHS) (Trochet, O’brien, et al., 2005), consistent with this variant disrupting PHOX2B function. We tested the importance of Q46 in cooperativity using PHOX2B wildtype and Q46R and found that this variant completely disrupted cooperativity (PHOX2B wildtype: 968 ± 67.4; PHOX2B Q46R: 1.27 ± 0.832) (Figure 2J). Conversely, we compared PITX3 wildtype and R46Q and found it significantly increased cooperativity, but only to a Tau of 2.33 (PITX3 wildtype: 0.157 ± 0.094; PITX3 R46Q: 2.33 ± 0.315; Figure 2K). This finding suggests that other residues contribute to PITX3’s low cooperativity. Consistent with this idea, PITX3 has a methionine at position 28, and we previously found that a V28M mutation in ALX4 significantly reduced cooperativity due to clashing with the N-terminal Arginine Rich Motif (ARM) of the second protein (Cain et al., 2023). Hence, we created a double PITX3 M28V;R46Q mutation and found that it increased cooperativity 6.5-fold compared to the PITX3 R46Q mutation and 100-fold compared to wildtype PITX3 (PITX3 M28V; R46Q: 15.3 ± 2.85; Figure 2K).

### HD position 50 impacts both DNA binding cooperativity and specificity

Using structural information and comparative sequence analysis, we identified 5 HD positions (2, 4, 28, 32, and 46) that differ between highly cooperative versus weakly and non-cooperative Paired-like factors (Figure 2; Table 1). Sequence alignment and cooperativity measurements further revealed that all tested Paired-like proteins with a lysine at position 50 had Tau values less than 30, whereas Q50 Paired- like TFs often had Tau values greater than 50 (Figure 1I; Figure 2A). As mentioned above, position 50’s importance in DNA binding specificity has been long established. A K50 HD prefers a TAAT<u>CC</u> sequence (Treisman et al., 1989) as K50 forms hydrogen bonds with the guanines on the opposing strand (Tucker-Kellogg et al., 1997), while Q50 typically fails to form direct contacts with flanking bases (Kissinger et al., 1990). Previous work also connected position 50’s residue with both spacer length, spacer nucleotide content preference, and cooperativity levels (D. Wilson et al., 1993; D. S. Wilson et al., 1996). To test the impact of having a Q50 vs K50 residue in cooperative DNA binding by Paired-like TFs, we mutated the highly cooperative ALX4 Q50 into a K50 and the weakly cooperative CRX K50 into a Q50. Since CRX has the E4 residue that inhibits cooperativity (see Figure 2F), we also generated a CRX E4N;K50Q double mutation and tested each on P3 sites with linker sequences optimized for a Q50 (TAAT – TAG – ATTA) or K50 (TAAT – CCG – ATTA) HD. On the Q50 probe, wildtype CRX (Tau = 0.98 ± 0.68) and ALX4 Q50K (Tau = 2.02 ± 1.08) were unable to bind with high cooperativity like wildtype ALX4 (Tau = 156.9 ± 19.6) (Figure 3A), whereas on the K50 probe, ALX4 Q50K (Tau = 47.7 ± 4.04) bound cooperatively, but significantly less than wildtype ALX4 (Tau = 343 ± 41.6) (Figure 3A). These findings suggest that a K50 protein is not conducive to strong cooperative binding on either P3 site; a result that is consistent with past studies (D. Wilson et al., 1993; Zheng et al., 2024). We also assessed the relative affinity of each protein for each probe by quantifying free probe depletion (Figure 1B). As expected, wildtype ALX4 bound with a higher affinity compared to ALX4 Q50K or CRX on the Q50 P3 probe (Figure 3B). However, we surprisingly found that wildtype ALX4 bound the K50 P3 dimer site with similar affinity as CRX and ALX4 Q50K (Figure 3B). In contrast, wildtype ALX4 had very low affinity for a monomer K50 site compared to its preferred Q50 monomer site (Figure 3C). These findings highlight how cooperativity allows for strong binding to a sub-optimal site. Conversely, due to the lack of strong cooperative binding, the K50 proteins were unable to maintain high DNA binding affinity for the Q50 P3 site (Figure 3B).

**Figure 3.**
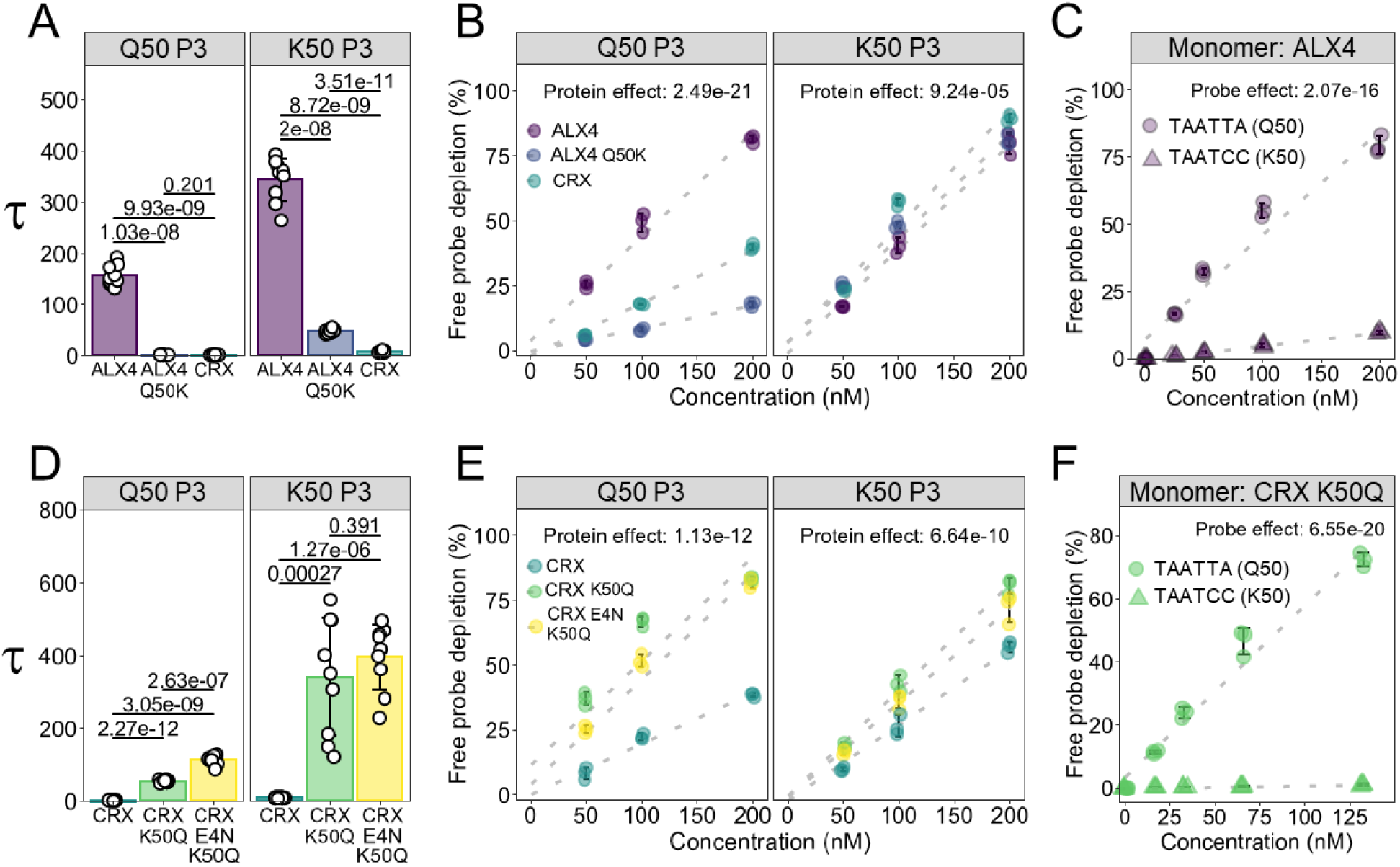
Dramatic differences in cooperativity between Q50 and K50 HD proteins. **(A)** Comparative cooperativity analysis of ALX4 WT, ALX4 Q50K, and CRX on the Q50 P3 probe (TAAT – TAG – ATTA) and K50 P3 probe (TAAT – CCG – ATTA). Proteins compared with a one-way ANOVA (3 proteins; n = 3) (Protein effect on Q50 probe: p = 4.14e-18; Protein effect on K50 probe: p = 1.78e-20). Pairwise comparisons were determined with a two-sided Welch’s t-test with a Holm multiple comparisons correction. **(B)** Free probe depletion was calculated for each binding reaction and compared via a two-way ANOVA. The effect of the protein variable is reported (3 proteins; 3 concentrations; n = 3). **(C)** ALX4 WT has high affinity for the Q50 monomer site (TAATTA), denoted with circles, but low affinity for the K50 monomer site (TAATCC), denoted with triangles. Free probe depletion was compared via a two-way ANOVA, and the effect of probe is reported (2 probes, 5 concentrations, n = 3). **(D)** Comparative cooperativity analysis of CRX WT, CRX K50Q, and CRX E4N;K50Q using Tau values calculated from triplicate EMSAs using each probe with a one-way ANOVA (3 proteins; n = 3) (Protein effect on Q50 probe: p = 2.5e-23; Protein effect on K50 probe: p = 1.59e-13). Pairwise comparisons were determined with a two-sided Welch’s t-test with a Holm multiple comparisons correction. **(E)** See B for details. **(F)** CRX K50Q has a higher affinity for the Q50 monomer site (TAATTA), denoted with circles, compared to the K50 monomer site (TAATCC), denoted with triangles. Free probe depletions were compared via a two-way ANOVA, and the effect of probe is reported (2 probes, 5 concentrations, n = 3).

Consistent with the Q50 residue being more compatible with cooperative DNA binding in ALX4, the CRX K50Q (Q50 P3 Tau = 54.6 ± 2.72; K50 P3 Tau = 340 ± 161) and CRX E4N;K50Q (Q50 P3 Tau = 112 ± 12.1; K50 P3 Tau = 395 ± 90.1) proteins displayed at least a 30-fold increase in cooperativity on both P3 probes compared to wildtype CRX (Q50 P3 Tau = 0.936 ± 0.488; K50 P3 Tau = 8.94 ± 1.78) (Figure 3D). This cooperative behavior enabled CRX K50Q and CRX E4N;K50Q to bind with similar affinity to the less ideal K50 P3 site (Figure 3E), whereas CRX K50Q had minimal binding on the preferred TAATCC K50 monomer site (Figure 3F). Taken together, these findings suggest that K50 residues are not conducive to strong cooperative binding and that the ability to bind DNA cooperatively can enable a protein to bind weaker half-sites with relatively high affinity.

### Sequence constraints between ANTP and Paired-Like HDs identifies additional residues critical for cooperative DNA binding

While structural analysis offers valuable insight into which residues form protein-protein contacts, having only two dimer structures across the Paired-like family provides a limited understanding as to which residues are permissive or inhibitory for cooperativity. To identify additional residues that make contributions to cooperativity, we compared sequence constraints of the Paired-like class versus the ANTP class, which bind highly similar TAAT monomer sites (Bürglin & Affolter, 2016; Jolma et al., 2013) but fail to cooperatively bind P3 sites (Figure 1I) (Cain et al., 2023). In theory, residue differences between these classes could reveal residues that are conducive (Paired-like) or inhibitory (ANTP class) to P3 site cooperativity, while such differences are expected to maintain monomer site binding. Based on past classifications (Bürglin & Affolter, 2016; Holland et al., 2007), we included 102 ANTP HDs and 36 Paired-like HDs in our analysis and performed a multiple sequence alignment of the 60 amino acid HD for all 138 factors. We converted this alignment to a similarity distance matrix and calculated the Euclidean distance between each factor. The HDs of all 36 Paired-like factors cluster together (Figure 4A) and show higher similarities to each other compared to the ANTP class (Figure 4B-C). To systematically identify differences between classes, we created a differential bit map by subtracting the relative content information of the ANTP class from the Paired-like class (Figure 4D). In this plot, positive bit information correlates with sequence constraint of the Paired-like class while negative bit information correlates with sequence constraint of the ANTP class. We found several areas of interest based on this analysis, including residue differences in the recognition helix as well as large differences in the turn between helix 1 and 2. As proof of concept, the differential bit analysis revealed that E32 is preferentially constrained in the Paired-like class, which is consistent with our findings on its importance in mediating protein-protein interactions that contribute to cooperative binding (Figure 2C-D; highlighted in gray in Figure 4D). In contrast, ANTP HDs largely have a hydrophobic residue at position 32 (Figure 4C-D). Similarly, studies on Prd S50Q found that an arginine at positions 28 and 43, which are the preferred residues in ANTP HDs (highlighted in gray in Figure 4D), have been previously shown to disrupt cooperativity (D. S. Wilson et al., 1995).

**Figure 4.**
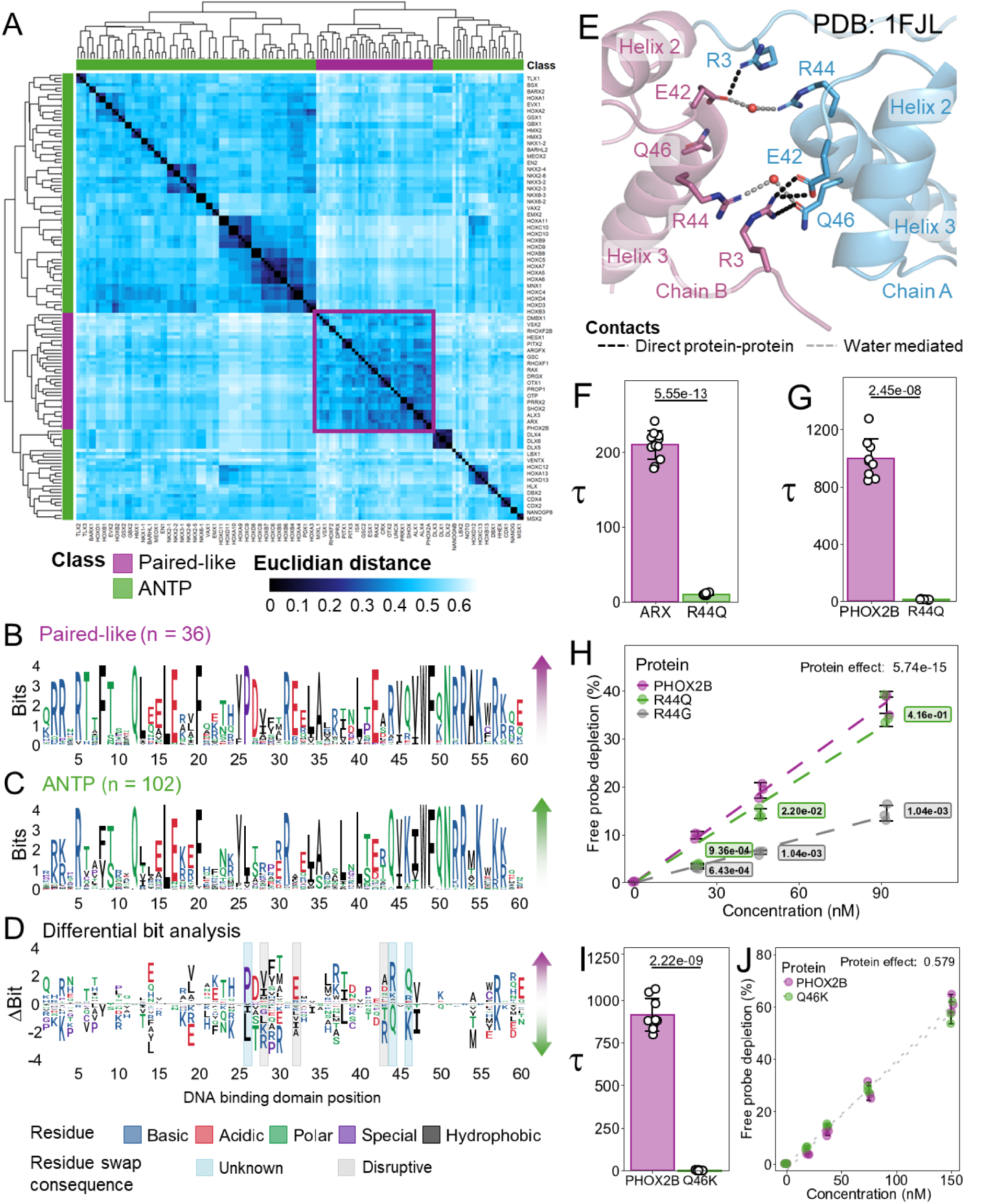
Paired-like and ANTP classes have different sequence constraints in the turn between helix 1 and helix 2 and select residues of the recognition helix. **(A)** The 60 amino acid HDs of the Paired-like class (purple) cluster together compared to the HDs of the ANTP class (green). Protein sequences were clustered via the complete linkage method. Heatmap shows the Euclidian distances derived from the sequence alignments of the 60 amino acid DNA binding domains of 102 ANTP factors and 36 Paired-like factors, in which the smaller Euclidean distances correspond with the highest similarity between proteins. **(B-C)** Information bit map of the DNA binding domains of Paired-like (B) and ANTP (C) factors. Residues are colored based on amino acid properties. **(D)** Sequence constraint changes between the Paired-like and ANTP classes. Bit information of the ANTP class was subtracted from the Paired-like class. Positive bit information correlates with sequence constraint of the Paired-like class. Negative bit information correlates with sequence constraint of the ANTP class. 3 HD positions with differential sequence constraint have been previously confirmed to alter cooperativity (denoted with gray shading), and 3 HD positions were selected for further testing (denoted with blue shading). **(E)** PYMOL schematic of the protein-protein contacts between the recognition helices and N-terminal ARMs in the Prd S50Q crystal structure (1FJL). Chain A is shown in blue and Chain B is shown in purple. Direct protein-protein contacts denoted with black dashed lines, and water mediated contacts denoted with gray dashed lines. **(F, G, I)** Tau cooperativity factors were calculated from triplicate EMSAs for every lane in which more than 5% of the probe was bound in quantitative EMSAs. Bars represent the average Tau for each protein, and each dot represents a Tau from an independent binding reaction. Error bars represent standard deviation. **(F)** Tau was compared via two-sided Welch’s t-test (n = 12). **(G)** Tau was compared via a one-way ANOVA with a Holm multiple comparisons correction (n = 9). Tau was omitted for PHOX2B R44G due to the lack of DNA binding. **(H,J)** Free probe depletion was calculated for each binding reaction. Free probe depletions were compared via a two-way ANOVA and effect of the protein variable is reported (3 concentrations; n = 3) **(H)** A Holm post-hoc was used to determine pairwise comparisons. Only comparisons to the wildtype protein are shown. **(I)** Tau was compared with a two-sided Welch’s t-test (PHOX2B wildtype n = 9; PHOX2B Q46K n = 12). **(F-J)** Uncropped gels used for quantitation are shown in Supplementary Figure 5.

### The role of HD positions 44 and 46 in cooperativity

Two obvious differences between Paired-like and ANTP HDs occur at positions 44 and 46. 30 of 36 Paired-like factors have an R44, whereas 99 of 102 ANTP factors have Q44. In the Paired S50Q structure, R44 makes water mediated contacts with either E42 or Q46 (Figure 4E) (D. S. Wilson et al., 1995). Most water molecules could not be modeled in the ALX4 structure due to insufficient resolution, so it is unknown whether these interactions are consistent across both structures. Intriguingly, clinical genetic studies identified R44Q missense variants associated with disease in both ARX and PHOX2B (Thai et al., 2020; Trochet, O’brien, et al., 2005), suggesting that this Paired-like to ANTP class residue swap sufficiently disrupts HD function to cause pathogenesis. To test how these variants impact DNA cooperativity, we performed comparative EMSAs with ARX wildtype (R44) and the ARX R44Q variant associated with intellectual disability and epilepsy (Thai et al., 2020). These studies revealed that R44Q reduced cooperativity over 20-fold (ARX WT: 209±18.7 vs ARX R44Q: 9.6±1.4; Figure 4F) with minimal changes to overall DNA binding affinity as assessed by amount of probe depletion (Supplementary Figure 5A). We similarly found that PHOX2B R44Q, which is associated with central hypoventilation syndrome (Trochet, O’brien, et al., 2005), strongly decreased cooperativity (PHOX2B wildtype: 996 ± 138; PHOX2B R44Q: 11.9 ± 1.94; Figure 4G). Interestingly, a distinct PHOX2B missense variant at this position (R44G) was found to be associated with neuroblastoma with Hirschsprung disease (Trochet et al., 2004), and this variant was previously found to decrease binding to monomer DNA sites (Trochet, Hong, et al., 2005). We similarly found that PHOX2B R44G strongly decreased overall DNA binding to the HD sites within the P3 probe, whereas PHOX2B R44Q maintained similar DNA binding affinity as wildtype PHOX2B (Figure 4H). Thus, disease variants that alter a Paired-like residue to an ANTP class residue can selectively disrupt cooperative DNA binding without dramatically impacting the ability to bind DNA, whereas a different missense variant at the same position can impact overall DNA binding.

Like position 44, residue 46 has distinct constraints between Paired-like and ANTP HDs with 29 of 36 Paired-like factors encoding a Q46, whereas 94 of 102 ANTP factors have a K46. As mentioned above, Q46 in Chain A of the Prd S50Q structure directly contacts R3 of Chain B and forms a water mediated contact with R44 of Chain B (Figure 4E). To determine the effect of K46 on cooperativity, we compared PHOX2B wildtype to PHOX2B Q46K and found that this variant fully disrupted cooperativity (Figure 4I; PHOX2B wildtype: 913.6 ± 94.7; PHOX2B Q46K: 1.4 ± 0.8), while having no impact on DNA binding affinity (Figure 4J). These data are consistent with our previous finding that PHOX2B Q46R completely disrupts cooperativity (Figure 2J), demonstrating that a positively charged residue at this position is not conducive to cooperativity, most likely due to clashing with R3 (Figure 4E; Table 1).

### The relationship between HD helical spacing and P3 site cooperativity

One of the most consistent differences between Paired-like and ANTP factors occurs at position 26, as all Paired-like factors have a proline, whereas 101 of 102 ANTP factors have a branched hydrophobic residue. Moreover, a P26L missense variant in ARX has been associated with neurological disease (Strømme et al., 2002), suggesting this Paired-like to ANTP HD swap disrupts protein function. Analysis of the Paired-like dimer structures reveals that P26 sits between five positions that mediate protein-protein contacts that contribute to cooperative DNA binding. However, P26 does not make any direct interactions (Figure 2A). Comparative EMSA analysis between ARX P26 and ARX P26L revealed an ∼10-fold decrease in cooperativity with only minor changes to DNA binding affinity as measured by probe depletion (Figure 5A; Supplementary Figure 6A). However, the mechanism behind this loss of cooperativity is unclear. Although L26 is accommodated and contributes to the hydrophobic core in structures of ANTP members (Figure 5B; Supplementary Figure 6C), *in silico* mutagenesis reveals that a leucine is predicted to clash with position 30 in helix 2 in the ALX4 structure (Figure 5C). To understand how a leucine fits within the hydrophobic core of ANTP factors but not Paired-like factors, we used published structures to compare the pairwise intrachain alpha carbon distances across the entire DNA binding domain of seven Paired-like proteins and five ANTP proteins. This analysis revealed that helices 1 and 2 are ∼2 angstroms closer together in the Paired-like structures, and this proximity change begins at position 26 (Figure 5D; Supplementary Figure 7). To systematically compare distances between the helices, we selected four inner helical pairs for each helix-helix combination. We found that all four helix 1 and 2 pairings were closer in proximity in the Paired-like structures compared to the ANTP structures, whereas there was no difference in proximity between residue pairs between helix 1 and 3 or helix 2 and 3 (Figure 5E; Supplementary Figure 8). This analysis provides insight into how a leucine can spatially fit in the ANTP hydrophobic core and not the Paired-like hydrophobic core.

**Figure 5.**
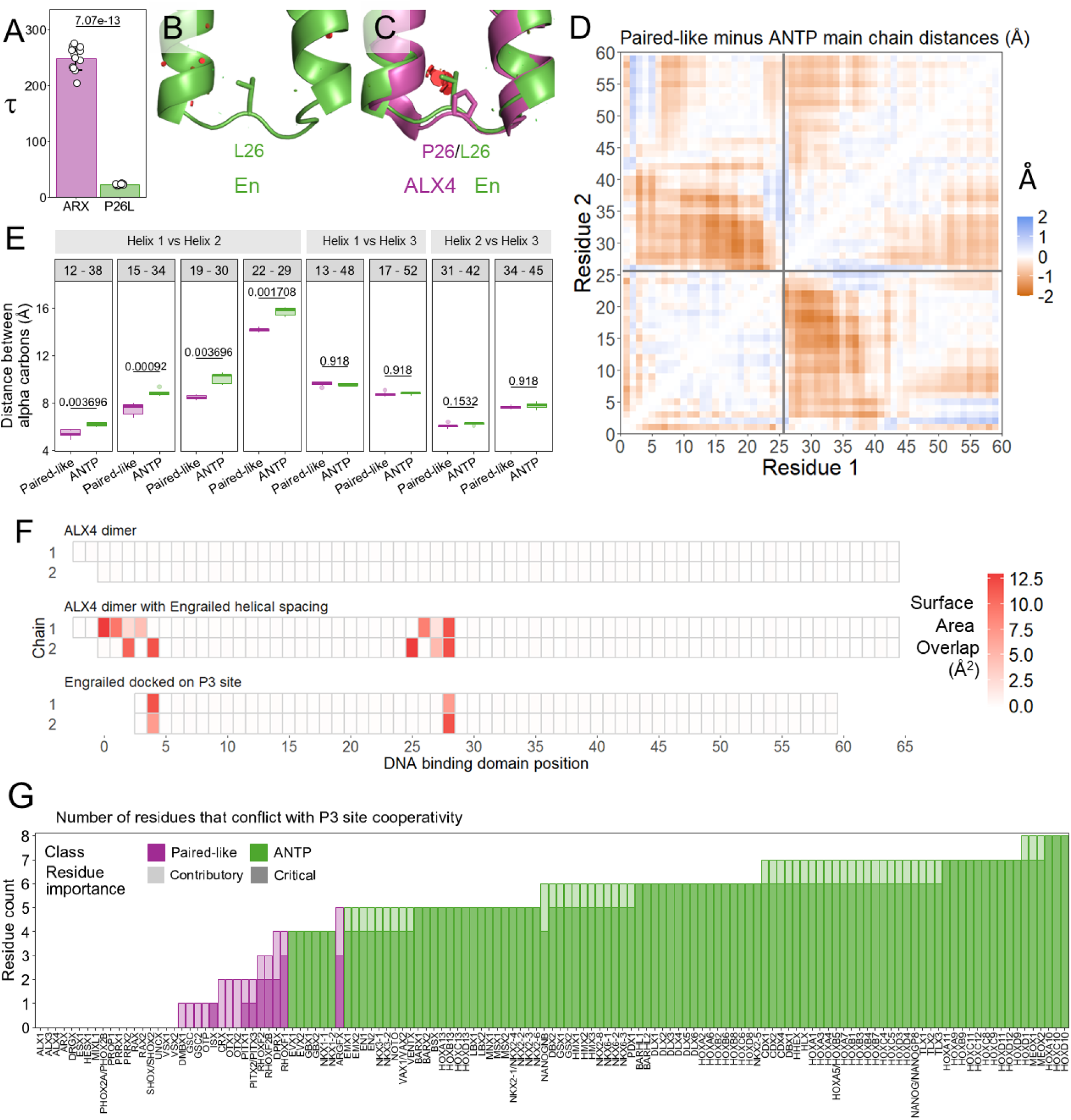
The P26L Paired-like to ANTP class swap found in the ARX disease variant decreases P3 site cooperativity. **(A)** ARX P26L causes an approximately 10-fold decrease in cooperativity. Tau cooperativity factors were calculated from triplicate EMSAs using every lane in which more than 5% of the probe was bound in quantitative EMSAs. Bars represent the average Tau for each protein, and each dot represents a Tau from an independent binding reaction. Error bars represent standard deviation. Tau was compared via two-sided Welch’s t-test. Gels are shown in Supplementary Figure 6. **(B)** PYMOL schematic of steric overlap of L26 in Engrailed (En) (1HDD). **(C)** PYMOL schematic of steric overlap of L26 in ALX4 chain A where L26 clashes with A30. ALX4 is shown in purple and Engrailed (En) is shown in green. **(D)** Average pairwise alpha carbon chain distances from solved structures reveals differences between 7 Paired-like proteins and 5 ANTP proteins. Heatmap colors correspond to average distance differences in angstroms. Pairwise distances for each protein are shown in Supplementary Figure 7. **(E)** Helix 1 and 2 are closer in proximity in Paired-like factors compared to ANTP factors. Lower and upper hinges denote the first and third quartiles, respectively. The lower and upper whiskers denote the most extreme value if there are no outliers or 1.5 times the interquartile range. Pairwise distances of inner helical pairs were compared with two-sided t-tests with a Holm multiple comparisons correction. Distances for each pairing in each structure are shown in Supplementary Figure 8. **(F)** Steric overlap of residue with non-bonded residues in opposing protein in dimer complex. **(G)** Number of residues that conflict with P3 site cooperativity based on determined rules (Table 1). Bar transparency denotes importance of the conflicting rule where higher transparency represents rules that contribute to the degree of cooperativity, and lower transparency represents rules that are critical to cooperativity. Conflicts for each factor are detailed in Supplementary Table 1.

To address how helical spacing influences cooperativity, we calculated the pairwise surface area overlaps between the two proteins of three distinct dimer complexes on the P3 site: (1) the wildtype ALX4 dimer; (2) the ALX4 dimer modeled with the wider helical spacing to accommodate the L26 residue found in the ANTP class; and (3) the Engrailed ANTP factor. These models revealed that the increased helical distance between helix 1 and 2 in the ANTP factors shifts the position of the turn between helix 1 and 2 and alters the energy favored rotamer conformations of residues in the turn between helix 1 and helix 2. Cumulatively, these changes cause a significant increase in surface area overlap at non-bonded residues at the protein-protein interfaces between the N-terminal ARM and turn between helix 1 and helix 2 (Figure 5F; Supplementary Figure 9), and these clashes would not be conducive to cooperative binding.

### Defining rules of HD cooperativity on P3 sites

Based on the comparative studies between cooperative and non-cooperative Paired-like HDs (Figure 2–Figure 3) and the ANTP class of HDs (Figure 4–Figure 5), we generated 11 rules that define cooperative binding to P3 sites (Table 1). Eight of these rules are necessary for P3 site cooperativity (denoted as critical), whereas three rules influence the degree of cooperativity, but are not required for cooperativity (denoted as contributory). Applying as few as two of the critical rules to the ANTP class of HDs reveals differences that are sufficient to explain why ANTP factors fail to cooperatively bind P3 sites. First, 99 of the 102 ANTP factors have a Q44, which drastically decreases cooperativity when inserted into a cooperative Paired-like HD (Figure 4F-G). Second, many ANTP HDs, including the 3 ANTP factors that have the required R44 (PDX1, EVX1, and EVX2), encode a positively charged residue at the 28^th^ position, whereas hydrophobic residues are preferred in the Paired-like class (Figure 4D). R28 is known to disrupt cooperativity on the P3 site (Cain et al., 2025; D. S. Wilson et al., 1995), likely due to the positively charged side chain clashing with the N-terminal ARM. Applying all 11 rules across the ANTP family revealed that it would take a minimum of four residue changes to make an ANTP factor cooperative on P3 sites (Figure 5G; Supplementary Table 1). We also applied these rules to all 36 Paired-like HDs and identified 16 factors that break one or more rule (Figure 5K). Seven of these Paired-like factors only conflict with contributory rules but follow all critical requirements (Figure 5G). These predictions are consistent with our findings that GSC, OTX2, and CRX exhibit relatively low levels of cooperativity as each break two contributory rules (Figure 1I; Figure 5G).

### Paired-like factor pathogenic variants impact P3 site cooperativity

We next applied these rules to disease variants across the Paired-like HD family to identify variants that impact cooperative DNA binding. Analysis of the HGMD (version 2025.2) identified 189 HD missense variants across the cooperative Paired-like factors that are associated with disease ((Stenson et al., 2017, 2020); Figure 6A). Collectively, 53 of the 189 identified disease variants impact a HD position that contributes to or is critical for cooperative DNA binding. However, it is important to note that some of the positions contain several different disease-associated residues with distinct biochemical properties (Figure 6A). For example, we found that the PHOX2B R44Q variant disrupts cooperativity (Figure 4G), whereas a PHOX2B R44G variant in the same HD position likely destabilizes the third helix and thereby decreases DNA binding affinity (Figure 4H). In addition to these PHOX2B disease variants, we already tested and confirmed that PHOX2B Q46R (Figure 4I), ARX P26L (Figure 5A) and ARX R44Q (Figure 4F) decrease cooperativity. Next, we apply these rules to predict disease variants that are likely to impact cooperativity versus DNA binding based upon the biochemical properties of each variant residue (Figure 6A). To validate these predictions, we selected 9 additional disease variants that occur at these positions for biochemical testing.

**Figure 6.**
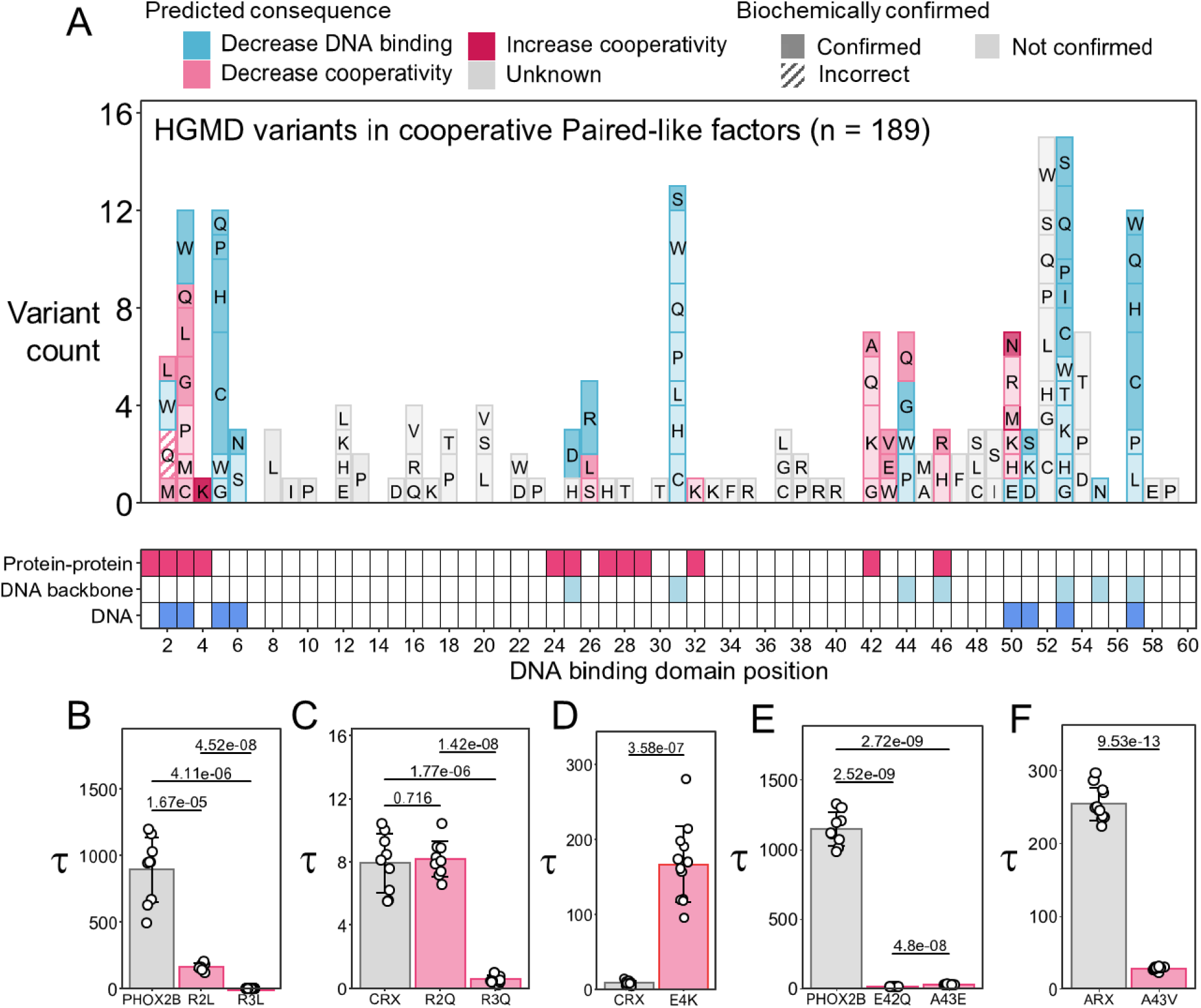
Pathogenic variants in Paired-like factors impact cooperative binding. **(A)** Pathogenic variants from HGMD (Stenson et al., 2017, 2020) were organized by DNA binding domain position. Biochemical consequence of each pathogenic variant was assessed based on compliance to the P3 site cooperativity rules (Table 1) as well as DNA binding domain position and altered residue property. Variants that are predicted to disrupt DNA binding are denoted in blue and variants predicted to impact cooperative binding are denoted in pink. Variant shading denotes whether the prediction has been confirmed biochemically. 14 variants were tested in biochemical assays in this study. 41 variants that are predicted to influence cooperativity or DNA binding were tested in prior studies (see Supplementary Table 2). Contact grid shows which DNA binding domain positions are involved in protein-protein interactions (pink), DNA backbone interactions (light blue), and DNA base interactions (dark blue). **(B-F)** Tau cooperativity factors were calculated for every lane in which more than 5% of the probe was bound in quantitative EMSAs. Bars represent the average Tau for each protein, and each dot represents a Tau from an independent binding reaction. Error bars represent standard deviation. **(B)** Tau was compared with a one-way ANOVA (3 proteins; n = 9; p = 1.81e-12). Pairwise two-sided Welch’s t-tests were performed with a Holm correction for multiple comparisons. **(C)** Tau was compared with a one-way ANOVA (3 proteins; n = 9; p = 1.37e-12). Pairwise two-sided Welch’s t-tests were performed with a Holm correction for multiple comparisons. **(D)** Tau was compared with a two-sided Welch’s t-test (n = 12). **(E)** Tau was compared with a one-way ANOVA (3 proteins; n = 9; p = 1.07e-22). Pairwise two-sided Welch’s t-tests were performed with a Holm correction for multiple comparisons. **(F)** Tau was compared with a two-sided Welch’s t-test (n = 12). Uncropped gels are shown in Supplementary Figure 10.

In total, 3 positions in the N-terminal ARM are critical for or contribute to cooperativity (Table 1). Of these positions, 6 missense variants in R2 (contributory) and 12 missense variants in R3 (critical) have been associated with disease states. In ALX4, we previously used alanine mutagenesis to show that having an arginine at both position 2 and 3 is required for cooperative P3 DNA binding due to their contributions to protein-protein interactions. In contrast, a single arginine at one of these positions was sufficient to maintain DNA binding affinity as either can insert into the minor groove and contact DNA. We also found that R3A disrupted cooperativity to a greater degree compared to R2A in ALX4 (Cain et al., 2025), consistent with R2 contributing to cooperative binding, whereas R3 is critical to cooperative binding (Table 1). While alanine was not found among the disease associated missense variants in Paired-like HDs at positions 2 and 3, leucine, which is larger, but biochemically similar as alanine, was associated with disease states in PHOX2B (R2L (Monies et al., 2017) and R3L (Trochet et al., 2004)) and ARX (R3L (Gong et al., 2022)) (Supplementary Table 2). Consistent with our prior ALX4 data, we found that both PHOX2B leucine variants decreased cooperativity with R3L doing so to a greater extent than R2L (PHOX2B wildtype: 894 ± 242; PHOX2B R2L: 167 ± 24.6; PHOX2B R3L: 5.40 ± 0.500; Figure 6B). Moreover, like ALX4 R2A and R3A, each of the PHOX2B variants caused only minor changes to overall DNA binding affinity as measured by free probe depletion (Supplementary Figure 10A).

We performed a similar analysis on variants in CRX, which has been reported to have R2Q, R3Q, and R3W variants associated with cone rod dystrophy (Carss et al., 2017; Swain et al., 1997). *In silico* mutagenesis predicts that CRX R3Q would fail to facilitate the salt bridge with E42 at the protein-protein interface (Supplementary Figure 11B) and suggests this residue would not maintain all hydrogen bonds in the minor groove of DNA (Supplementary Figure 11E). Because of this, we expect that a CRX R3Q variant would use its R2 residue to insert into and bond with the minor groove of DNA, resulting in a loss of the protein-protein interface between the N-terminal ARM and turn between helix 1 and helix 2. However, this residue is not expected to sterically clash at either interface (Supplementary Figure 11B, E). This contrasts with the CRX R3W variant which is predicted to strongly clash at the protein-DNA interface and decrease overall DNA binding (Supplementary Figure 11F). Similar to the R3Q variant, the CRX R2Q variant, much like leucine, glycine, or alanine at position 2, would fail to form the required minor groove interactions but would not sterically clash at either interface and disrupt DNA binding. Consistent with our *in silico* modeling, we found that CRX R3Q disrupts cooperativity (Figure 6C) but largely maintains DNA binding affinity (Supplementary Figure 10B), whereas CRX R3W strongly disrupts DNA binding affinity (Supplementary Figure 10G). Unexpectedly, however, we found that the CRX R2Q variant mildly decreased DNA binding affinity (Supplementary Figure 10B) rather than impacting cooperativity (Figure 6C). It is important to note that the ALX4 R2A and PHOX2B R2L variants decreased cooperativity to a tau of 20 (Cain et al., 2025) and 167 (Figure 6B) respectively, which are both greater than CRX wildtype’s tau. Because of this, it is possible that the R2Q variant would have had a greater impact on cooperativity in a protein with a higher wildtype cooperativity.

The 4^th^ residue within the N-terminal ARM also contributes to cooperative DNA binding. To date, there is only one disease associated variant in the 4^th^ position as CRX E4K is associated with Leber congenital amaurosis (Li et al., 2011). We previously showed that altering the CRX E4 residue into an asparagine (E4N) resulted in higher cooperativity than wildtype CRX (Figure 2F) as this mutation removes charge clashes between E4 and E32 and possibly facilitates polar bonds between E32 and N4, like those seen in the ALX4 dimer structure (Figure 2B). As a lysine is positively charged and not predicted to clash with the negatively charged glutamic acid, this disease variant may also increase cooperative binding (Table 1). Consistent with this prediction, CRX E4K bound with an 18-fold increase in cooperativity relative to wildtype CRX (CRX wildtype: 9.04 ± 2.79; CRX E4K: 167 ± 51.1; Figure 6D). The increased cooperativity may act similarly to a second Leber congenital amaurosis variant, CRX K50N (Rule 11), which was previously shown to increase cooperativity on the P3 site, and thereby cause ectopic binding that silenced progenitor chromatin states rather than directing differentiation of progenitor photoreceptors (Zheng et al., 2024).

Lastly, we found that HD positions 42, 43, 44, and 46 within the third helix are critical for cooperative DNA binding and position 50 is contributory. In total, 20 different Paired-like HD missense variants at these positions have been associated with disease. Of these, we tested the congenital central hypoventilation syndrome variants, PHOX2B E42Q (Zhou et al., 2021) and PHOX2B A43E (Parodi et al., 2008), and found both disrupt cooperativity with only minor changes in affinity (Figure 6E; Supplementary Figure 10D). PHOX2B E42Q would be predicted to disrupt the R3-E42 salt bridge (Cain et al., 2025; D. S. Wilson et al., 1995). This is consistent with previous work that demonstrated that ALX4 E42A fully disrupts cooperativity (Cain et al., 2025) as well as a genomics study that showed that CRX E42A decreased dimeric binding in *in vitro* DNA binding assays as well as in genomic assays in the mouse retina (Zheng et al., 2024). PHOX2B A43E most likely clashes at the interface between the two recognition helices, similarly to what was seen in the ALX4 A43V and ALX4 A43D variants (Cain et al., 2023). Moreover, we confirmed that the disease variant associated with autism spectrum disorder, ARX A43V (Thai et al., 2020), also caused a 7.5-fold decrease in cooperativity (Figure 6F).

In summary, we predicted that 38 human disease variants would alter cooperativity (Figure 6A; Supplementary Table 2). After biochemical testing, we found that 11 of 12 of the predicted cooperativity variants selectively disrupt cooperativity without large impacts to DNA binding affinity. These studies complement two recent studies that identified four human disease variants that alter cooperativity (Cain et al., 2025; Zheng et al., 2024). Collectively, these findings highlight how cooperativity can increase DNA binding specificity and that variants that disrupt this cooperative DNA binding behavior can sufficiently alter protein function to cause disease.

## 4. Discussion

The Paired-like HD genes constitute a large family of TFs that are critical for proper development of many tissues and organs. HD missense alleles have been associated with abnormal craniofacial, neurological, ophthalmological, and cardiac development (Stenson et al., 2017, 2020). Intriguingly, disease alleles in the same Paired-like gene can lead to a range of phenotypes, highlighting potential genotype/phenotype relationships that can alter disease penetrance and expressivity. These findings raise two questions: How do members of the Paired-like HDs gain target specificity to regulate diverse developmental processes, and how do missense alleles in these genes impact HD function to cause disease? Here, we found that the Paired-like HDs vary greatly in their ability to cooperatively bind the palindromic P3 DNA site. By combining sequence constraint analysis within the Paired-like subfamily and against the non-cooperative ANTP HDs, protein-protein contact information from structural studies, and quantitative DNA binding measurements, we identified 11 rules that are either critical for or contribute to the dramatic differences in cooperative DNA binding between members of the Paired-like HDs. We then applied these rules to: (1) Predict the cooperative behavior of the remaining Paired-like HDs; (2) Explain why members of the ANTP subfamily of HD factors fail to cooperatively bind the P3 site; (3) Assess the functional impacts of human disease variants across the Paired-like HDs. From the variant allele analysis, we identified 38 disease associated variants predicted to alter cooperative DNA binding and performed validation studies to show that 11 of the 12 tested variants selectively altered cooperativity without greatly impacting DNA binding affinity. Collectively, these studies have two important implications: First, they show how differences in HD residues at positions not normally associated with DNA binding specificity can contribute to cooperative DNA binding and thereby help explain the different binding behaviors of Paired-like factors. Second, they reveal how disease associated missense variants at key HD positions in Paired-like factors can be stratified into different groups based upon their impact on cooperative DNA binding versus overall DNA binding affinity.

### Sequence constraint as a method to identify HD residues that contribute to cooperative DNA binding and specificity

The way in which HD TFs gain sufficient specificity to bind distinct genomic targets using highly conserved DNA binding domains has been a long-standing question in the field. Here, we took advantage of the large number of HDs in humans to compare the evolutionary sequence constraint across the Paired-like class, a subfamily of HDs in which many, but not all, members can cooperatively bind the P3 site, and the ANTP class, a HD subfamily that does not cooperatively bind the P3 site. With this analysis, we found that select Paired-like HDs cooperatively bind the longer palindromic P3 site using additional residues that do not directly contact DNA but contribute to complex formation via direct protein-protein interactions or indirect mechanisms such as altered helical spacing. Importantly, many of these residues are largely specific to the Paired-like class and not conserved in the non-cooperative ANTP class (Figure 4D). Further, these residues are not conserved across all Paired-like class members and often vary in factors that are weakly or non-cooperative (Figure 2A). This contrasts with many of the residues involved in direct protein-DNA contacts, such as the R5 and N51 residues that make essential contacts to TAAT sequences within the minor and major groove, respectively, which are largely conserved across HDs. Thus, while Paired-like and ANTP class HDs bind highly similar TAAT monomer sites using the same residues at key DNA contacting positions, residue differences between Paired-like and ANTP factors at non-DNA contacting positions, especially within the turn between Helix 1 and 2 (positions 26, 28, and 32) and at the beginning of the 3^rd^ helix (positions 42, 43, 44, and 46) increase HD DNA binding specificity by contributing to and stabilizing the formation of cooperative dimer complexes on the P3 site.

Of the 11 rules found to contribute to Paired-like HD cooperativity on P3 sites, HD positions 2, 3, 28, 42, and 43 were initially described in the original structural studies on the ALX4 and Prd S50Q proteins (Cain et al., 2025; D. S. Wilson et al., 1995). Here, we validated those studies as well as further identified and elucidated the impacts of HD positions 4, 26, 32, 44, 46, and 50 on cooperative complex formation. Four of these residues (4, 32, 46, and 50) were identified by direct comparative analysis of cooperative versus non-cooperative Paired-like factors. For example, our data using ALX4 and CRX proteins that swap only the 50^th^ amino acid residue reveals that a Q50 residue is conducive to high cooperativity, and this cooperativity enhances DNA binding specificity by enabling proteins to bind HD sites with suboptimal flanking regions (Figure 3). However, the structural mechanism behind the decreased cooperativity seen in the K50 HDs is unclear, but we can offer two possibilities: (1) Q50 typically interacts with DNA through weaker van der waals interactions or water mediated interactions compared to K50, which typically forms hydrogen bonds with the guanines of the opposing strand on a TAAT**CC** monomer site (Tucker-Kellogg et al., 1997). Hence, it is possible that the stronger bonds with the central nucleotides of the spacer sequence between the HD sites cannot be displaced and thereby prevents a second K50 protein from binding to the site. (2) We have previously shown that the Gsx2 HD, which binds as a dimer on two TAAT sites spaced 7bps apart, has enhanced cooperativity on DNA with an AT-rich spacer versus a GC-rich spacer (Webb et al., 2024). As much of the spacer sequence is not directly bound by the Gsx2 protein, these data suggest that DNA flexibility and/or DNA shape can directly contribute to cooperativity. Thus, it is possible that DNA shape and flexibility contribute to differences in cooperativity based on spacer nucleotide differences. However, in the case of the Paired-like factors, the HD sites are much closer together and the “spacer” sequences make numerous direct interactions with the HD proteins. Hence, if a similar mechanism affects the K50 vs Q50 Paired-like factors, there may be tradeoffs between the best spacer sequences for optimal shape/flexibility for cooperativity and the best spacer sequences for specificity through interactions with the K50 vs Q50 residues.

By leveraging sequence constraint differences and residue preferences between the Paired-like and ANTP classes, we identified three additional residues (26, 44, and 46) that directly contribute to cooperative DNA binding. Of these residues, structural analysis revealed mechanistic underpinnings for the R44Q and Q46K Paired-like to ANTP-like swaps due to the disruption of direct or water-mediated protein-protein interactions. However, the reason behind the loss of cooperativity in the P26L Paired-like to ANTP-like swap was not immediately apparent in the ALX4 or the Prd S50Q dimer structures. An in-depth structural analysis revealed that helix 1 and helix 2 are closer in proximity in the Paired-like structures compared to the ANTP structures (Figure 5D-E). We further found that this change in spacing is predicted to cause significant clashing at the protein-protein interfaces between the N-terminal ARM and turn between helix 1 and helix 2 (Figure 5F). However, we can only speculate the cause of the distinct helical spacings between these families. It is possible that the proline in the Paired-like factors increases rigidity in the turn between helices, providing the required strict and tight spacing. However, it is also possible that other covarying residues contribute to the distinct helical spacing between helix 1 and helix 2. For example, a previous covariation analysis found that HDs containing a branched aliphatic residue at position 26 also formed a salt bridge between positions 19 and 30 in 92% of cases. In contrast, if a HD contains a proline at position 26, as found in all Paired-like HDs, positions 19 and 30 are always hydrophobic (Clarke, 1995; Torrado et al., 2009). It is important to note that in most HDs, there is a single salt bridge that connects each helical pairing with positions 19 and 30 between helix 1 and 2, positions 17 and 52 between helix 1 and 3, and positions 31 and 42 between helix 2 and 3 (Torrado et al., 2009). Of these, the salt bridge mediating residues for helix 1 and 3 as well as helix 2 and 3 are conserved between the Paired-like and ANTP factors (Figure 4B-D), which correlates well with the conserved helical spacing across these pairings (Figure 5E). However, the residues at positions 19 and 30 are distinct between the families (Figure 4B-D), which may correlate with the observed change in helical spacing. This strong covariant relationship could also be related to either the 19-30 hydrophobic bridge or L26 being required for proper hydrophobic core formation. In the ALX4 structure, V19 and A30 insert into the hydrophobic core at the connection between helix 1 and 2 in a similar position to L26 in the Engrailed structure (Supplementary Figure 6C-D). Future studies will be needed to investigate the influence of the 19-30 salt bridge versus hydrophobic bridge as well as increased rigidity possibly provided by P26 versus the L26 hydrophobic core contributions in the determination of the helical spacing, stability, and cooperativity.

### Elucidating the impact of Paired-like HD cooperativity on protein function and disease variant mechanisms

By applying our cooperativity rules to human variants found in the HGMD (Stenson et al., 2017, 2020), we identified 38 variants predicted to impact cooperativity, whereas an additional 75 variants would likely impact overall DNA binding affinity (Figure 6A). In some cases, cooperativity helps clarify potential genotype to phenotype relationships of human disease variants. For example, variants in ARX have been associated with epilepsy, lissencephaly, intellectual disability, corpus collosum abnormalities, and autism (Stenson et al., 2017, 2020). Interestingly, there appears to be a loose correlation between variant impact on protein function and disease severity. All four ARX variants that occur at position 5, a position known to be critical to DNA binding and to strongly disrupt DNA binding when altered in ARX (Barrera et al., 2016; Cho et al., 2012; Shoubridge et al., 2012), cause severe lissencephaly or corpus collosum abnormalities (Kato et al., 2004; Kitamura et al., 2002; Lei et al., 2022; Uyanik et al., 2003), whereas variants that are predicted to selectively disrupt cooperativity such as ARX R3L and ARX P26S as well as variants we confirmed disrupt cooperativity, ARX P26L (Figure 5A), ARX A43V (Figure 6F), and ARX R44Q (Figure 4F) are associated with epilepsy and intellectual disability (Gong et al., 2022; Papuc et al., 2019; Strømme et al., 2002; Thai et al., 2020). This correlation can be seen at single DNA binding domain positions as well. As an example, the above mentioned ARX P26L variant that disrupts cooperativity, but largely maintains DNA binding affinity, has been associated with epilepsy, whereas ARX P26R disrupts all DNA binding (Barrera et al., 2016; Shoubridge et al., 2012) and has been associated with more severe structural defects such as lissencephaly (Kato et al., 2004). Altogether, these studies suggest that a variant’s impact on protein function could be derived from disease associations in select cases where one factor has associations with disease states of varying severity.

A limitation of our study is that we tested protein fragments that only encode the DNA binding domain on only one or two specific P3 sites that were preferred by Paired-like Q50 and K50 proteins in past SELEX assays (Cain et al., 2023; Jolma et al., 2013; D. Wilson et al., 1993). However, previous work has shown that although the spacer sequence content alters the magnitude of cooperative binding, it does not change the ability of ALX1, ALX4, or ARX to bind cooperatively to the P3 site (Cain et al., 2023). Despite these findings, we cannot rule out the possibility that select proteins and variants may behave differently on distinct P3 sites. This study also fails to capture the behavior of the full-length protein in its genomic environment. However, several Paired-like factors have genomic binding datasets available and show evidence of *in vivo* binding to the P3 site. For example, ALX4 was found to bind the P3 site in human cranial neural crest cells (Cain et al., 2025; Kim et al., 2024); Phox2a enriched for P3 dimer sites in cortical motor neurons (Cain et al., 2023; Mazzoni et al., 2013); and PHOX2B predominantly binds to only P3 sites in KELLY, BE2C, and CLBGA neuroblastoma cell lines (Boeva et al., 2017; Cain et al., 2023; Durbin et al., 2018). Hence, these genomic studies provide evidence that our biochemistry results are representative of what occurs *in vitro*.

Perhaps the best understood Paired-like HD factor in terms of genomic binding and transcriptional function is the weakly cooperative CRX protein. CRX is essential for proper photoreceptor development within the eye and genetic variants in CRX have been associated with cone-rod dystrophy, retinitis pigmentosa, macular dystrophy, and Leber congenital amaurosis (Stenson et al., 2017, 2020). At the transcriptional level, a massive parallel reporter assay comparing monomer and P3 dimer sites showed that CRX binding to dimeric sites correlates with higher transcriptional activation levels (Hughes et al., 2018). Moreover, a past genomic study in the mouse retina revealed that CRX variants that either increase or decrease cooperativity to P3 sites can cause disease (Zheng et al., 2024). For example, a CRX K50N variant associated with Leber congenital amaurosis both increased CRX cooperativity and altered DNA binding specificity, much like we found using the CRX K50Q variant (Figure 3). Importantly, the enhanced cooperativity and specificity change caused by the CRX K50N led to ectopic binding at Q50 sites that are not normally bound by wildtype CRX, thereby compromising photoreceptor development (Zheng et al., 2023, 2024). In contrast, a CRX E42A variant associated with cone rod dystrophy was found to disrupt cooperativity and behaved similarly as the PHOX2B E42Q (Figure 6E) and ALX4 E42A variants (Cain et al., 2025). The cooperativity-deficient CRX E42A variant led to accelerated chromatin remodeling in early photoreceptor development due to an increase in K50 monomer site binding but loss of remodeling at the end of development due to a loss of K50 dimer site binding (Zheng et al., 2024). Taken together, these studies reveal the importance of accurate cooperativity levels for the proper function of Paired-like HDs and suggest that stratifying disease variants based upon molecular defect may reveal new insights into genotype/phenotype relationships associated with disease.

## 5. Methods

### Protein preparation

For each Paired-like protein tested in this study, a 66 amino acid region including the 60 amino acid HD as well as 5 amino acids N-terminal and 1 amino acid C-terminal of the HD was PCR amplified from cDNAs with Accuzyme DNA polymerase (Bioline) or KOD Xtreme Hot Start DNA polymerase (Novagen) following manufacturer’s protocols. Primers and template DNAs used to amplify each gene are detailed in Supplementary table 3. PCR products were cut with either NdeI/XhoI or BamHI/NotI (NEB) and ligated into pET14P (sequences are listed in Supplemental Information) with T4 ligase (NEB) following manufacturer’s protocol. All sequences were confirmed via Sanger sequencing. Constructs were transformed into BL21 or C41(DE3) *E.coli,* grown in LB media to an optical density between 0.6 and 0.8, and induced in 0.5 mM isopropyl β-D-1-thiogalactopyranoside (IPTG). Proteins were purified with Ni-NTA beads (Qiagen). Protein purity was confirmed via SDS-PAGE with GelCode blue staining (Thermo Scientific) and total protein concentrations were determined via Bradford assay (BioRad). If protein purities differed between wildtype and variant proteins, the variant protein concentration was normalized to wildtype based on the band intensity of the protein of interest.

### Electrophoretic mobility shift assays

Probes were prepared by annealing a IRDye 700 nM labeled 5’ - AGCCACGCCCGCACTA - 3’ oligo to an oligo containing the binding sites of interest and then filling in the DNA with Klenow (NEB). The sequences of the oligos containing the binding sites of interest are denoted in Supplemental Information. All EMSA binding reactions were incubated for 20 minutes at room temperature with 10 mM Tris, pH 7.5; 50 mM NaCl; 1 mM MgCl_2_; 4% glycerol; 0.5 mM DTT; 0.5 mM EDTA; 50 µg/ml poly(dI–dC); and 200 µg/ml of BSA, 34 nM of denoted fluorescent probe, and the protein of interest. Protein concentrations are listed in figure captions for each respective experiment. Binding reactions were run for 2 hours at 150 volts on a 4% polyacrylamide gel and then scanned on the Li-Cor Odyssey CLx scanner. Band intensity measurements were made in the Li-Cor Image Studio software. All uncropped and unprocessed gels are provided in the Supplementary Figures. Tau was calculated by the equation below where [P_2_D] represents dimer binding, [PD] represents monomer binding, and [D] represents unbound probe as previously described (D. Wilson et al., 1993). Tau cooperativity factors were calculated for every lane in which more than 5% of the probe was bound.

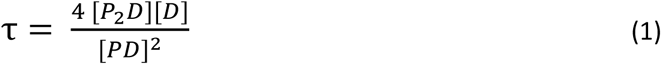

### Sequence constraint analysis

We identified 228 human HDs from CISBP (Weirauch et al., 2014), and found DNA binding domain annotation for 227 of these in UNIPROT (Bateman et al., 2025). HD classifications were performed as defined by Holland *et al* 2007 (Holland et al., 2007). We continued our analysis with the Paired-like and ANTP factors, which both lack additional DNA binding domains. For simplicity, we only continued with factors with exactly 60 amino acid DNA binding domains. We generated multiple sequence alignments of the 102 ANTP and 36 Paired-like DNA binding domains (Bodenhofer et al., 2015), converted these alignments to a matrix of Euclidian distances (Pelé et al., 2012), and then clustered with the complete linkage method and visualized with pheatmap. Bit information of DNA binding domain sequences was calculated and plotted with ggseqlogo (Wagih, 2017). Differential bit information was calculated by subtracting the bit information of the ANTP factors from the Paired-like factors at each residue and position.

### Helical spacing analysis

Alpha carbon distances were measured with the get_distance function in PYMOL (v2.5.7). Structures used in this analysis were published previously (Cain et al., 2025; Hovde et al., 2001; Joint Center for Structural Genomics (JCSG) & Partnership for Stem Cell Biology (STEMCELL), 2014; Kissinger et al., 1990; Morgunova et al., 2025; Srivastava et al., 2024; Webb et al., 2024; D. S. Wilson et al., 1995). The ALX4 structure was modeled with the Engrailed helical spacing by segmenting the structure into the N-terminal ARM (residues 1-8), helix 1 (residues 9-25), helix 2 (residues 26-39), and helix 3 (residues 40-60); aligning each helix to the like region of the Engrailed structure; creating bonds between the segments to generate a single object; and cleaning the newly bonded regions with the clean function. Steric overlap was calculated in PYMOL using the following approach: For each intermolecular pairwise combination, residue 1 and residue 2 were made into independent PYMOL objects and the surface area was calculated with the get_area command. An object was then created with both residues, and the combined surface area of both residues was calculated. The estimated overlap was derived by subtracting the combined surface area from the sum of the independent residue surface areas. Pairwise steric overlap of bonded residues was not reported.

## 6. Code availability

All custom scripts can be found at GitHub: https://github.com/cainbn97/PairedLikeCooperativity.

## 7. Acknowledgements

This work was supported by the National Institute of Health grants to B.G (R01-GM079428 and R35-GM158075) and a Center for Pediatric Genomics grant from CCHMC to B.G. The graphical abstract was created in BioRender (Cain, B. (2025) https://BioRender.com/5grruj2). We would like to thank Richard Mann, Shaoxun Liu, and Jeffrey Steimle for their helpful comments and suggestions on an earlier version of this manuscript.

## 8. Author contributions

The scientific study was conceived and planned by B.C. and B.G. Construct synthesis, protein purification, and EMSAs were performed by B.C., C.W., and F.R. All bioinformatic and structural analyses were performed by B.C. The manuscript was written by B.C. and B.G. and edited by all authors.

