## Supplementary Figures for "Decoding the mechanisms of cooperative DNA binding by the Paired-like homeodomain family"

### 1. Supplementary figure

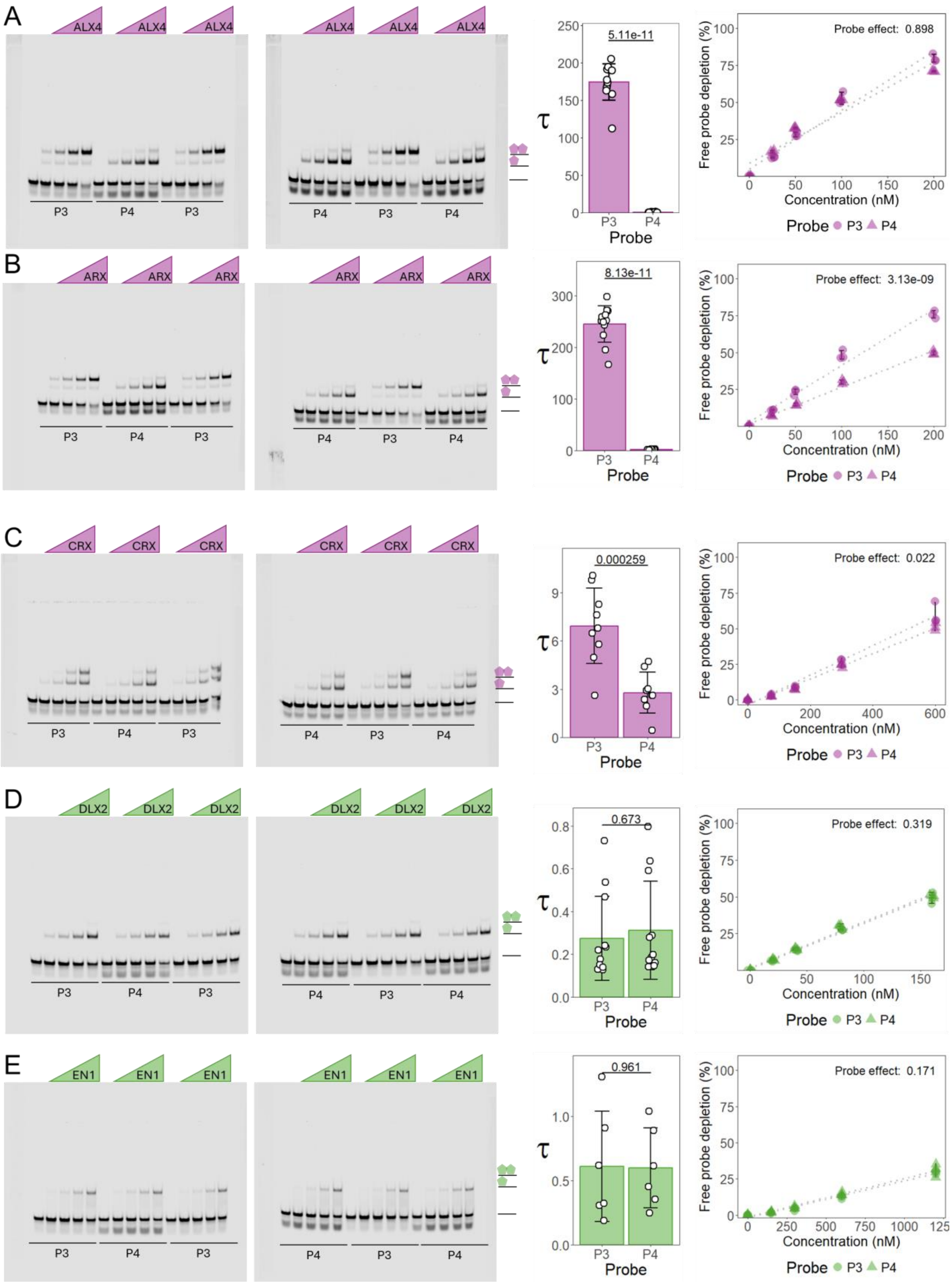

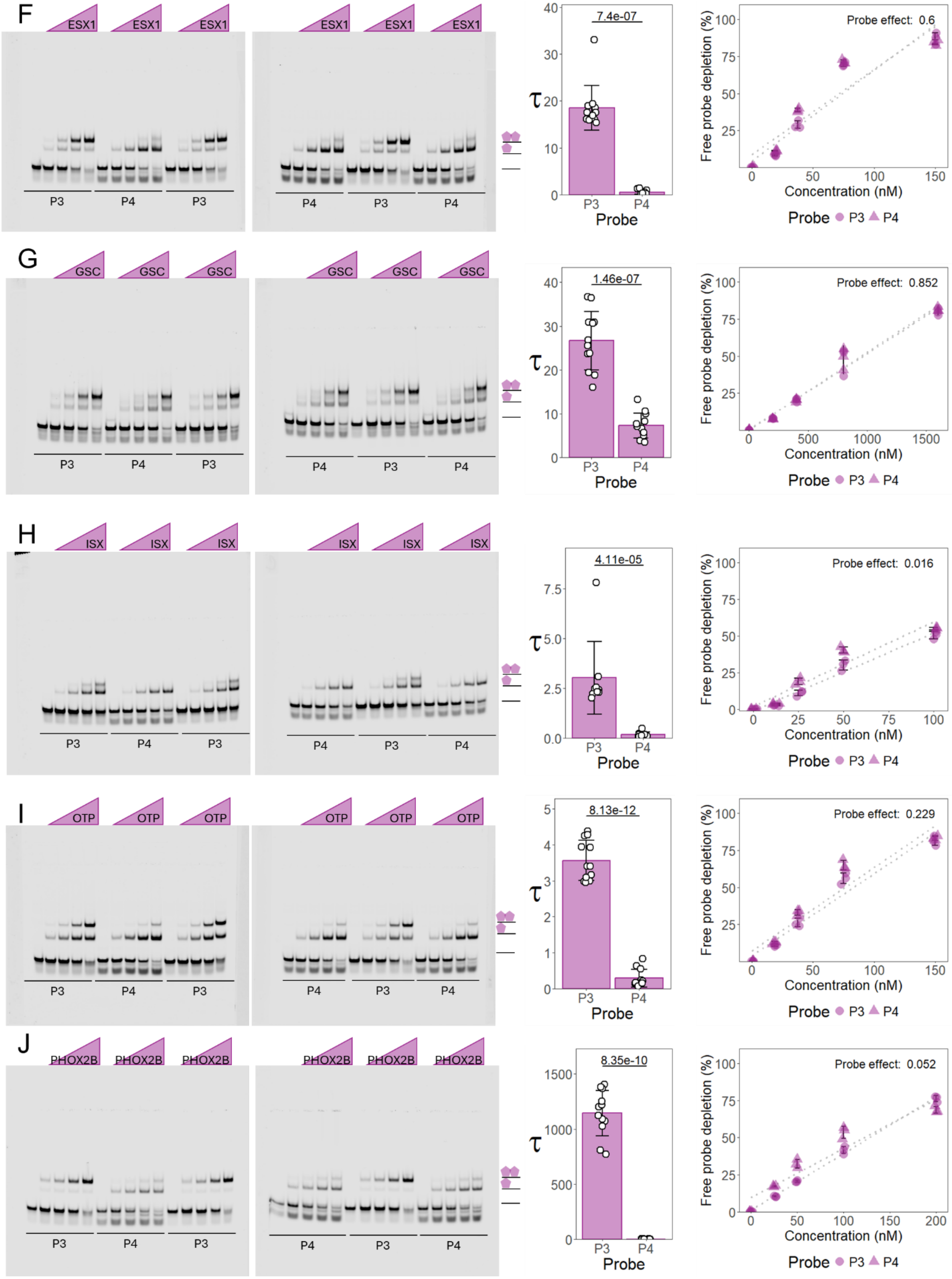
F
ESX1
P3
P4
7.4e-07
Probe effect: 0.6
G
GSC
1.46e-07
Probe effect: 0.852
H
ISX
4.11e-05
Probe effect: 0.016
I
OTP
8.13e-12
Probe effect: 0.229
J
PHOX2B
8.35e-10
Probe effect: 0.052
τ
Probe
Free probe depletion (%)
Concentration (nM)
Probe ● P3 ▲ P4

3

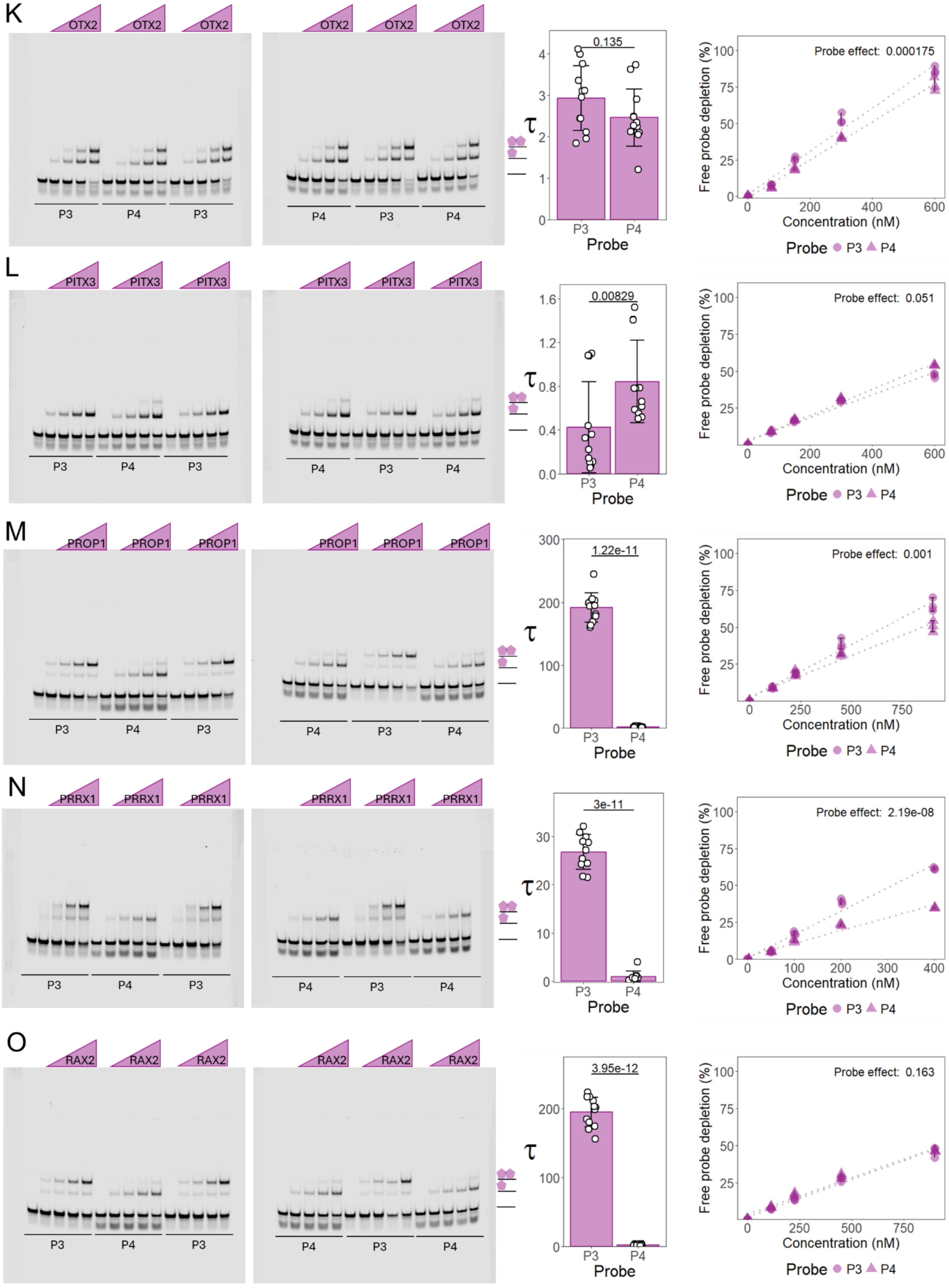
K
OTX2
P3
P4
τ
0.135
Probe
Free probe depletion (%)
Probe effect: 0.000175
Concentration (nM)
L
PITX3
0.00829
Probe effect: 0.051
M
PROP1
1.22e-11
Probe effect: 0.001
N
PRRX1
3e-11
Probe effect: 2.19e-08
O
RAX2
3.95e-12
Probe effect: 0.163

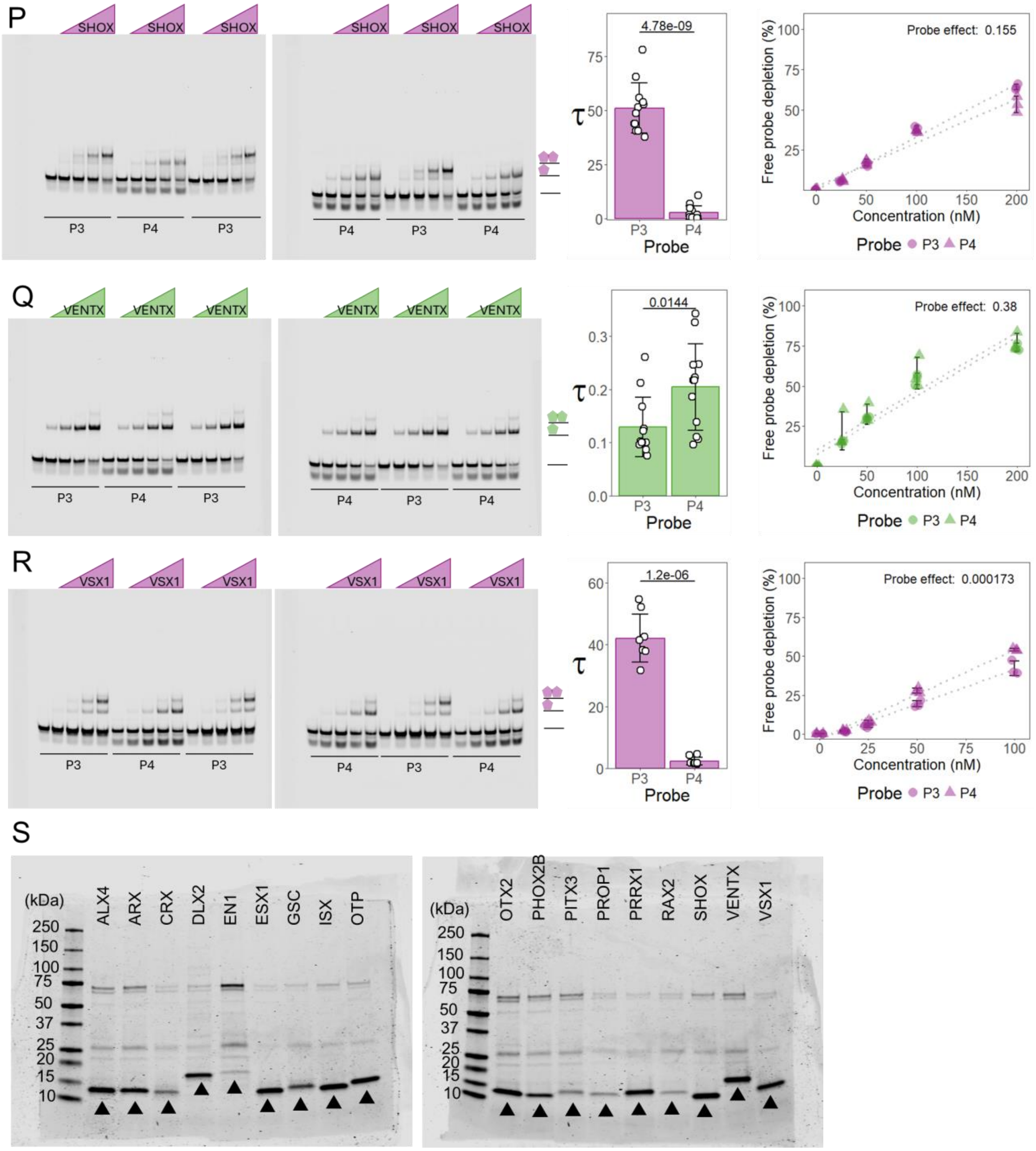


**Supplementary Figure 1.** All replicates for quantitative EMSAs used for Tau measurements in Figure 1I-J. Q50 HDs were tested on a P3 probe that contains a TAAT – TAG – ATTA sequence and a P4 probe that contains a TAAT – TAGG – ATTA sequence. K50 HDs (CRX, GSC, OTX2, and PITX3) were tested on a P3 probe that contains a TAAT – CCG – ATTA sequence and a P4 probe that contains a TAAT – CCGG – ATTA sequence.

Tau cooperativity factors were calculated for every lane in which more than 5% of the probe was bound. Bars represent the average Tau for each protein and the indicated probe with each dot representing a Tau from an independent binding reaction. Error bars represent standard deviation. The percentage of free probe depletion was calculated for each binding reaction. P3 and P4 free probe depletions for each independent binding reaction are denoted by circles and triangles, respectively. Free probe depletions were compared via a two-way ANOVA and the effect of the probe variable is reported (5 concentrations; n = 3). Paired-like factors are shown in purple and ANTP factors are shown in green. **(A)** ALX4 (amino acids = 209-274) was tested at 25, 50, 100, and 200 nM. Tau was compared via a Welch's two-sided t-test (n = 12). **(B)** ARX (amino acids = 323-388) was tested at 25, 50, 100, and 200 nM. Tau was compared via a two-sided Welch's t-test (n = 12) **(C)** CRX (amino acids = 34-99) was tested at 18.75, 37.5, 75, and 150 nM. Tau was compared via a two-sided t-test (n = 9). **(D)** DLX2 (amino acids = 137-229) was tested at 20, 40, 80, and 160 nM. Tau was compared via a two-sided t-test (n = 12). **(E)** EN1 (amino acids = 290-400) was tested at 150, 300, 600, and 1200 nM. Tau was compared via a two-sided t-test (n = 6). **(F)** ESX1 (amino acids = 134-199) was tested at 18.75, 37.5, 75, and 150 nM. Tau was compared via a two-sided Wilcox rank sum test (n = 12). **(G)** GSC (amino acids = 155-220) was tested at 200, 400, 800, and 1600 nM. Tau was compared via a two-sided Welch's t-test (n = 12). **(H)** ISX (amino acids = 77-142) was tested at 12.5, 25, 50, and 100 nM. Tau was compared via a two-sided Wilcox rank sum test (n = 9). **(I)** OTP (amino acids = 99-164) was tested at 18.75, 37.5, 75, and 150 nM. Tau was compared via a two-sided Welch's t-test (n = 12). **(J)** PHOX2B (amino acids = 93-158) was tested at 25, 50, 100, and 200 nM. Tau was compared via a two-sided Welch's t-test (n = 12). **(K)** OTX2 (amino acids = 33-98) was tested at 75, 150, 300, and 600 nM. Tau was compared via a two-sided t-test (n = 12). **(L)** PITX3 (amino acids = 57-122) was tested at 250, 500, 1000, and 2000 nM. Tau was compared via a two-sided Wilcox rank sum test (n = 12). **(M)** PROP1 (amino acids = 64-129) was tested at 112.5, 225, 450, and 900 nM. Tau was compared via a two-sided Welch's t-test (n = 12). **(N)** PRRX1 (amino acids = 89-154) was tested at 50, 100, 200, and 400 nM. Tau was compared via a two-sided Welch's t-test (P3 site n = 11; P4 site n = 10). **(O)** RAX2 (amino acids = 22-87) was tested at 112.5, 225, 450, and 900 nM. Tau was compared via a two-sided Welch's t-test (n = 12). **(P)** SHOX (amino acids = 113-177) was tested at 25, 50, 100, and 200 nM. Tau was compared via a two-sided Welch's t-test (n = 12). **(Q)** VENTX (amino acids = 69-178) was tested at 25, 50, 100, and 200 nM. Tau was compared via a two-sided t-test (n = 12). **(R)** VSX1 (amino acids = 159-224) was tested at 12.5, 25, 50, and 100 nM. Tau was compared via a two-sided Welch's t-test (P3 site n = 8; P4 site n = 9). **(S)** Protein purity was assessed with an SDS-PAGE. 4 µM of protein was loaded into each well. Band of interest is denoted by black triangle.

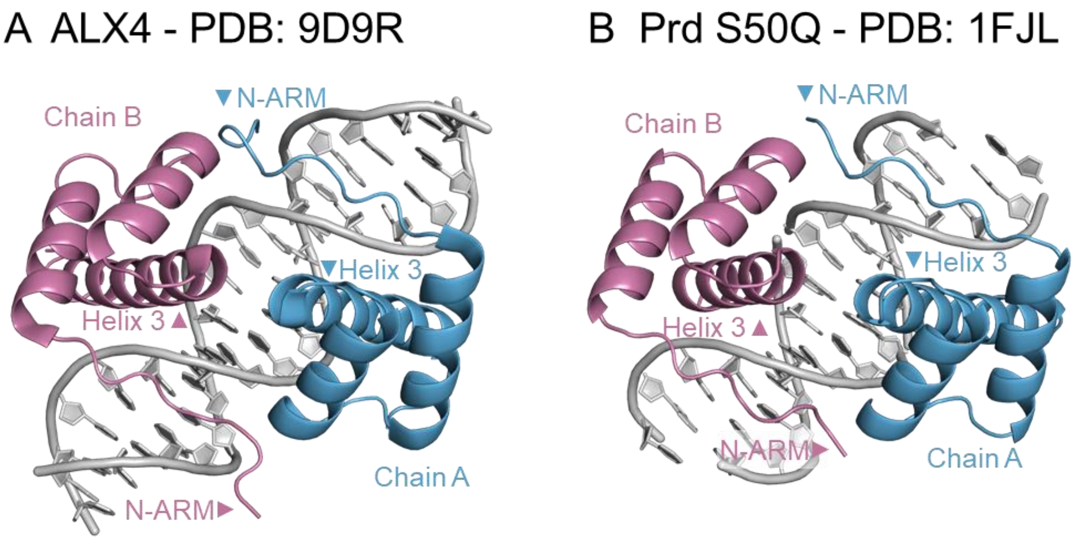


**Supplementary Figure 2.** ALX4 (A) and Prd S50Q (B) bind the P3 site in a head-to-head orientation. Chain A is shown in blue whereas Chain B is shown in purple.

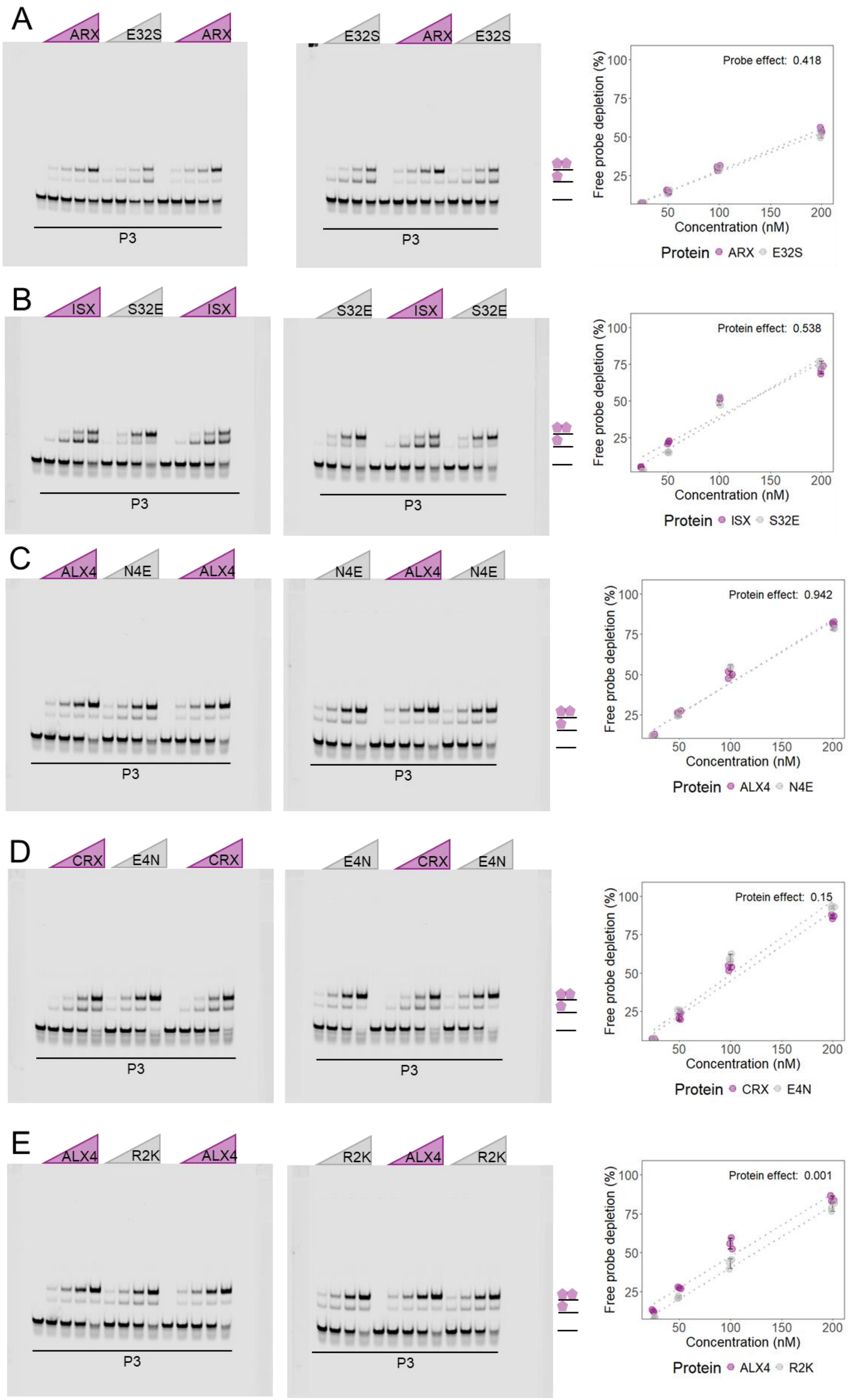
A
ARX
E32S
ARX
E32S
ARX
E32S
P3
P3
Probe effect: 0.418
Free probe depletion (%)
Concentration (nM)
Protein ARX E32S
B
ISX
S32E
ISX
S32E
ISX
S32E
P3
P3
Protein effect: 0.538
Free probe depletion (%)
Concentration (nM)
Protein ISX S32E
C
ALX4
N4E
ALX4
N4E
ALX4
N4E
P3
P3
Protein effect: 0.942
Free probe depletion (%)
Concentration (nM)
Protein ALX4 N4E
D
CRX
E4N
CRX
E4N
CRX
E4N
P3
P3
Protein effect: 0.15
Free probe depletion (%)
Concentration (nM)
Protein CRX E4N
E
ALX4
R2K
ALX4
R2K
ALX4
R2K
P3
P3
Protein effect: 0.001
Free probe depletion (%)
Concentration (nM)
Protein ALX4 R2K

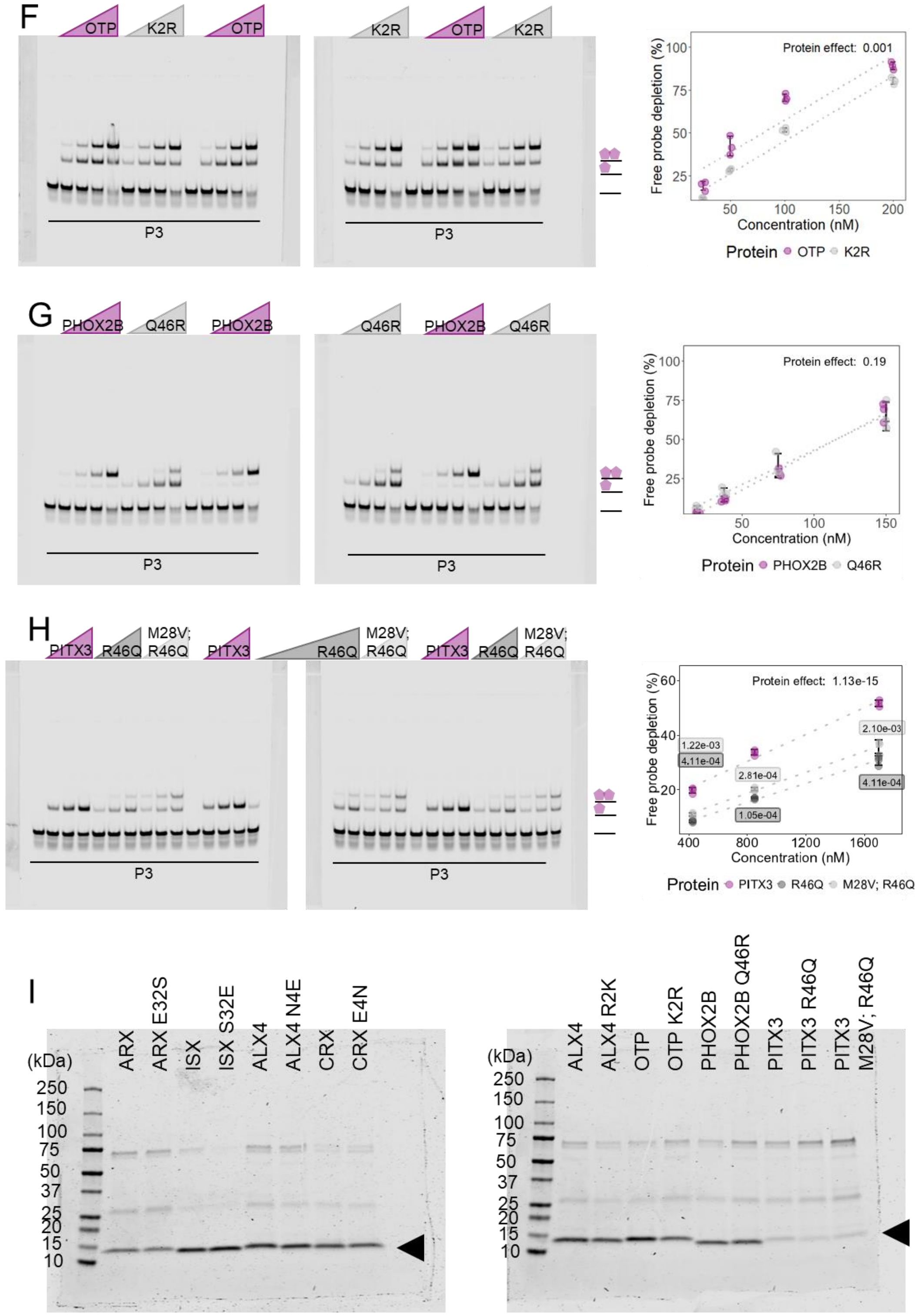
F
OTP
K2R
OTP
K2R
OTP
K2R
P3
P3
Protein effect: 0.001
Free probe depletion (%)
Concentration (nM)
Protein OTP K2R
G
PHOX2B
Q46R
PHOX2B
Q46R
PHOX2B
Q46R
P3
P3
Protein effect: 0.19
Free probe depletion (%)
Concentration (nM)
Protein PHOX2B Q46R
H
PITX3
R46Q
M28V; R46Q
PITX3
R46Q
M28V; R46Q
PITX3
R46Q
M28V; R46Q
P3
P3
Protein effect: 1.13e-15
1.22e-03
4.11e-04
2.81e-04
1.05e-04
2.10e-03
4.11e-04
Free probe depletion (%)
Concentration (nM)
Protein PITX3 R46Q M28V; R46Q
I
(kDa)
250
150
100
75
50
37
25
20
15
10
ARX
ARX E32S
ISX
ISX S32E
ALX4
ALX4 N4E
CRX
CRX E4N
ALX4
ALX4 R2K
OTP
OTP K2R
PHOX2B
PHOX2B Q46R
PITX3
PITX3 R46Q
PITX3 M28V; R46Q

**Supplementary Figure 3.** All replicates for quantitative EMSAs used for Tau measurements in Figure. Q50 HDs were tested on a P3 probe that contains a TAAT – TAG – ATTA sequence and a P4 probe that contains a TAAT – TAGG – ATTA sequence. K50 HDs (CRX and PITX3) were tested on a P3 probe that contains a TAAT – CCG – ATTA sequence. The percentage of free probe depletion was calculated for each binding reaction. Free probe depletions were compared via a two-way ANOVA and the effect of the protein variable is reported (4 concentrations; n = 3). **(A)** ALX4 wildtype and R2K (amino acids = 209-274) were tested at 25, 50, 100, and 200 nM. Tau was compared with a two-sided Welch's t-test. **(B)** OTP wildtype and OTP K2R (amino acids = 99-164) were tested at 25, 50, 100, and 200 nM. Tau was compared via a two-sided t-test. **(C)** ALX4 wildtype and N4E (amino acids = 209-274) were tested at 25, 50, 100, and 200 nM. Tau was compared via a two-sided Welch's t-test. **(D)** CRX wildtype and E4N (amino acids = 34-99) were tested at 25, 50, 100, and 200 nM. Tau was compared via a two-sided Welch's t-test. **(E)** ARX wildtype and E32S (amino acids = 323-388) were tested at 25, 50, 100, and 200 nM. Tau was compared via a two-sided Welch's t-test. **(F)** ISX wildtype and S32E (amino acids = 77-142) were tested at 25, 50, 100, and 200 nM. Tau was compared via a two-sided Welch's t-test. **(G)** PHOX2B wildtype and Q46R (amino acids = 93-158) were tested at 18.75, 37.5, 75, and 150 nM. Tau was compared via a two-sided Welch's t-test. **(H)** PITX3 wildtype, R46Q, and M28V;R46Q (amino acids = 57-122) were tested at 425, 850, and 1700 nM. Free probe depletions were compared via a two-way ANOVA and the effect of the protein variable is reported (3 concentrations; n = 3) with a Holm post-hoc to determine pairwise comparisons. Only comparisons to the wildtype protein are shown. **(I)** Protein purity was assessed with an SDS-PAGE. 3.5 µM of protein was loaded into each well. Band of interest is denoted by black triangle.

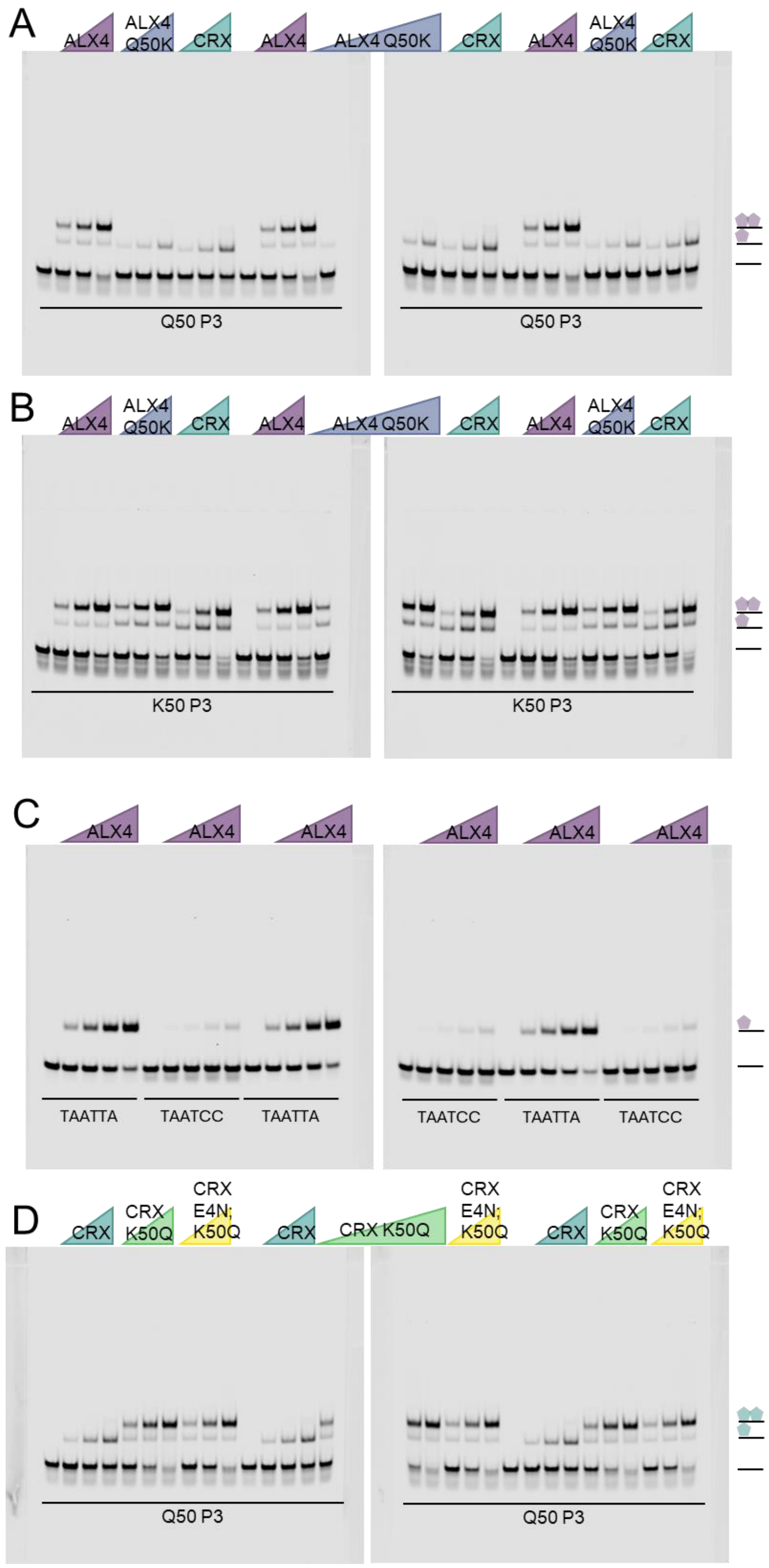

A
ALX4
ALX4 Q50K
CRX
ALX4
ALX4 Q50K
CRX
ALX4
ALX4 Q50K
CRX
Q50 P3
Q50 P3
B
ALX4
ALX4 Q50K
CRX
ALX4
ALX4 Q50K
CRX
ALX4
ALX4 Q50K
CRX
K50 P3
K50 P3
C
ALX4
ALX4
ALX4
ALX4
ALX4
ALX4
TAATTA
TAATCC
TAATTA
TAATCC
TAATTA
TAATCC
D
CRX
CRX K50Q
CRX E4N; K50Q
CRX
CRX K50Q
CRX E4N; K50Q
CRX
CRX K50Q
CRX E4N; K50Q
Q50 P3
Q50 P3


69

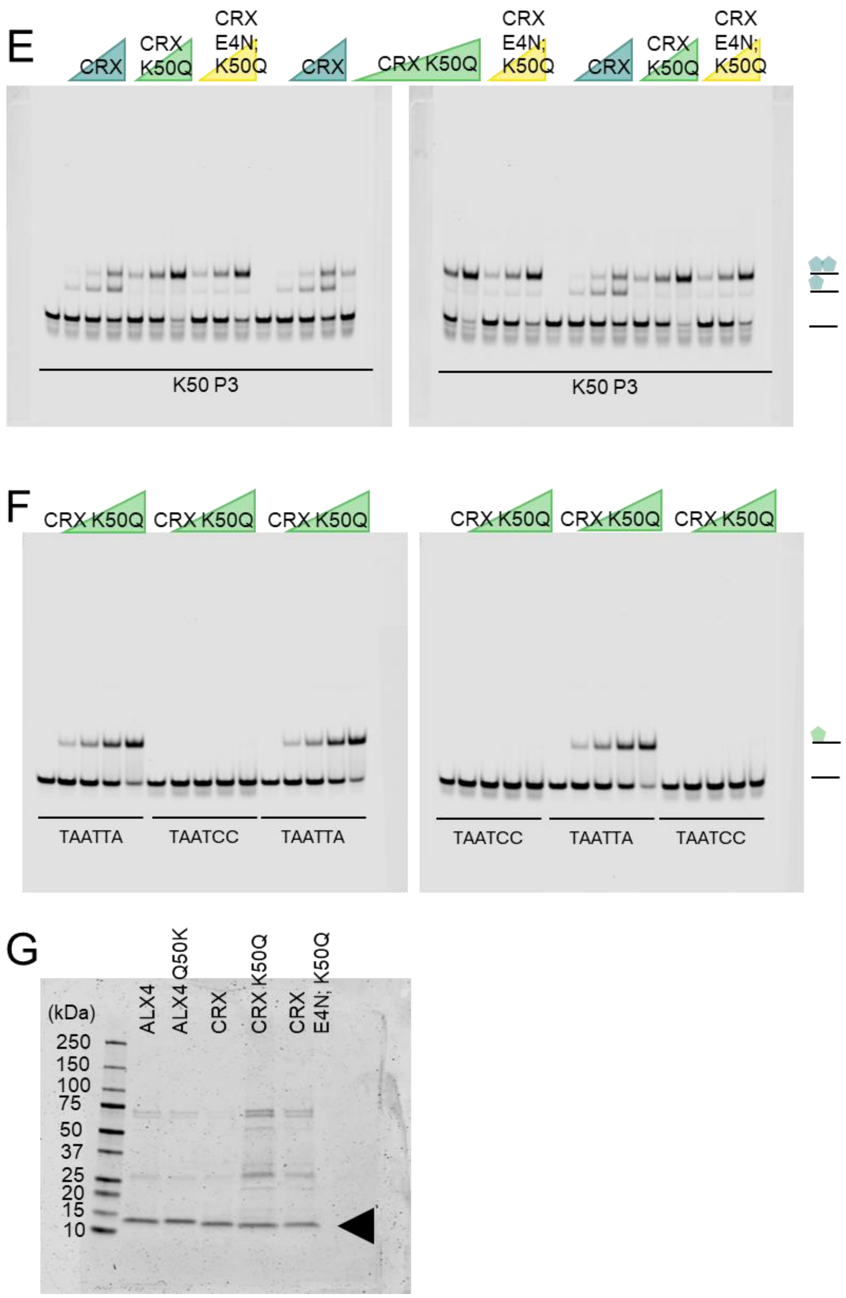


**Supplementary Figure 4.** Uncropped gels quantified in Figure 3. **(A-B)** ALX4 wildtype and Q50K (amino acids = 209-274) and CRX wildtype (amino acids = 34-99) were tested at 50, 100, and 200 nM on a Q50 P3 site (TAAT – TAG – ATTA) (A) and K50 P3 site (TAAT – CCG – ATTA) (B). **(C)** ALX4 wildtype (amino acids = 209-274) were tested at 25, 50, 100, and 200 nM on a Q50 monomer site (TAATTA) and a K50 monomer site (TAATCC). **(D-E)** CRX wildtype, CRX K50Q, and CRX E4N; K50Q (amino acids = 34-99) were tested at 50, 100, and 200 nM on a Q50 P3 site (TAAT – TAG – ATTA) (D) and K50 P3 site (TAAT – CCG – ATTA) (E). **(F)** CRX K50Q (amino acids = 34-99) was tested at 16.6, 33.2, 66.3, and 132.6 nM on a Q50 monomer site (TAATTA) and a K50 monomer site (TAATCC). **(G)** Protein purity was assessed with an SDS-PAGE. 2 μM of protein was loaded into each well. Band of interest is denoted by black triangle.

81

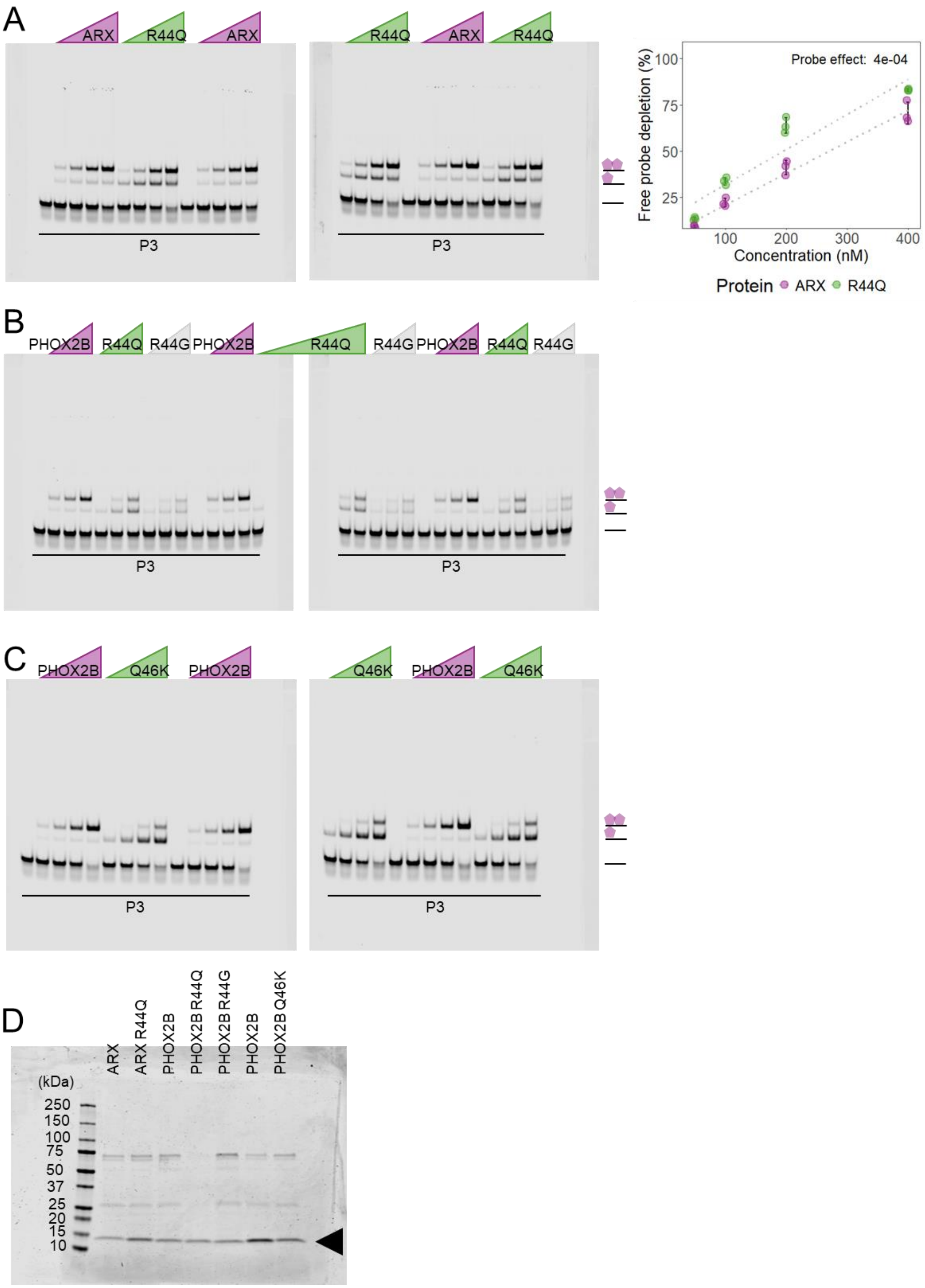
A
ARX
R44Q
ARX
R44Q
ARX
R44Q
P3
P3
Probe effect: 4e-04
Free probe depletion (%)
100
75
50
25
100
200
300
400
Concentration (nM)
Protein ARX R44Q
B
PHOX2B
R44Q
R44G
PHOX2B
R44Q
R44G
PHOX2B
R44Q
R44G
P3
P3
C
PHOX2B
Q46K
PHOX2B
Q46K
PHOX2B
Q46K
P3
P3
D
ARX
ARX R44Q
PHOX2B
PHOX2B R44Q
PHOX2B R44G
PHOX2B
PHOX2B Q46K
(kDa)
250
150
100
75
50
37
25
20
15
10

**Supplementary Figure 5. (A-D)** Uncropped gels from Figure 4F-J. **(A)** ARX wildtype and R44Q (amino acids = 323-388) were tested at 50, 100, 200, and 400 nM. Free probe depletion was calculated for each binding reaction. Free probe depletions were compared via a two-way ANOVA and the effect of the protein variable is reported (4 concentrations; n = 3). **(B)** PHOX2B wildtype, R44Q, and R44G (amino acids = 93-158) were tested at 23, 46, and 92 nM. **(C)** PHOX2B wildtype and Q46K (amino acids = 93-158) were tested at 18.75, 37.5, 75, and 150 nM. **(D)** Protein purity was assessed with an SDS-PAGE. 3 µM of protein was loaded into each well. Band of interest is denoted by black triangle.

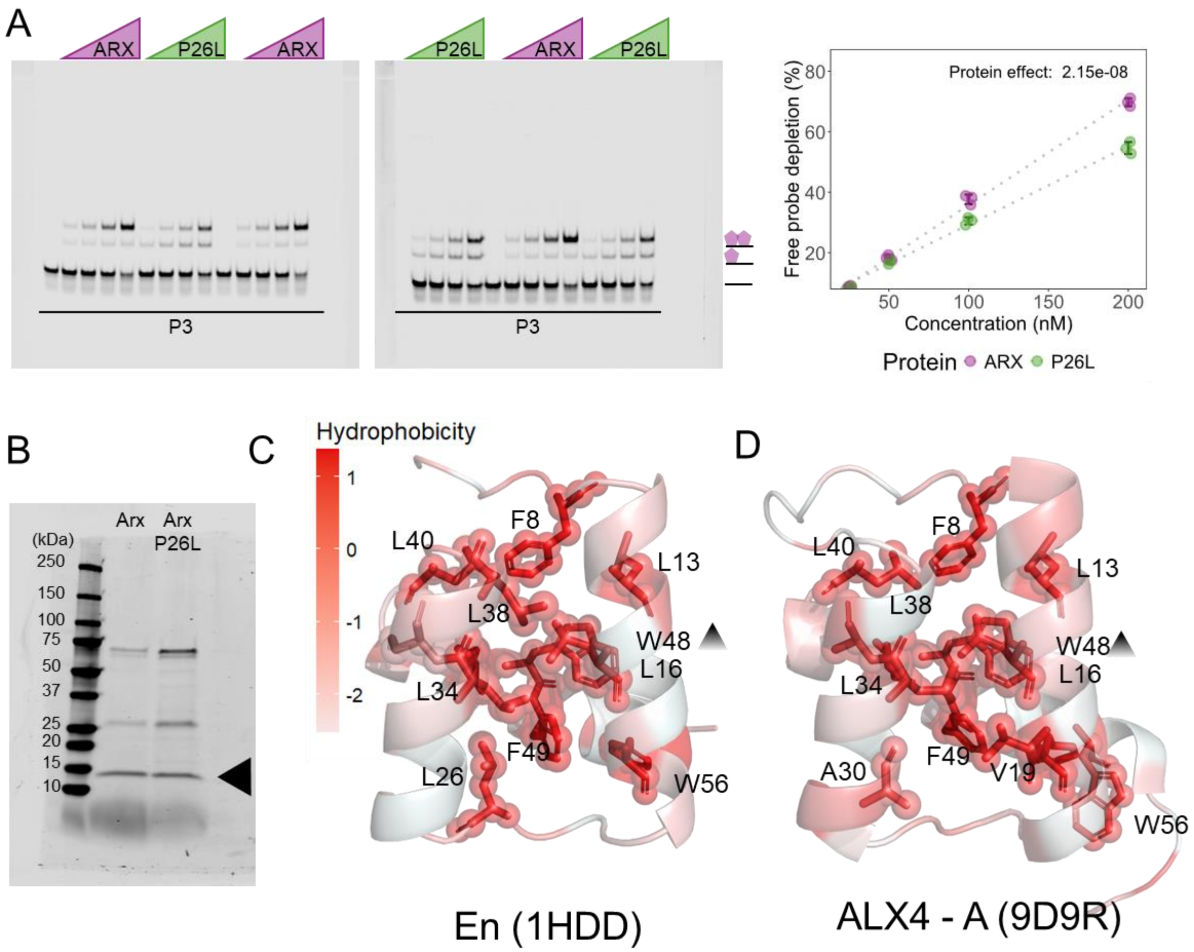


**Supplementary Figure 6. (A)** Uncropped gels from Figure 5A. ARX wildtype and P26L (amino acids = 323-388) were tested at 25, 50, 100, and 200 nM. Free probe depletion was calculated for each binding reaction. Free probe depletions were compared via a two-way ANOVA and the effect of the protein variable is reported (4 concentrations; n = 3). **(B)** Protein purity was assessed with an SDS-PAGE. 2.5 µM of protein was loaded into each well. Band of interest is denoted by black triangle. **(C-D)** Residues contributing to the hydrophobic core in Engrailed (En) and ALX4. Residues are colored based on residue hydrophobicity (Eisenberg et al., 1984). Residues contributing to the hydrophobic core are labeled and shown in stick and sphere formation. Triangle highlights that the indicated residue is behind another residue.

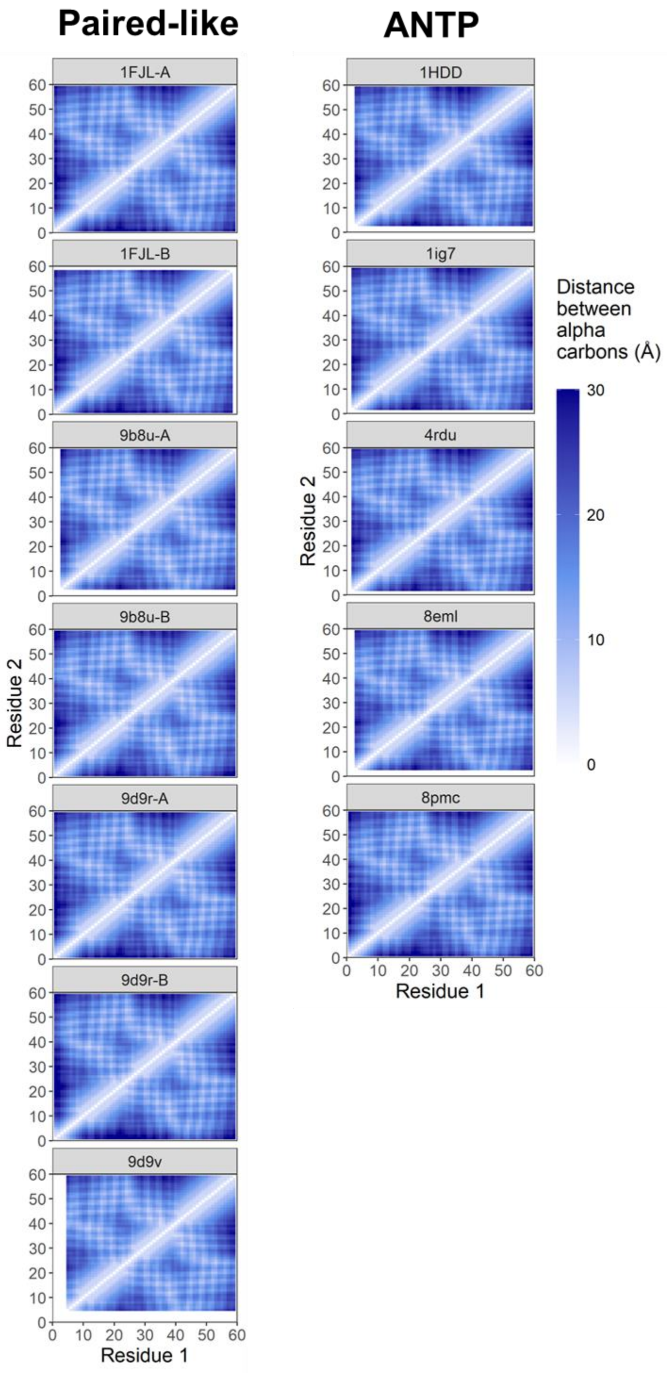


**Supplementary Figure 7.** Pairwise main chain distances for 7 Paired-like factors (Cain et al., 2025; Joint Center for Structural Genomics (JCSG) & Partnership for Stem Cell Biology (STEMCELL), 2014; Srivastava et al., 2024; Wilson et al., 1995) and 5 ANTP factors (Hovde et al., 2001; Kissinger et al., 1990; Morgunova et al., 2025; Webb et al., 2024). Heatmaps are labeled by PDB ID and organized by subclass.

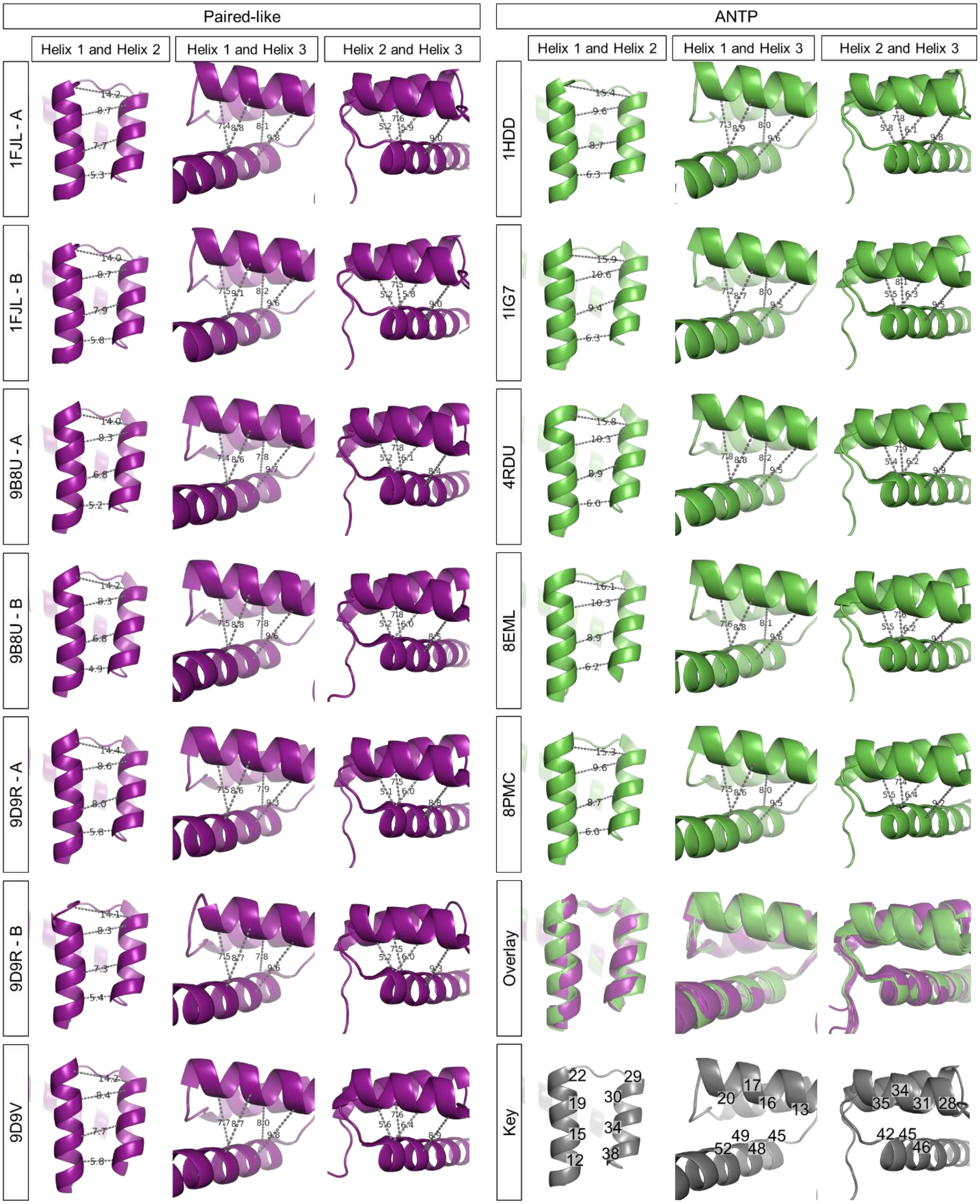


**Supplementary Figure 8.** Inner helical pair distance measurements and views for each structure used in the analysis. Overlay of all Paired-like (purple) and ANTP (green) factors. HD residue positions are labeled in Key.

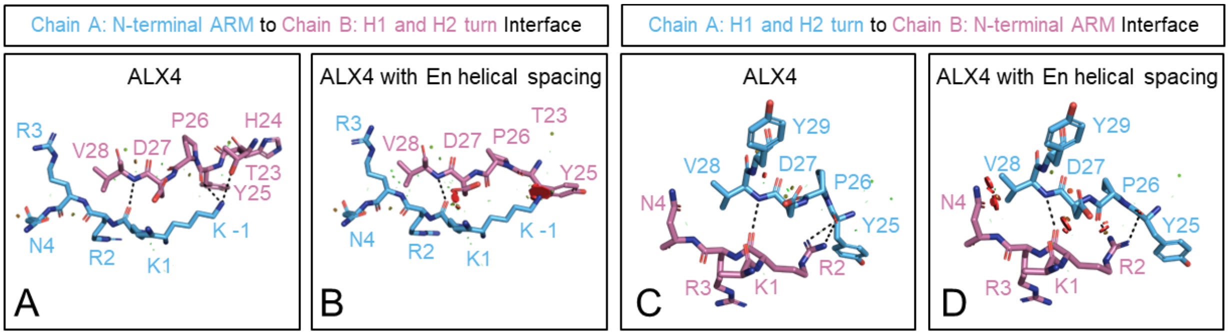


**Supplementary Figure 9.** PYMOL schematics in which Chain A is shown in blue and Chain B is shown in purple. Steric overlap between Chains is shown as discs, where the size and color of the disc correlates with amount of steric overlap. **(A)** PYMOL schematic of the interface between the N-terminal ARM of Chain A and the turn between helix 1 and 2 of Chain B in wildtype ALX4. **(B)** *In silico* prediction of the interface between the N-terminal ARM of Chain A and the turn between helix 1 and 2 of Chain B in an ALX4 protein modeled with En helical spacing. **(C)** PYMOL schematic of the interface between the turn between helix 1 and 2 of Chain A and the N-terminal ARM of Chain B in wildtype ALX4. **(D)** *In silico* prediction of the interface between the turn between helix 1 and 2 of Chain A and the N-terminal ARM of Chain B in an ALX4 protein modeled with En helical spacing.

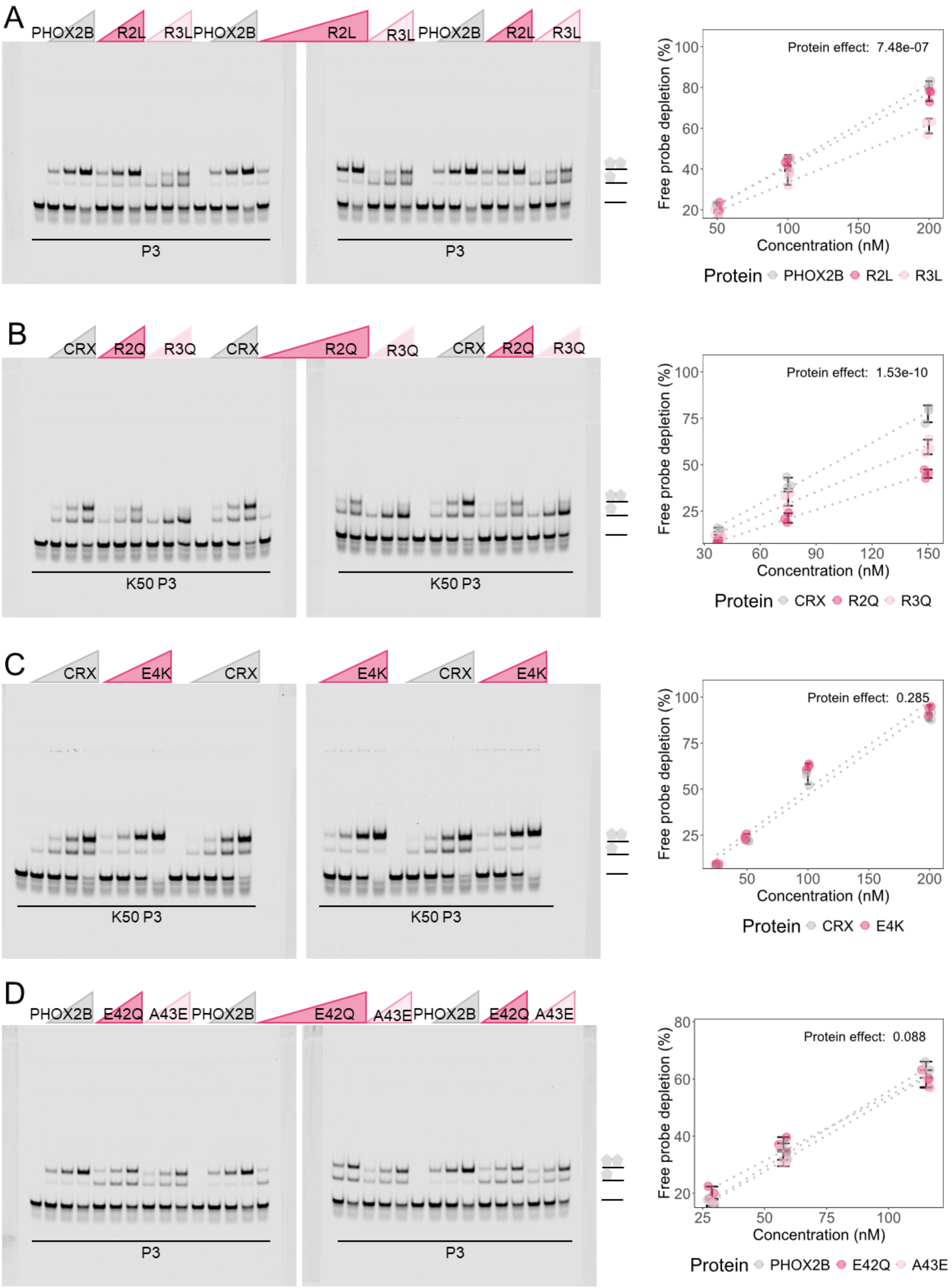

A
PHOX2B
R2L
R3L
PHOX2B
R2L
R3L
PHOX2B
R2L
R3L
P3
P3
Protein effect: 7.48e-07
Free probe depletion (%)
Concentration (nM)
Protein PHOX2B R2L R3L
B
CRX
R2Q
R3Q
CRX
R2Q
R3Q
CRX
R2Q
R3Q
K50 P3
K50 P3
Protein effect: 1.53e-10
Free probe depletion (%)
Concentration (nM)
Protein CRX R2Q R3Q
C
CRX
E4K
CRX
E4K
CRX
E4K
K50 P3
K50 P3
Protein effect: 0.285
Free probe depletion (%)
Concentration (nM)
Protein CRX E4K
D
PHOX2B
E42Q
A43E
PHOX2B
E42Q
A43E
PHOX2B
E42Q
A43E
P3
P3
Protein effect: 0.088
Free probe depletion (%)
Concentration (nM)
Protein PHOX2B E42Q A43E

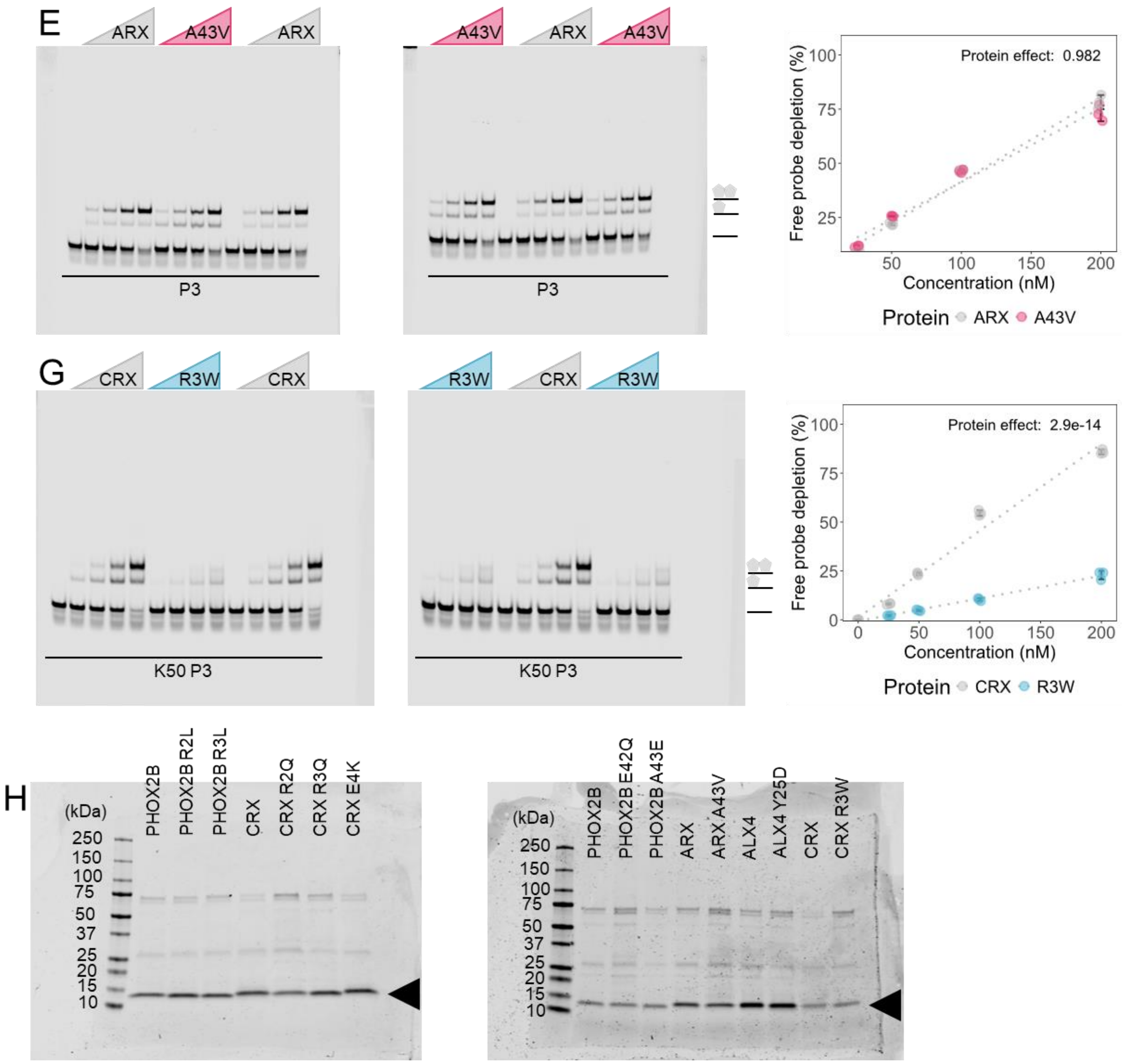


**Supplementary Figure 10.** Uncropped gels quantified in Figure 6B-E. **(A-F)** Free probe depletion was calculated for each binding reaction and is denoted by a circle. Free probe depletions were compared via a two-way ANOVA and the effect of the protein variable is reported. The number of concentrations and replicates used in the ANOVA are reported in each subpanel caption. **(A)** PHOX2B wildtype, R2L, and R3L (amino acids = 93-158) were tested on 50, 100, and 200 nM (3 concentrations; n = 3). **(B)** CRX wildtype, R2Q, and R3Q (amino acids = 34-99) were tested at 37.5, 75, and 150 nM (3 concentrations; n = 3). **(C)** CRX wildtype and E4K were tested at 25, 50, 100, and 200 nM (4 concentrations; n = 3). **(D)** PHOX2B wildtype, E42Q, and A43E (amino acids = 93-158) were tested at 28.8, 57.5, and 115 nM (3 concentrations; n = 3). **(E)** ARX wildtype and A43V (amino acids = 323-388) were tested at 25, 50, 100, and 200 nM (4 concentrations; n = 3) **(G)** CRX wildtype and R3W (amino acids = 34-99) were tested at 25, 50, 100, and 200 nM (4 concentrations; n = 3). **(H)** Protein purity was assessed with an SDS-PAGE. 2.72 µM of protein was loaded into each well. Band of interest is denoted by black triangle.

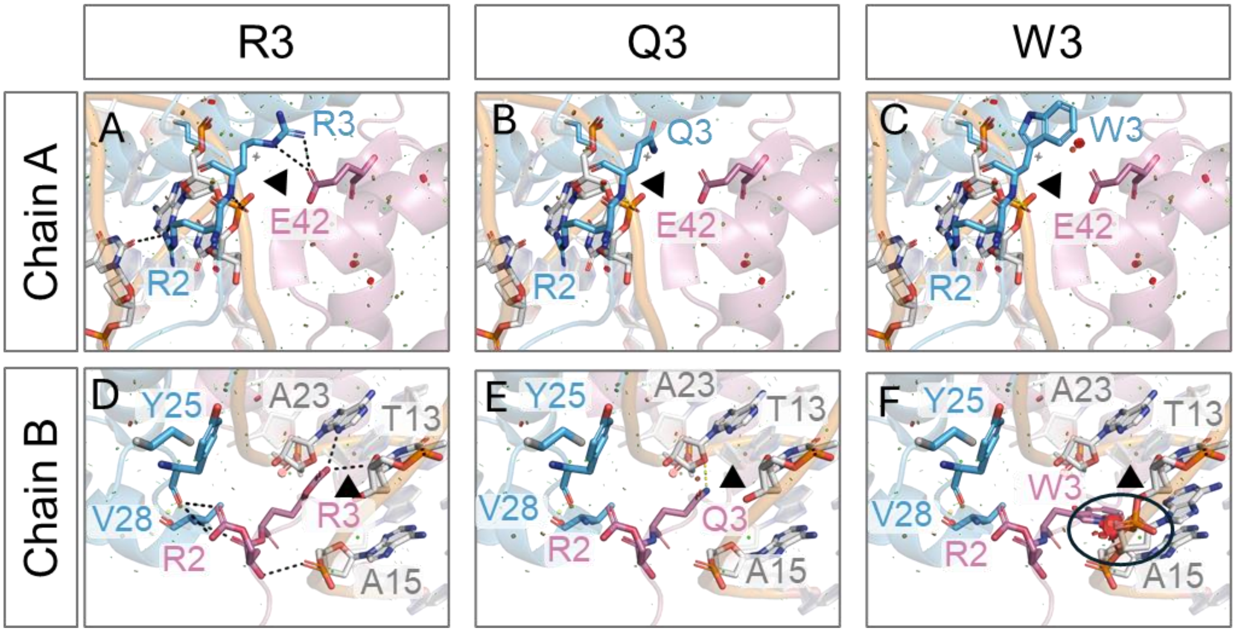


**Supplementary Figure 11.** PYMOL schematics of *in silico* mutagenesis of variants associated with cone-rod dystrophy at position 3 in CRX modeled in the ALX4 dimer structure (9D9R). Chain A is shown in blue, and Chain B is shown in purple. Black dashed lines denote bonds made in wildtype structure. Yellow dashed lines shown denote bonds formed by human disease variant. Steric overlap is denoted by discs where the severity of the clash is denoted by the color and size of the discs.

### 2. Supplemental table captions

**Supplementary Table 1.** Residues that conflict with P3 site cooperativity requirements in Table 1 for each Paired-like and ANTP factor.

**Supplementary Table 2.** Missense variants in cooperative Paired-like factor DNA binding domains associated with disease in HGMD (Stenson et al., 2017, 2020). Functional predictions made by cooperativity rules in Table 1 and DNA contact positions shown in Figure 6A.

**Supplementary Table 3.** Detailed cloning information for protein preparation.
