## Supplementary Information for "Decoding the mechanisms of cooperative DNA binding by the Paired-like homeodomain family"

PCR amplified region    Codon mutated

>ISX in pUC57

```
TCGCGCGTTTCGGTGATGACGGTGAAAACCTCTGACACATGCAGCTCCCGGAGACGGTCACAGCTTGTCTGTAAGCG
GATGCCGGGAGCAGACAAGCCCGTCAGGGCGCGTCAGCGGGTGTTGGCGGGTGTCGGGGCTGGCTTAACTATGCGGC
ATCAGAGCAGATTGTACTGAGAGTGCACCATATGCGGTGTGAAATACCGCACAGATGCGTAAGGAGAAAATACCGCA
TCAGGCGCCATTCGCCATTCAGGCTGCGCAACTGTTGGGAAGGGCGATCGGTGCGGGCCTCTTCGCTATTACGCCAG
CTGGCGAAAGGGGGATGTGCTGCAAGGCGATTAAGTTGGGTAACGCCAGGGTTTTCCCAGTCACGACGTTGTAAAAC
GACGGCCAGTGAATTCGAGCTCGGTACCTCGCGAATGCATCTAGATGAACGCGGCCGCTAGGCAGACAACTATCCCA
GGTTCCAAGCTGGAGAGGCCCCCACAGGATCAGCCCCAAGAAGAGAAAAAGAACAAGCGGAGAGTTCGCACTACCTT
CACCACAGAGCAGCTACAGGAGCTGGAGAAGCTCTTCCACTTCACCCATTACCCTGACATCCATGTCCGTTCCCAAT
TGGCTTCCAGGATTAACCTGCCTGAAGCTCGAGTACAGATCTGGTTCCAGAATCAGAGAGCCAAGTGGAGGAAGCAG
GAGAAGAGTGGGAACCTCAGTGCCCCACAGCAGCCTGGTGAGGCCGGCTTGGCCCTGCCCTCAAACATGGACGTGTC
TGGTTAGGGATCCTAAGATCGGATCCCGGGCCCGTCGACTGCAGAGGCCTGCATGCAAGCTTGGCGTAATCATGGTC
ATAGCTGTTTCCTGTGTGAAATTGTTATCCGCTCACAATTCCACACAACATACGAGCCGGAAGCATAAAGTGTAAAG
CCTGGGGTGCCTAATGAGTGAGCTAACTCACATTAATTGCGTTGCGCTCACTGCCCGCTTTCCAGTCGGGAAACCTG
TCGTGCCAGCTGCATTAATGAATCGGCCAACGCGCGGGGAGAGGCGGTTTGCGTATTGGGCGCTCTTCCGCTTCCTC
GCTCACTGACTCGCTGCGCTCGGTCGTTCGGCTGCGGCGAGCGGTATCAGCTCACTCAAAGGCGGTAATACGGTTAT
CCACAGAATCAGGGGATAACGCAGGAAAGAACATGTGAGCAAAAGGCCAGCAAAAGGCCAGGAACCGTAAAAAGGCC
GCGTTGCTGGCGTTTTTCCATAGGCTCCGCCCCCCTGACGAGCATCACAAAAATCGACGCTCAAGTCAGAGGTGGCG
AAACCCGACAGGACTATAAAGATACCAGGCGTTTCCCCCTGGAAGCTCCCTCGTGCGCTCTCCTGTTCCGACCCTGC
CGCTTACCGGATACCTGTCCGCCTTTCTCCCTTCGGGAAGCGTGGCGCTTTCTCATAGCTCACGCTGTAGGTATCTC
AGTTCGGTGTAGGTCGTTCGCTCCAAGCTGGGCTGTGTGCACGAACCCCCCGTTCAGCCCGACCGCTGCGCCTTATC
CGGTAACTATCGTCTTGAGTCCAACCCGGTAAGACACGACTTATCGCCACTGGCAGCAGCCACTGGTAACAGGATTA
GCAGAGCGAGGTATGTAGGCGGTGCTACAGAGTTCTTGAAGTGGTGGCCTAACTACGGCTACACTAGAAGAACAGTA
TTTGGTATCTGCGCTCTGCTGAAGCCAGTTACCTTCGGAAAAAGAGTTGGTAGCTCTTGATCCGGCAAACAAACCAC
CGCTGGTAGCGGTGGTTTTTTTGTTTGCAAGCAGCAGATTACGCGCAGAAAAAAAGGATCTCAAGAAGATCCTTTGA
TCTTTTCTACGGGGTCTGACGCTCAGTGGAACGAAAACTCACGTTAAGGGATTTTGGTCATGAGATTATCAAAAAGG
ATCTTCACCTAGATCCTTTTAAATTAAAAATGAAGTTTTAAATCAATCTAAAGTATATATGAGTAAACTTGGTCTGA
CAGTTACCAATGCTTAATCAGTGAGGCACCTATCTCAGCGATCTGTCTATTTCGTTCATCCATAGTTGCCTGACTCC
CCGTCGTGTAGATAACTACGATACGGGAGGGCTTACCATCTGGCCCCAGTGCTGCAATGATACCGCGAGACCCACGC
TCACCGGCTCCAGATTTATCAGCAATAAACCAGCCAGCCGGAAGGGCCGAGCGCAGAAGTGGTCCTGCAACTTTATC
CGCCTCCATCCAGTCTATTAATTGTTGCCGGGAAGCTAGAGTAAGTAGTTCGCCAGTTAATAGTTTGCGCAACGTTG
TTGCCATTGCTACAGGCATCGTGGTGTCACGCTCGTCGTTTGGTATGGCTTCATTCAGCTCCGGTTCCCAACGATCA
AGGCGAGTTACATGATCCCCCATGTTGTGCAAAAAAGCGGTTAGCTCCTTCGGTCCTCCGATCGTTGTCAGAAGTAA
GTTGGCCGCAGTGTTATCACTCATGGTTATGGCAGCACTGCATAATTCTCTTACTGTCATGCCATCCGTAAGATGCT
TTTCTGTGACTGGTGAGTACTCAACCAAGTCATTCTGAGAATAGTGTATGCGGCGACCGAGTTGCTCTTGCCCGGCG
TCAATACGGGATAATACCGCGCCACATAGCAGAACTTTAAAAGTGCTCATCATTGGAAAACGTTCTTCGGGGCGAAA
ACTCTCAAGGATCTTACCGCTGTTGAGATCCAGTTCGATGTAACCCACTCGTGCACCCAACTGATCTTCAGCATCTT
TTACTTTCACCAGCGTTTCTGGGTGAGCAAAAACAGGAAGGCAAAATGCCGCAAAAAAGGGAATAAGGGCGACACGG
AAATGTTGAATACTCATACTCTTCCTTTTTCAATATTATTGAAGCATTTATCAGGGTTATTGTCTCATGAGCGGATA
CATATTTGAATGTATTTAGAAAAATAAACAAATAGGGGTTCCGCGCACATTTCCCCGAAAAGTGCCACCTGACGTCT
AAGAAACCATTATTATCATGACATTAACCTATAAAAATAGGCGTATCACGAGGCCCTTTCGTC
```

>ARX E32S in Arx_OMu17806D_pcDNA3.1+/C-(K)-DYK

GACGGATCGGGAGATCTCCCGATCCCCTATGGTGCACTCTCAGTACAATCTGCTCTGATGCCGCATAGTTAAGCCAG
TATCTGCTCCCTGCTTGTGTGTTGGAGGTCGCTGAGTAGTGCGCGAGCAAAATTTAAGCTACAACAAGGCAAGGCTT
GACCGACAATTGCATGAAGAATCTGCTTAGGGTTAGGCGTTTTGCGCTGCTTCGCGATGTACGGGCCAGATATACGC
GTTGACATTGATTATTGACTAGTTATTAATAGTAATCAATTACGGGGTCATTAGTTCATAGCCCATATATGGAGTTC
CGCGTTACATAACTTACGGTAAATGGCCCGCCTGGCTGACCGCCCAACGACCCCCGCCCATTGACGTCAATAATGAC
GTATGTTCCCATAGTAACGCCAATAGGGACTTTCCATTGACGTCAATGGGTGGAGTATTTACGGTAAACTGCCCACT
TGGCAGTACATCAAGTGTATCATATGCCAAGTACGCCCCCTATTGACGTCAATGACGGTAAATGGCCCGCCTGGCAT
TATGCCCAGTACATGACCTTATGGGACTTTCCTACTTGGCAGTACATCTACGTATTAGTCATCGCTATTACCATGGT
GATGCGGTTTTGGCAGTACATCAATGGGCGTGGATAGCGGTTTGACTCACGGGGATTTCCAAGTCTCCACCCCATTG
ACGTCAATGGGAGTTTGTTTTGGCACCAAAATCAACGGGACTTTCCAAAATGTCGTAACAACTCCGCCCCATTGACG
CAAATGGGCGGTAGGCGTGTACGGTGGGAGGTCTATATAAGCAGAGCTCTCTGGCTAACTAGAGAACCCACTGCTTA
CTGGCTTATCGAAATTAATACGACTCACTATAGGGAGACCCAAGCTGGCTAGCGTTTAAACTTAAGCTTGGTACCGA
GCTCGGATCCGCCACCATGAGCAATCAGTACCAGGAAGAGGGCTGCTCTGAGAGGCCGGAGTGCAAGAGTAAATCTC
CAACTTTGCTCTCCTCCTACTGCATCGACAGCATCCTGGGCCGGAGAAGCCCATGCAAAATGAGGCTTCTGGGAGCC
GCGCAGAGCTTGCCTGCCCCGCTGGCCAGCCGCGCTGACCAGGAAAAAGCCATGCAAGGCTCCCCCAAGAGCAGCAG
CGCCCCGTTCGAGGCTGAGCTGCACCTGCCACCCAAGCTGCGGCGCCTCTACGGCCCGGGCGGGGGCCGCCTCCTCC
AGGGCGCGGCGGCGGCGGCTGCAGCGGCGGCGGCGGCCGCAGCGGCGGCCACGGCCACTGGCACCGCAGGTCCCCGT
GGAGAGGTTCCTCCACCGCCACCACCAGCTGCACGGCCAGGGGAGCGTCAGGACAGCGCAGGGGCCGTGGCTGCCGC
CGCTGCCGCCGCTGCCTGGGACACGCTCAAGATCAGCCAGGCGCCGCAGGTGAGCATCAGTCGTAGCAAGTCGTACC
GCGAGAACGGGGCGCCCTTCGTGCCCCCGCCGCCAGCCCTGGACGAGCTGAGTGGCCCGGGGGGCGTCGCGCACCCC
GAGGAGCGCCTCAGCGCGGCCAGCGGACCCGGCAGTGCCCCCGCGGCGGGTGGTGGCACGGGCGCCGAGGACGACGA
GGAGGAGCTGTTGGAGGATGAGGAAGATGAAGAGGAGGAAGAGGAGCTGCTGGAGGACGACGACGAGGAGCTGCTGG
AGGACGATGCCCGCGCGCTGCTCAAGGAGCCCCGGCGCTGCTCGGTGGCCACCACCGGCACAGTGGCCGCCGCCGCC
GCCGCTGCTGCCGCCGCCGTGGCCACCGAGGGCGGGGAGCTGTCGCCCAAGGAAGAGCTGTTGCTGCACCCGGAGGA
CGCCGAGGGCAAGGATGGTGAGGACAGCGTGTGCCTGTCGGCCGGCAGCGACTCGGAGGAGGGGCTGCTGAAGCGCA
AACAGAGGCGCTACCGCACCACGTTCACCAGTTACCAGCTGGAGGAACTGGAGCGGGCTTTCCAGAAGACGCACTAC
CCTGACGTCTTCACCAGGTCCGAGCTGGCCATGAGGCTGGACCTGACAGAGGCCCGAGTCCAGGTGTGGTTCCAGAA
CCGTCGGGCCAAGTGGCGCAAGCGGGAGAAGGCTGGCGCACAGACCCACCCCCCGGGGCTGCCCTTTCCAGGGCCGC
TCTCGGCTACCCACCCTCTCAGCCCCTACCTGGACGCCAGCCCCTTTCCCCCGCATCACCCTGCGCTTGATTCGGCC
TGGACCGCCGCGGCGGCGGCGGCTGCAGCTGCCTTCCCGAGCCTACCTCCGCCCCCAGGCTCTGCTAGCCTGCCACC
CAGCGGGGCGCCGCTGGGTCTGAGCACTTTTCTAGGAGCAGCGGTGTTCCGCCACCCAGCCTTCATCAGCCCAGCAT
TTGGCAGGCTCTTTTCCACTATGGCCCCCCTGACCAGCGCGTCGACTGCGGCTGCGCTCCTGAGACAGCCCACACCC
GCGGTGGAGGGCGCAGTGGCCTCGGGCGCGCTGGCCGACCCGGCCACCGCCGCGGCAGACAGACGCGCGTCAAGCAT
AGCCGCGCTGAGGCTCAAGGCTAAGGAGCACGCCGCGCAGCTCACGCAGCTCAACATCCTGCCGGGCACCAGCACGG
GGAAGGAGGTGTGCGATTACAAGGATGACGACGATAAGTGATAAACCCGCTGATCAGCCTCGACTGTGCCTTCTAGT
TGCCAGCCATCTGTTGTTTGCCCCTCCCCCGTGCCTTCCTTGACCCTGGAAGGTGCCACTCCCACTGTCCTTTCCTA
ATAAAATGAGGAAATTGCATCGCATTGTCTGAGTAGGTGTCATTCTATTCTGGGGGGTGGGGTGGGGCAGGACAGCA
AGGGGGAGGATTGGGAAGACAATAGCAGGCATGCTGGGGATGCGGTGGGCTCTATGGCTTCTGAGGCGGAAAGAACC
AGCTGGGGCTCTAGGGGGTATCCCCACGCGCCCTGTAGCGGCGCATTAAGCGCGGCGGGTGTGGTGGTTACGCGCAG
CGTGACCGCTACACTTGCCAGCGCCCTAGCGCCCGCTCCTTTCGCTTTCTTCCCTTCCTTTCTCGCCACGTTCGCCG
GCTTTCCCCGTCAAGCTCTAAATCGGGGGCTCCCTTTAGGGTTCCGATTTAGTGCTTTACGGCACCTCGACCCCAAA
AAACTTGATTAGGGTGATGGTTCACGTAGTGGGCCATCGCCCTGATAGACGGTTTTTCGCCCTTTGACGTTGGAGTC

CACGTTCTTTAATAGTGGACTCTTGTTCCAAACTGGAACAACACTCAACCCTATCTCGGTCTATTCTTTTGATTTAT
AAGGGATTTTGCCGATTTCGGCCTATTGGTTAAAAAATGAGCTGATTTAACAAAAATTTAACGCGAATTAATTCTGT
GGAATGTGTGTCAGTTAGGGTGTGGAAAGTCCCCAGGCTCCCCAGCAGGCAGAAGTATGCAAAGCATGCATCTCAAT
TAGTCAGCAACCAGGTGTGGAAAGTCCCCAGGCTCCCCAGCAGGCAGAAGTATGCAAAGCATGCATCTCAATTAGTC
AGCAACCATAGTCCCGCCCCTAACTCCGCCCATCCCGCCCCTAACTCCGCCCAGTTCCGCCCATTCTCCGCCCCATG
GCTGACTAATTTTTTTTATTTATGCAGAGGCCGAGGCCGCCTCTGCCTCTGAGCTATTCCAGAAGTAGTGAGGAGGC
TTTTTTGGAGGCCTAGGCTTTTGCAAAAAGCTCCCGGGAGCTTGTATATCCATTTTCGGATCTGATCAAGAGACAGG
ATGAGGATCGTTTCGCATGATTGAACAAGATGGATTGCACGCAGGTTCTCCGGCCGCTTGGGTGGAGAGGCTATTCG
GCTATGACTGGGCACAACAGACAATCGGCTGCTCTGATGCCGCCGTGTTCCGGCTGTCAGCGCAGGGGCGCCCGGTT
CTTTTTGTCAAGACCGACCTGTCCGGTGCCCTGAATGAACTGCAGGACGAGGCAGCGCGGCTATCGTGGCTGGCCAC
GACGGGCGTTCCTTGCGCAGCTGTGCTCGACGTTGTCACTGAAGCGGGAAGGGACTGGCTGCTATTGGGCGAAGTGC
CGGGGCAGGATCTCCTGTCATCTCACCTTGCTCCTGCCGAGAAAGTATCCATCATGGCTGATGCAATGCGGCGGCTG
CATACGCTTGATCCGGCTACCTGCCCATTCGACCACCAAGCGAAACATCGCATCGAGCGAGCACGTACTCGGATGGA
AGCCGGTCTTGTCGATCAGGATGATCTGGACGAAGAGCATCAGGGGCTCGCGCCAGCCGAACTGTTCGCCAGGCTCA
AGGCGCGCATGCCCGACGGCGAGGATCTCGTCGTGACCCATGGCGATGCCTGCTTGCCGAATATCATGGTGGAAAAT
GGCCGCTTTTCTGGATTCATCGACTGTGGCCGGCTGGGTGTGGCGGACCGCTATCAGGACATAGCGTTGGCTACCCG
TGATATTGCTGAAGAGCTTGGCGGCGAATGGGCTGACCGCTTCCTCGTGCTTTACGGTATCGCCGCTCCCGATTCGC
AGCGCATCGCCTTCTATCGCCTTCTTGACGAGTTCTTCTGAGCGGGACTCTGGGGTTCGAAATGACCGACCAAGCGA
CGCCCAACCTGCCATCACGAGATTTCGATTCCACCGCCGCCTTCTATGAAAGGTTGGGCTTCGGAATCGTTTTCCGG
GACGCCGGCTGGATGATCCTCCAGCGCGGGGATCTCATGCTGGAGTTCTTCGCCCACCCCAACTTGTTTATTGCAGC
TTATAATGGTTACAAATAAAGCAATAGCATCACAAATTTCACAAATAAAGCATTTTTTTCACTGCATTCTAGTTGTG
GTTTGTCCAAACTCATCAATGTATCTTATCATGTCTGTATACCGTCGACCTCTAGCTAGAGCTTGGCGTAATCATGG
TCATAGCTGTTTCCTGTGTGAAATTGTTATCCGCTCACAATTCCACACAACATACGAGCCGGAAGCATAAAGTGTAA
AGCCTGGGGTGCCTAATGAGTGAGCTAACTCACATTAATTGCGTTGCGCTCACTGCCCGCTTTCCAGTCGGGAAACC
TGTCGTGCCAGCTGCATTAATGAATCGGCCAACGCGCGGGGAGAGGCGGTTTGCGTATTGGGCGCTCTTCCGCTTCC
TCGCTCACTGACTCGCTGCGCTCGGTCGTTCGGCTGCGGCGAGCGGTATCAGCTCACTCAAAGGCGGTAATACGGTT
ATCCACAGAATCAGGGGATAACGCAGGAAAGAACATGTGAGCAAAAGGCCAGCAAAAGGCCAGGAACCGTAAAAAGG
CCGCGTTGCTGGCGTTTTTCCATAGGCTCCGCCCCCCTGACGAGCATCACAAAAATCGACGCTCAAGTCAGAGGTGG
CGAAACCCGACAGGACTATAAAGATACCAGGCGTTTCCCCCTGGAAGCTCCCTCGTGCGCTCTCCTGTTCCGACCCT
GCCGCTTACCGGATACCTGTCCGCCTTTCTCCCTTCGGGAAGCGTGGCGCTTTCTCATAGCTCACGCTGTAGGTATC
TCAGTTCGGTGTAGGTCGTTCGCTCCAAGCTGGGCTGTGTGCACGAACCCCCCGTTCAGCCCGACCGCTGCGCCTTA
TCCGGTAACTATCGTCTTGAGTCCAACCCGGTAAGACACGACTTATCGCCACTGGCAGCAGCCACTGGTAACAGGAT
TAGCAGAGCGAGGTATGTAGGCGGTGCTACAGAGTTCTTGAAGTGGTGGCCTAACTACGGCTACACTAGAAGAACAG
TATTTGGTATCTGCGCTCTGCTGAAGCCAGTTACCTTCGGAAAAAGAGTTGGTAGCTCTTGATCCGGCAAACAAACC
ACCGCTGGTAGCGGTGGTTTTTTTGTTTGCAAGCAGCAGATTACGCGCAGAAAAAAAGGATCTCAAGAAGATCCTTT
GATCTTTTCTACGGGGTCTGACGCTCAGTGGAACGAAAACTCACGTTAAGGGATTTTGGTCATGAGATTATCAAAAA
GGATCTTCACCTAGATCCTTTTAAATTAAAAATGAAGTTTTAAATCAATCTAAAGTATATATGAGTAAACTTGGTCT
GACAGTTACCAATGCTTAATCAGTGAGGCACCTATCTCAGCGATCTGTCTATTTCGTTCATCCATAGTTGCCTGACT
CCCCGTCGTGTAGATAACTACGATACGGGAGGGCTTACCATCTGGCCCCAGTGCTGCAATGATACCGCGAGACCCAC
GCTCACCGGCTCCAGATTTATCAGCAATAAACCAGCCAGCCGGAAGGGCCGAGCGCAGAAGTGGTCCTGCAACTTTA
TCCGCCTCCATCCAGTCTATTAATTGTTGCCGGGAAGCTAGAGTAAGTAGTTCGCCAGTTAATAGTTTGCGCAACGT
TGTTGCCATTGCTACAGGCATCGTGGTGTCACGCTCGTCGTTTGGTATGGCTTCATTCAGCTCCGGTTCCCAACGAT
CAAGGCGAGTTACATGATCCCCCATGTTGTGCAAAAAAGCGGTTAGCTCCTTCGGTCCTCCGATCGTTGTCAGAAGT

AAGTTGGCCGCAGTGTTATCACTCATGGTTATGGCAGCACTGCATAATTCTCTTACTGTCATGCCATCCGTAAGATG
CTTTTCTGTGACTGGTGAGTACTCAACCAAGTCATTCTGAGAATAGTGTATGCGGCGACCGAGTTGCTCTTGCCCGG
CGTCAATACGGGATAATACCGCGCCACATAGCAGAACTTTAAAAGTGCTCATCATTGGAAAACGTTCTTCGGGGCGA
AAACTCTCAAGGATCTTACCGCTGTTGAGATCCAGTTCGATGTAACCCACTCGTGCACCCAACTGATCTTCAGCATC
TTTTACTTTCACCAGCGTTTCTGGGTGAGCAAAAACAGGAAGGCAAAATGCCGCAAAAAAGGGAATAAGGGCGACAC
GGAAATGTTGAATACTCATACTCTTCCTTTTTCAATATTATTGAAGCATTTATCAGGGTTATTGTCTCATGAGCGGA
TACATATTTGAATGTATTTAGAAAAATAAACAAATAGGGGTTCCGCGCACATTTCCCCGAAAAGTGCCACCTGACGT
C

>ISX S32E in pUC57

TCGCGCGTTTCGGTGATGACGGTGAAAACCTCTGACACATGCAGCTCCCGGAGACGGTCACAGCTTGTCTGTAAGCG
GATGCCGGGAGCAGACAAGCCCGTCAGGGCGCGTCAGCGGGTGTTGGCGGGTGTCGGGGCTGGCTTAACTATGCGGC
ATCAGAGCAGATTGTACTGAGAGTGCACCATATGCGGTGTGAAATACCGCACAGATGCGTAAGGAGAAAATACCGCA
TCAGGCGCCATTCGCCATTCAGGCTGCGCAACTGTTGGGAAGGGCGATCGGTGCGGGCCTCTTCGCTATTACGCCAG
CTGGCGAAAGGGGGATGTGCTGCAAGGCGATTAAGTTGGGTAACGCCAGGGTTTTCCCAGTCACGACGTTGTAAAAC
GACGGCCAGTGAATTCGAGCTCGGTACCTCGCGAATGCATCTAGATGAACGCGGCCGCTAGGCAGACAACTATCCCA
GGTTCCAAGCTGGAGAGGCCCCCACAGGATCAGCCCCAAGAAGAGAAAAAGAACAAGCGGAGAGTTCGCACTACCTT
CACCACAGAGCAGCTACAGGAGCTGGAGAAGCTCTTCCACTTCACCCATTACCCTGACATCCATGTCCGTGAACAAT
TGGCTTCCAGGATTAACCTGACTGAAGCTCGAGTACAGATCTGGTTCCAGAATCAGAGAGCCAAGTGGAGGAAGCAG
GAGAAGAGTGGGAACCTCAGTGCCCCACAGCAGCCTGGTGAGGCCGGCTTGGCCCTGCCCTCAAACATGGACGTGTC
TGGTTAGGGATCCTAAGATCGGATCCCGGGCCCGTCGACTGCAGAGGCCTGCATGCAAGCTTGGCGTAATCATGGTC
ATAGCTGTTTCCTGTGTGAAATTGTTATCCGCTCACAATTCCACACAACATACGAGCCGGAAGCATAAAGTGTAAAG
CCTGGGGTGCCTAATGAGTGAGCTAACTCACATTAATTGCGTTGCGCTCACTGCCCGCTTTCCAGTCGGGAAACCTG
TCGTGCCAGCTGCATTAATGAATCGGCCAACGCGCGGGGAGAGGCGGTTTGCGTATTGGGCGCTCTTCCGCTTCCTC
GCTCACTGACTCGCTGCGCTCGGTCGTTCGGCTGCGGCGAGCGGTATCAGCTCACTCAAAGGCGGTAATACGGTTAT
CCACAGAATCAGGGGATAACGCAGGAAAGAACATGTGAGCAAAAGGCCAGCAAAAGGCCAGGAACCGTAAAAAGGCC
GCGTTGCTGGCGTTTTTCCATAGGCTCCGCCCCCCTGACGAGCATCACAAAAATCGACGCTCAAGTCAGAGGTGGCG
AAACCCGACAGGACTATAAAGATACCAGGCGTTTCCCCCTGGAAGCTCCCTCGTGCGCTCTCCTGTTCCGACCCTGC
CGCTTACCGGATACCTGTCCGCCTTTCTCCCTTCGGGAAGCGTGGCGCTTTCTCATAGCTCACGCTGTAGGTATCTC
AGTTCGGTGTAGGTCGTTCGCTCCAAGCTGGGCTGTGTGCACGAACCCCCCGTTCAGCCCGACCGCTGCGCCTTATC
CGGTAACTATCGTCTTGAGTCCAACCCGGTAAGACACGACTTATCGCCACTGGCAGCAGCCACTGGTAACAGGATTA
GCAGAGCGAGGTATGTAGGCGGTGCTACAGAGTTCTTGAAGTGGTGGCCTAACTACGGCTACACTAGAAGAACAGTA
TTTGGTATCTGCGCTCTGCTGAAGCCAGTTACCTTCGGAAAAAGAGTTGGTAGCTCTTGATCCGGCAAACAAACCAC
CGCTGGTAGCGGTGGTTTTTTTGTTTGCAAGCAGCAGATTACGCGCAGAAAAAAAGGATCTCAAGAAGATCCTTTGA
TCTTTTCTACGGGGTCTGACGCTCAGTGGAACGAAAACTCACGTTAAGGGATTTTGGTCATGAGATTATCAAAAAGG
ATCTTCACCTAGATCCTTTTAAATTAAAAATGAAGTTTTAAATCAATCTAAAGTATATATGAGTAAACTTGGTCTGA
CAGTTACCAATGCTTAATCAGTGAGGCACCTATCTCAGCGATCTGTCTATTTCGTTCATCCATAGTTGCCTGACTCC
CCGTCGTGTAGATAACTACGATACGGGAGGGCTTACCATCTGGCCCCAGTGCTGCAATGATACCGCGAGACCCACGC
TCACCGGCTCCAGATTTATCAGCAATAAACCAGCCAGCCGGAAGGGCCGAGCGCAGAAGTGGTCCTGCAACTTTATC
CGCCTCCATCCAGTCTATTAATTGTTGCCGGGAAGCTAGAGTAAGTAGTTCGCCAGTTAATAGTTTGCGCAACGTTG
TTGCCATTGCTACAGGCATCGTGGTGTCACGCTCGTCGTTTGGTATGGCTTCATTCAGCTCCGGTTCCCAACGATCA
AGGCGAGTTACATGATCCCCCATGTTGTGCAAAAAAGCGGTTAGCTCCTTCGGTCCTCCGATCGTTGTCAGAAGTAA
GTTGGCCGCAGTGTTATCACTCATGGTTATGGCAGCACTGCATAATTCTCTTACTGTCATGCCATCCGTAAGATGCT
TTTCTGTGACTGGTGAGTACTCAACCAAGTCATTCTGAGAATAGTGTATGCGGCGACCGAGTTGCTCTTGCCCGGCG
TCAATACGGGATAATACCGCGCCACATAGCAGAACTTTAAAAGTGCTCATCATTGGAAAACGTTCTTCGGGGCGAAA
ACTCTCAAGGATCTTACCGCTGTTGAGATCCAGTTCGATGTAACCCACTCGTGCACCCAACTGATCTTCAGCATCTT
TTACTTTCACCAGCGTTTCTGGGTGAGCAAAAACAGGAAGGCAAAATGCCGCAAAAAAGGGAATAAGGGCGACACGG
AAATGTTGAATACTCATACTCTTCCTTTTTCAATATTATTGAAGCATTTATCAGGGTTATTGTCTCATGAGCGGATA
CATATTTGAATGTATTTAGAAAAATAAACAAATAGGGGTTCCGCGCACATTTCCCCGAAAAGTGCCACCTGACGTCT
AAGAAACCATTATTATCATGACATTAACCTATAAAAATAGGCGTATCACGAGGCCCTTTCGTC

>PHOX2B Q46R in PHOX2B_OHu26329C_pcDNA3.1(+)-N-HA

GACGGATCGGGAGATCTCCCGATCCCCTATGGTGCACTCTCAGTACAATCTGCTCTGATGCCGCATAGTTAAGCCAG
TATCTGCTCCCTGCTTGTGTGTTGGAGGTCGCTGAGTAGTGCGCGAGCAAAATTTAAGCTACAACAAGGCAAGGCTT
GACCGACAATTGCATGAAGAATCTGCTTAGGGTTAGGCGTTTTGCGCTGCTTCGCGATGTACGGGCCAGATATACGC
GTTGACATTGATTATTGACTAGTTATTAATAGTAATCAATTACGGGGTCATTAGTTCATAGCCCATATATGGAGTTC
CGCGTTACATAACTTACGGTAAATGGCCCGCCTGGCTGACCGCCCAACGACCCCCGCCCATTGACGTCAATAATGAC
GTATGTTCCCATAGTAACGCCAATAGGGACTTTCCATTGACGTCAATGGGTGGAGTATTTACGGTAAACTGCCCACT
TGGCAGTACATCAAGTGTATCATATGCCAAGTACGCCCCCTATTGACGTCAATGACGGTAAATGGCCCGCCTGGCAT
TATGCCCAGTACATGACCTTATGGGACTTTCCTACTTGGCAGTACATCTACGTATTAGTCATCGCTATTACCATGGT
GATGCGGTTTTGGCAGTACATCAATGGGCGTGGATAGCGGTTTGACTCACGGGGATTTCCAAGTCTCCACCCCATTG
ACGTCAATGGGAGTTTGTTTTGGCACCAAAATCAACGGGACTTTCCAAAATGTCGTAACAACTCCGCCCCATTGACG
CAAATGGGCGGTAGGCGTGTACGGTGGGAGGTCTATATAAGCAGAGCTCTCTGGCTAACTAGAGAACCCACTGCTTA
CTGGCTTATCGAAATTAATACGACTCACTATAGGGAGACCCAAGCTGGCTAGCGTTTAAACTTAAGCTTGCCACCAT
GTACCCATACGATGTTCCAGATTACGCTGGCTCTGGATATAAAATGGAATATTCTTACCTCAATTCCTCTGCCTACG
AGTCCTGTATGGCTGGGATGGACACCTCGAGCCTGGCTTCAGCCTATGCTGACTTCAGTTCCTGCAGCCAGGCCAGT
GGCTTCCAGTATAACCCGATAAGGACCACTTTTGGGGCCACGTCCGGCTGCCCTTCCCTCACGCCGGGATCCTGCAG
CCTGGGCACCCTCAGGGACCACCAGAGCAGTCCGTACGCCGCAGTTCCTTACAAACTCTTCACGGACCACGGCGGCC
TCAACGAGAAGCGCAAGCAGCGGCGCATCCGCACCACTTTCACCAGTGCCCAGCTCAAAGAGCTGGAAAGGGTCTTC
GCGGAGACTCACTACCCCGACATCTACACTCGGGAGGAGCTGGCCCTGAAGATCGACCTCACAGAGGCGCGAGTCCG
GGTGTGGTTCCAGAACCGCCGCGCCAAGTTTCGCAAGCAGGAGCGCGCAGCGGCAGCCGCAGCGGCCGCGGCCAAGA
ACGGCTCCTCGGGCAAAAAGTCTGACTCTTCCAGGGACGACGAGAGCAAAGAGGCCAAGAGCACTGACCCGGACAGC
ACTGGGGGCCCAGGTCCCAATCCCAACCCCACCCCCAGCTGCGGGGCGAATGGAGGCGGCGGCGGCGGGCCCAGCCC
GGCTGGAGCTCCGGGGGCGGCGGGGCCCGGGGGCCCGGGAGGCGAACCCGGCAAGGGCGGCGCAGCAGCAGCGGCGG
CGGCCGCGGCAGCGGCGGCGGCGGCAGCGGCAGCGGCGGCAGCTGGAGGCCTGGCTGCGGCTGGGGGCCCTGGACAA
GGCTGGGCTCCCGGCCCCGGCCCCATCACCTCCATCCCGGATTCGCTTGGGGGTCCCTTCGCCAGCGTCCTATCTTC
GCTCCAAAGACCCAACGGTGCCAAAGCCGCCTTAGTGAAGAGCAGTATGTTCTGAGGTACCGAGCTCGGATCCGAAT
TCTGCAGATATCCAGCACAGTGGCGGCCGCTCGAGTCTAGAGGGCCCGTTTAAACCCGCTGATCAGCCTCGACTGTG
CCTTCTAGTTGCCAGCCATCTGTTGTTTGCCCCTCCCCCGTGCCTTCCTTGACCCTGGAAGGTGCCACTCCCACTGT
CCTTTCCTAATAAAATGAGGAAATTGCATCGCATTGTCTGAGTAGGTGTCATTCTATTCTGGGGGGTGGGGTGGGGC
AGGACAGCAAGGGGGAGGATTGGGAAGACAATAGCAGGCATGCTGGGGATGCGGTGGGCTCTATGGCTTCTGAGGCG
GAAAGAACCAGCTGGGGCTCTAGGGGGTATCCCCACGCGCCCTGTAGCGGCGCATTAAGCGCGGCGGGTGTGGTGGT
TACGCGCAGCGTGACCGCTACACTTGCCAGCGCCCTAGCGCCCGCTCCTTTCGCTTTCTTCCCTTCCTTTCTCGCCA
CGTTCGCCGGCTTTCCCCGTCAAGCTCTAAATCGGGGGCTCCCTTTAGGGTTCCGATTTAGTGCTTTACGGCACCTC
GACCCCAAAAAACTTGATTAGGGTGATGGTTCACGTAGTGGGCCATCGCCCTGATAGACGGTTTTTCGCCCTTTGAC
GTTGGAGTCCACGTTCTTTAATAGTGGACTCTTGTTCCAAACTGGAACAACACTCAACCCTATCTCGGTCTATTCTT
TTGATTTATAAGGGATTTTGCCGATTTCGGCCTATTGGTTAAAAAATGAGCTGATTTAACAAAAATTTAACGCGAAT
TAATTCTGTGGAATGTGTGTCAGTTAGGGTGTGGAAAGTCCCCAGGCTCCCCAGCAGGCAGAAGTATGCAAAGCATG
CATCTCAATTAGTCAGCAACCAGGTGTGGAAAGTCCCCAGGCTCCCCAGCAGGCAGAAGTATGCAAAGCATGCATCT
CAATTAGTCAGCAACCATAGTCCCGCCCCTAACTCCGCCCATCCCGCCCCTAACTCCGCCCAGTTCCGCCCATTCTC
CGCCCCATGGCTGACTAATTTTTTTTATTTATGCAGAGGCCGAGGCCGCCTCTGCCTCTGAGCTATTCCAGAAGTAG
TGAGGAGGCTTTTTTGGAGGCCTAGGCTTTTGCAAAAAGCTCCCGGGAGCTTGTATATCCATTTTCGGATCTGATCA
AGAGACAGGATGAGGATCGTTTCGCATGATTGAACAAGATGGATTGCACGCAGGTTCTCCGGCCGCTTGGGTGGAGA
GGCTATTCGGCTATGACTGGGCACAACAGACAATCGGCTGCTCTGATGCCGCCGTGTTCCGGCTGTCAGCGCAGGGG

CGCCCGGTTCTTTTTGTCAAGACCGACCTGTCCGGTGCCCTGAATGAACTGCAGGACGAGGCAGCGCGGCTATCGTG
GCTGGCCACGACGGGCGTTCCTTGCGCAGCTGTGCTCGACGTTGTCACTGAAGCGGGAAGGGACTGGCTGCTATTGG
GCGAAGTGCCGGGGCAGGATCTCCTGTCATCTCACCTTGCTCCTGCCGAGAAAGTATCCATCATGGCTGATGCAATG
CGGCGGCTGCATACGCTTGATCCGGCTACCTGCCCATTCGACCACCAAGCGAAACATCGCATCGAGCGAGCACGTAC
TCGGATGGAAGCCGGTCTTGTCGATCAGGATGATCTGGACGAAGAGCATCAGGGGCTCGCGCCAGCCGAACTGTTCG
CCAGGCTCAAGGCGCGCATGCCCGACGGCGAGGATCTCGTCGTGACCCATGGCGATGCCTGCTTGCCGAATATCATG
GTGGAAAATGGCCGCTTTTCTGGATTCATCGACTGTGGCCGGCTGGGTGTGGCGGACCGCTATCAGGACATAGCGTT
GGCTACCCGTGATATTGCTGAAGAGCTTGGCGGCGAATGGGCTGACCGCTTCCTCGTGCTTTACGGTATCGCCGCTC
CCGATTCGCAGCGCATCGCCTTCTATCGCCTTCTTGACGAGTTCTTCTGAGCGGGACTCTGGGGTTCGAAATGACCG
ACCAAGCGACGCCCAACCTGCCATCACGAGATTTCGATTCCACCGCCGCCTTCTATGAAAGGTTGGGCTTCGGAATC
GTTTTCCGGGACGCCGGCTGGATGATCCTCCAGCGCGGGGATCTCATGCTGGAGTTCTTCGCCCACCCCAACTTGTT
TATTGCAGCTTATAATGGTTACAAATAAAGCAATAGCATCACAAATTTCACAAATAAAGCATTTTTTTCACTGCATT
CTAGTTGTGGTTTGTCCAAACTCATCAATGTATCTTATCATGTCTGTATACCGTCGACCTCTAGCTAGAGCTTGGCG
TAATCATGGTCATAGCTGTTTCCTGTGTGAAATTGTTATCCGCTCACAATTCCACACAACATACGAGCCGGAAGCAT
AAAGTGTAAAGCCTGGGGTGCCTAATGAGTGAGCTAACTCACATTAATTGCGTTGCGCTCACTGCCCGCTTTCCAGT
CGGGAAACCTGTCGTGCCAGCTGCATTAATGAATCGGCCAACGCGCGGGGAGAGGCGGTTTGCGTATTGGGCGCTCT
TCCGCTTCCTCGCTCACTGACTCGCTGCGCTCGGTCGTTCGGCTGCGGCGAGCGGTATCAGCTCACTCAAAGGCGGT
AATACGGTTATCCACAGAATCAGGGGATAACGCAGGAAAGAACATGTGAGCAAAAGGCCAGCAAAAGGCCAGGAACC
GTAAAAAGGCCGCGTTGCTGGCGTTTTTCCATAGGCTCCGCCCCCCTGACGAGCATCACAAAAATCGACGCTCAAGT
CAGAGGTGGCGAAACCCGACAGGACTATAAAGATACCAGGCGTTTCCCCCTGGAAGCTCCCTCGTGCGCTCTCCTGT
TCCGACCCTGCCGCTTACCGGATACCTGTCCGCCTTTCTCCCTTCGGGAAGCGTGGCGCTTTCTCATAGCTCACGCT
GTAGGTATCTCAGTTCGGTGTAGGTCGTTCGCTCCAAGCTGGGCTGTGTGCACGAACCCCCCGTTCAGCCCGACCGC
TGCGCCTTATCCGGTAACTATCGTCTTGAGTCCAACCCGGTAAGACACGACTTATCGCCACTGGCAGCAGCCACTGG
TAACAGGATTAGCAGAGCGAGGTATGTAGGCGGTGCTACAGAGTTCTTGAAGTGGTGGCCTAACTACGGCTACACTA
GAAGAACAGTATTTGGTATCTGCGCTCTGCTGAAGCCAGTTACCTTCGGAAAAAGAGTTGGTAGCTCTTGATCCGGC
AAACAAACCACCGCTGGTAGCGGTGGTTTTTTTGTTTGCAAGCAGCAGATTACGCGCAGAAAAAAAGGATCTCAAGA
AGATCCTTTGATCTTTTCTACGGGGTCTGACGCTCAGTGGAACGAAAACTCACGTTAAGGGATTTTGGTCATGAGAT
TATCAAAAAGGATCTTCACCTAGATCCTTTTAAATTAAAAATGAAGTTTTAAATCAATCTAAAGTATATATGAGTAA
ACTTGGTCTGACAGTTACCAATGCTTAATCAGTGAGGCACCTATCTCAGCGATCTGTCTATTTCGTTCATCCATAGT
TGCCTGACTCCCCGTCGTGTAGATAACTACGATACGGGAGGGCTTACCATCTGGCCCCAGTGCTGCAATGATACCGC
GAGACCCACGCTCACCGGCTCCAGATTTATCAGCAATAAACCAGCCAGCCGGAAGGGCCGAGCGCAGAAGTGGTCCT
GCAACTTTATCCGCCTCCATCCAGTCTATTAATTGTTGCCGGGAAGCTAGAGTAAGTAGTTCGCCAGTTAATAGTTT
GCGCAACGTTGTTGCCATTGCTACAGGCATCGTGGTGTCACGCTCGTCGTTTGGTATGGCTTCATTCAGCTCCGGTT
CCCAACGATCAAGGCGAGTTACATGATCCCCCATGTTGTGCAAAAAAGCGGTTAGCTCCTTCGGTCCTCCGATCGTT
GTCAGAAGTAAGTTGGCCGCAGTGTTATCACTCATGGTTATGGCAGCACTGCATAATTCTCTTACTGTCATGCCATC
CGTAAGATGCTTTTCTGTGACTGGTGAGTACTCAACCAAGTCATTCTGAGAATAGTGTATGCGGCGACCGAGTTGCT
CTTGCCCGGCGTCAATACGGGATAATACCGCGCCACATAGCAGAACTTTAAAAGTGCTCATCATTGGAAAACGTTCT
TCGGGGCGAAAACTCTCAAGGATCTTACCGCTGTTGAGATCCAGTTCGATGTAACCCACTCGTGCACCCAACTGATC
TTCAGCATCTTTTACTTTCACCAGCGTTTCTGGGTGAGCAAAAACAGGAAGGCAAAATGCCGCAAAAAAGGGAATAA
GGGCGACACGGAAATGTTGAATACTCATACTCTTCCTTTTTCAATATTATTGAAGCATTTATCAGGGTTATTGTCTC
ATGAGCGGATACATATTTGAATGTATTTAGAAAAATAAACAAATAGGGGTTCCGCGCACATTTCCCCGAAAAGTGCC
ACCTGACGTC

>PITX3 R46Q in pUC57

TCGCGCGTTTCGGTGATGACGGTGAAAACCTCTGACACATGCAGCTCCCGGAGACGGTCACAGCTTGTCTGTAAGCG
GATGCCGGGAGCAGACAAGCCCGTCAGGGCGCGTCAGCGGGTGTTGGCGGGTGTCGGGGCTGGCTTAACTATGCGGC
ATCAGAGCAGATTGTACTGAGAGTGCACCATATGCGGTGTGAAATACCGCACAGATGCGTAAGGAGAAAATACCGCA
TCAGGCGCCATTCGCCATTCAGGCTGCGCAACTGTTGGGAAGGGCGATCGGTGCGGGCCTCTTCGCTATTACGCCAG
CTGGCGAAAGGGGGATGTGCTGCAAGGCGATTAAGTTGGGTAACGCCAGGGTTTTCCCAGTCACGACGTTGTAAAAC
GACGGCCAGTGAATTCGAGCTCGGTACCCATATGTCGCTGAAAAAGAAGCAGCGGCGGCAGCGCACGCACTTCACCA
GCCAGCAGCTACAGGAGCTAGAGGCGACCTTCCAGAGGAACCGCTACCCCGACATGAGCACGCGCGAGGAGATCGCC
GTGTGGACCAACCTCACCGAGGCCCGCGTGCAGGTGTGGTTCAAGAACCGGCGCGCCAAATGGCGGAAGCGCGAGCG
CTAACTCGAGGGATCCCGGGCCCGTCGACTGCAGAGGCCTGCATGCAAGCTTGGCGTAATCATGGTCATAGCTGTTT
CCTGTGTGAAATTGTTATCCGCTCACAATTCCACACAACATACGAGCCGGAAGCATAAAGTGTAAAGCCTGGGGTGC
CTAATGAGTGAGCTAACTCACATTAATTGCGTTGCGCTCACTGCCCGCTTTCCAGTCGGGAAACCTGTCGTGCCAGC
TGCATTAATGAATCGGCCAACGCGCGGGGAGAGGCGGTTTGCGTATTGGGCGCTCTTCCGCTTCCTCGCTCACTGAC
TCGCTGCGCTCGGTCGTTCGGCTGCGGCGAGCGGTATCAGCTCACTCAAAGGCGGTAATACGGTTATCCACAGAATC
AGGGGATAACGCAGGAAAGAACATGTGAGCAAAAGGCCAGCAAAAGGCCAGGAACCGTAAAAAGGCCGCGTTGCTGG
CGTTTTTCCATAGGCTCCGCCCCCCTGACGAGCATCACAAAAATCGACGCTCAAGTCAGAGGTGGCGAAACCCGACA
GGACTATAAAGATACCAGGCGTTTCCCCCTGGAAGCTCCCTCGTGCGCTCTCCTGTTCCGACCCTGCCGCTTACCGG
ATACCTGTCCGCCTTTCTCCCTTCGGGAAGCGTGGCGCTTTCTCATAGCTCACGCTGTAGGTATCTCAGTTCGGTGT
AGGTCGTTCGCTCCAAGCTGGGCTGTGTGCACGAACCCCCCGTTCAGCCCGACCGCTGCGCCTTATCCGGTAACTAT
CGTCTTGAGTCCAACCCGGTAAGACACGACTTATCGCCACTGGCAGCAGCCACTGGTAACAGGATTAGCAGAGCGAG
GTATGTAGGCGGTGCTACAGAGTTCTTGAAGTGGTGGCCTAACTACGGCTACACTAGAAGAACAGTATTTGGTATCT
GCGCTCTGCTGAAGCCAGTTACCTTCGGAAAAAGAGTTGGTAGCTCTTGATCCGGCAAACAAACCACCGCTGGTAGC
GGTGGTTTTTTTGTTTGCAAGCAGCAGATTACGCGCAGAAAAAAAGGATCTCAAGAAGATCCTTTGATCTTTTCTAC
GGGGTCTGACGCTCAGTGGAACGAAAACTCACGTTAAGGGATTTTGGTCATGAGATTATCAAAAAGGATCTTCACCT
AGATCCTTTTAAATTAAAAATGAAGTTTTAAATCAATCTAAAGTATATATGAGTAAACTTGGTCTGACAGTTACCAA
TGCTTAATCAGTGAGGCACCTATCTCAGCGATCTGTCTATTTCGTTCATCCATAGTTGCCTGACTCCCCGTCGTGTA
GATAACTACGATACGGGAGGGCTTACCATCTGGCCCCAGTGCTGCAATGATACCGCGAGACCCACGCTCACCGGCTC
CAGATTTATCAGCAATAAACCAGCCAGCCGGAAGGGCCGAGCGCAGAAGTGGTCCTGCAACTTTATCCGCCTCCATC
CAGTCTATTAATTGTTGCCGGGAAGCTAGAGTAAGTAGTTCGCCAGTTAATAGTTTGCGCAACGTTGTTGCCATTGC
TACAGGCATCGTGGTGTCACGCTCGTCGTTTGGTATGGCTTCATTCAGCTCCGGTTCCCAACGATCAAGGCGAGTTA
CATGATCCCCCATGTTGTGCAAAAAAGCGGTTAGCTCCTTCGGTCCTCCGATCGTTGTCAGAAGTAAGTTGGCCGCA
GTGTTATCACTCATGGTTATGGCAGCACTGCATAATTCTCTTACTGTCATGCCATCCGTAAGATGCTTTTCTGTGAC
TGGTGAGTACTCAACCAAGTCATTCTGAGAATAGTGTATGCGGCGACCGAGTTGCTCTTGCCCGGCGTCAATACGGG
ATAATACCGCGCCACATAGCAGAACTTTAAAAGTGCTCATCATTGGAAAACGTTCTTCGGGGCGAAAACTCTCAAGG
ATCTTACCGCTGTTGAGATCCAGTTCGATGTAACCCACTCGTGCACCCAACTGATCTTCAGCATCTTTTACTTTCAC
CAGCGTTTCTGGGTGAGCAAAAACAGGAAGGCAAAATGCCGCAAAAAAGGGAATAAGGGCGACACGGAAATGTTGAA
TACTCATACTCTTCCTTTTTCAATATTATTGAAGCATTTATCAGGGTTATTGTCTCATGAGCGGATACATATTTGAA
TGTATTTAGAAAAATAAACAAATAGGGGTTCCGCGCACATTTCCCCGAAAAGTGCCACCTGACGTCTAAGAAACCAT
TATTATCATGACATTAACCTATAAAAATAGGCGTATCACGAGGCCCTTTCGTC

>PITX3 M28V; R46Q in pUC57

TCGCGCGTTTCGGTGATGACGGTGAAAACCTCTGACACATGCAGCTCCCGGAGACGGTCACAGCTTGTCTGTAAGCG
GATGCCGGGAGCAGACAAGCCCGTCAGGGCGCGTCAGCGGGTGTTGGCGGGTGTCGGGGCTGGCTTAACTATGCGGC
ATCAGAGCAGATTGTACTGAGAGTGCACCATATGCGGTGTGAAATACCGCACAGATGCGTAAGGAGAAAATACCGCA
TCAGGCGCCATTCGCCATTCAGGCTGCGCAACTGTTGGGAAGGGCGATCGGTGCGGGCCTCTTCGCTATTACGCCAG
CTGGCGAAAGGGGGATGTGCTGCAAGGCGATTAAGTTGGGTAACGCCAGGGTTTTCCCAGTCACGACGTTGTAAAAC
GACGGCCAGTGAATTCGAGCTCGGTACCTCGCGAATGCATCTAGATCATATGTCGCTGAAAAAGAAGCAGCGGCGGC
AGCGCACGCACTTCACCAGCCAGCAGCTACAGGAGCTAGAGGCGACCTTCCAGAGGAACCGCTACCCCGACGTGAGC
ACGCGCGAGGAGATCGCCGTGTGGACCAACCTCACCGAGGCCCGCGTGCAGGTGTGGTTCAAGAACCGGCGCGCCAA
ATGGCGGAAGCGCGAGCGCTAACTCGAGATCGGATCCCGGGCCCGTCGACTGCAGAGGCCTGCATGCAAGCTTGGCG
TAATCATGGTCATAGCTGTTTCCTGTGTGAAATTGTTATCCGCTCACAATTCCACACAACATACGAGCCGGAAGCAT
AAAGTGTAAAGCCTGGGGTGCCTAATGAGTGAGCTAACTCACATTAATTGCGTTGCGCTCACTGCCCGCTTTCCAGT
CGGGAAACCTGTCGTGCCAGCTGCATTAATGAATCGGCCAACGCGCGGGGAGAGGCGGTTTGCGTATTGGGCGCTCT
TCCGCTTCCTCGCTCACTGACTCGCTGCGCTCGGTCGTTCGGCTGCGGCGAGCGGTATCAGCTCACTCAAAGGCGGT
AATACGGTTATCCACAGAATCAGGGGATAACGCAGGAAAGAACATGTGAGCAAAAGGCCAGCAAAAGGCCAGGAACC
GTAAAAAGGCCGCGTTGCTGGCGTTTTTCCATAGGCTCCGCCCCCCTGACGAGCATCACAAAAATCGACGCTCAAGT
CAGAGGTGGCGAAACCCGACAGGACTATAAAGATACCAGGCGTTTCCCCCTGGAAGCTCCCTCGTGCGCTCTCCTGT
TCCGACCCTGCCGCTTACCGGATACCTGTCCGCCTTTCTCCCTTCGGGAAGCGTGGCGCTTTCTCATAGCTCACGCT
GTAGGTATCTCAGTTCGGTGTAGGTCGTTCGCTCCAAGCTGGGCTGTGTGCACGAACCCCCCGTTCAGCCCGACCGC
TGCGCCTTATCCGGTAACTATCGTCTTGAGTCCAACCCGGTAAGACACGACTTATCGCCACTGGCAGCAGCCACTGG
TAACAGGATTAGCAGAGCGAGGTATGTAGGCGGTGCTACAGAGTTCTTGAAGTGGTGGCCTAACTACGGCTACACTA
GAAGAACAGTATTTGGTATCTGCGCTCTGCTGAAGCCAGTTACCTTCGGAAAAAGAGTTGGTAGCTCTTGATCCGGC
AAACAAACCACCGCTGGTAGCGGTGGTTTTTTTGTTTGCAAGCAGCAGATTACGCGCAGAAAAAAAGGATCTCAAGA
AGATCCTTTGATCTTTTCTACGGGGTCTGACGCTCAGTGGAACGAAAACTCACGTTAAGGGATTTTGGTCATGAGAT
TATCAAAAAGGATCTTCACCTAGATCCTTTTAAATTAAAAATGAAGTTTTAAATCAATCTAAAGTATATATGAGTAA
ACTTGGTCTGACAGTTACCAATGCTTAATCAGTGAGGCACCTATCTCAGCGATCTGTCTATTTCGTTCATCCATAGT
TGCCTGACTCCCCGTCGTGTAGATAACTACGATACGGGAGGGCTTACCATCTGGCCCCAGTGCTGCAATGATACCGC
GAGACCCACGCTCACCGGCTCCAGATTTATCAGCAATAAACCAGCCAGCCGGAAGGGCCGAGCGCAGAAGTGGTCCT
GCAACTTTATCCGCCTCCATCCAGTCTATTAATTGTTGCCGGGAAGCTAGAGTAAGTAGTTCGCCAGTTAATAGTTT
GCGCAACGTTGTTGCCATTGCTACAGGCATCGTGGTGTCACGCTCGTCGTTTGGTATGGCTTCATTCAGCTCCGGTT
CCCAACGATCAAGGCGAGTTACATGATCCCCCATGTTGTGCAAAAAAGCGGTTAGCTCCTTCGGTCCTCCGATCGTT
GTCAGAAGTAAGTTGGCCGCAGTGTTATCACTCATGGTTATGGCAGCACTGCATAATTCTCTTACTGTCATGCCATC
CGTAAGATGCTTTTCTGTGACTGGTGAGTACTCAACCAAGTCATTCTGAGAATAGTGTATGCGGCGACCGAGTTGCT
CTTGCCCGGCGTCAATACGGGATAATACCGCGCCACATAGCAGAACTTTAAAAGTGCTCATCATTGGAAAACGTTCT
TCGGGGCGAAAACTCTCAAGGATCTTACCGCTGTTGAGATCCAGTTCGATGTAACCCACTCGTGCACCCAACTGATC
TTCAGCATCTTTTACTTTCACCAGCGTTTCTGGGTGAGCAAAAACAGGAAGGCAAAATGCCGCAAAAAAGGGAATAA
GGGCGACACGGAAATGTTGAATACTCATACTCTTCCTTTTTCAATATTATTGAAGCATTTATCAGGGTTATTGTCTC
ATGAGCGGATACATATTTGAATGTATTTAGAAAAATAAACAAATAGGGGTTCCGCGCACATTTCCCCGAAAAGTGCC
ACCTGACGTCTAAGAAACCATTATTATCATGACATTAACCTATAAAAATAGGCGTATCACGAGGCCCTTTCGTC

>ALX4 Q50K in ALX4_OHu22706C_pcDNA3.1(+)-N-HA

GACGGATCGGGAGATCTCCCGATCCCCTATGGTGCACTCTCAGTACAATCTGCTCTGATGCCGCATAGTTAAGCCAG
TATCTGCTCCCTGCTTGTGTGTTGGAGGTCGCTGAGTAGTGCGCGAGCAAAATTTAAGCTACAACAAGGCAAGGCTT
GACCGACAATTGCATGAAGAATCTGCTTAGGGTTAGGCGTTTTGCGCTGCTTCGCGATGTACGGGCCAGATATACGC
GTTGACATTGATTATTGACTAGTTATTAATAGTAATCAATTACGGGGTCATTAGTTCATAGCCCATATATGGAGTTC
CGCGTTACATAACTTACGGTAAATGGCCCGCCTGGCTGACCGCCCAACGACCCCCGCCCATTGACGTCAATAATGAC
GTATGTTCCCATAGTAACGCCAATAGGGACTTTCCATTGACGTCAATGGGTGGAGTATTTACGGTAAACTGCCCACT
TGGCAGTACATCAAGTGTATCATATGCCAAGTACGCCCCCTATTGACGTCAATGACGGTAAATGGCCCGCCTGGCAT
TATGCCCAGTACATGACCTTATGGGACTTTCCTACTTGGCAGTACATCTACGTATTAGTCATCGCTATTACCATGGT
GATGCGGTTTTGGCAGTACATCAATGGGCGTGGATAGCGGTTTGACTCACGGGGATTTCCAAGTCTCCACCCCATTG
ACGTCAATGGGAGTTTGTTTTGGCACCAAAATCAACGGGACTTTCCAAAATGTCGTAACAACTCCGCCCCATTGACG
CAAATGGGCGGTAGGCGTGTACGGTGGGAGGTCTATATAAGCAGAGCTCTCTGGCTAACTAGAGAACCCACTGCTTA
CTGGCTTATCGAAATTAATACGACTCACTATAGGGAGACCCAAGCTGGCTAGCGTTTAAACTTAAGCTTGCCACCAT
GTACCCATACGATGTTCCAGATTACGCTGGCTCTGGAAATGCTGAGACTTGCGTCTCTTACTGCGAGTCGCCGGCCG
CTGCCATGGACGCCTACTACAGCCCGGTGTCGCAGAGTCGGGAGGGCTCGTCGCCTTTTAGGGCATTTCCCGGAGGC
GACAAGTTCGGCACAACTTTCCTGTCGGCCGCCGCCAAAGCACAGGGATTCGGGGACGCCAAGAGCCGGGCCCGTTA
CGGCGCTGGGCAGCAGGACCTGGCGACACCCCTGGAGAGTGGAGCTGGGGCGCGGGGCTCCTTTAACAAGTTCCAGC
CCCAGCCGTCGACCCCGCAGCCCCAGCCGCCGCCGCAGCCGCAGCCGCAGCAGCAGCAGCCGCAGCCCCAGCCGCCC
GCGCAACCGCATCTTTACTTGCAGCGAGGCGCCTGCAAGACGCCCCCGGACGGCAGCCTCAAACTCCAGGAAGGCAG
CAGCGGCCACAGCGCGGCCTTGCAGGTTCCCTGCTACGCTAAAGAGAGCTCCCTGGGTGAGCCAGAGTTACCCCCTG
ACTCTGACACTGTGGGGATGGACAGCAGCTACCTGAGTGTCAAGGAGGCTGGGGTGAAGGGGCCCCAGGACCGGGCC
AGCTCAGACCTCCCCAGCCCATTGGAGAAGGCCGACTCAGAGAGCAACAAGGGCAAGAAGCGGCGGAACCGGACCAC
CTTCACCAGCTACCAGCTGGAGGAGCTGGAGAAGGTCTTCCAGAAGACCCACTACCCAGACGTGTATGCGCGGGAAC
AGCTGGCCATGAGGACAGACCTCACTGAGGCCCGCGTGCAGGTCTGGTTCAAGAACCGAAGGGCCAAGTGGAGGAAG
CGGGAGCGTTTTGGGCAGATGCAGCAGGTTCGAACCCACTTCTCCACTGCATATGAGCTGCCCCTCCTCACCCGAGC
TGAGAACTACGCCCAGATTCAGAACCCGTCCTGGCTCGGCAACAACGGGGCTGCCTCACCAGTGCCAGCCTGCGTGG
TCCCCTGCGACCCGGTGCCTGCCTGCATGTCCCCTCATGCCCACCCCCCTGGCTCTGGGGCCAGCAGCGTCACCGAC
TTCCTGAGTGTGTCTGGGGCTGGCAGTCACGTGGGCCAGACGCACATGGGCAGCCTGTTTGGAGCAGCCAGCCTCAG
CCCAGGCCTCAATGGCTACGAGCTCAACGGCGAGCCGGACCGCAAGACCTCGAGCATCGCGGCCCTCCGCATGAAGG
CCAAGGAGCACAGTGCGGCCATTTCCTGGGCCACATGAGGATCCGAATTCTGCAGATATCCAGCACAGTGGCGGCCG
CTCGAGTCTAGAGGGCCCGTTTAAACCCGCTGATCAGCCTCGACTGTGCCTTCTAGTTGCCAGCCATCTGTTGTTTG
CCCCTCCCCCGTGCCTTCCTTGACCCTGGAAGGTGCCACTCCCACTGTCCTTTCCTAATAAAATGAGGAAATTGCAT
CGCATTGTCTGAGTAGGTGTCATTCTATTCTGGGGGGTGGGGTGGGGCAGGACAGCAAGGGGGAGGATTGGGAAGAC
AATAGCAGGCATGCTGGGGATGCGGTGGGCTCTATGGCTTCTGAGGCGGAAAGAACCAGCTGGGGCTCTAGGGGGTA
TCCCCACGCGCCCTGTAGCGGCGCATTAAGCGCGGCGGGTGTGGTGGTTACGCGCAGCGTGACCGCTACACTTGCCA
GCGCCCTAGCGCCCGCTCCTTTCGCTTTCTTCCCTTCCTTTCTCGCCACGTTCGCCGGCTTTCCCCGTCAAGCTCTA
AATCGGGGGCTCCCTTTAGGGTTCCGATTTAGTGCTTTACGGCACCTCGACCCCAAAAAACTTGATTAGGGTGATGG
TTCACGTAGTGGGCCATCGCCCTGATAGACGGTTTTTCGCCCTTTGACGTTGGAGTCCACGTTCTTTAATAGTGGAC
TCTTGTTCCAAACTGGAACAACACTCAACCCTATCTCGGTCTATTCTTTTGATTTATAAGGGATTTTGCCGATTTCG
GCCTATTGGTTAAAAAATGAGCTGATTTAACAAAAATTTAACGCGAATTAATTCTGTGGAATGTGTGTCAGTTAGGG
TGTGGAAAGTCCCCAGGCTCCCCAGCAGGCAGAAGTATGCAAAGCATGCATCTCAATTAGTCAGCAACCAGGTGTGG
AAAGTCCCCAGGCTCCCCAGCAGGCAGAAGTATGCAAAGCATGCATCTCAATTAGTCAGCAACCATAGTCCCGCCCC
TAACTCCGCCCATCCCGCCCCTAACTCCGCCCAGTTCCGCCCATTCTCCGCCCCATGGCTGACTAATTTTTTTTATT

TATGCAGAGGCCGAGGCCGCCTCTGCCTCTGAGCTATTCCAGAAGTAGTGAGGAGGCTTTTTTGGAGGCCTAGGCTT
TTGCAAAAAGCTCCCGGGAGCTTGTATATCCATTTTCGGATCTGATCAAGAGACAGGATGAGGATCGTTTCGCATGA
TTGAACAAGATGGATTGCACGCAGGTTCTCCGGCCGCTTGGGTGGAGAGGCTATTCGGCTATGACTGGGCACAACAG
ACAATCGGCTGCTCTGATGCCGCCGTGTTCCGGCTGTCAGCGCAGGGGCGCCCGGTTCTTTTTGTCAAGACCGACCT
GTCCGGTGCCCTGAATGAACTGCAGGACGAGGCAGCGCGGCTATCGTGGCTGGCCACGACGGGCGTTCCTTGCGCAG
CTGTGCTCGACGTTGTCACTGAAGCGGGAAGGGACTGGCTGCTATTGGGCGAAGTGCCGGGGCAGGATCTCCTGTCA
TCTCACCTTGCTCCTGCCGAGAAAGTATCCATCATGGCTGATGCAATGCGGCGGCTGCATACGCTTGATCCGGCTAC
CTGCCCATTCGACCACCAAGCGAAACATCGCATCGAGCGAGCACGTACTCGGATGGAAGCCGGTCTTGTCGATCAGG
ATGATCTGGACGAAGAGCATCAGGGGCTCGCGCCAGCCGAACTGTTCGCCAGGCTCAAGGCGCGCATGCCCGACGGC
GAGGATCTCGTCGTGACCCATGGCGATGCCTGCTTGCCGAATATCATGGTGGAAAATGGCCGCTTTTCTGGATTCAT
CGACTGTGGCCGGCTGGGTGTGGCGGACCGCTATCAGGACATAGCGTTGGCTACCCGTGATATTGCTGAAGAGCTTG
GCGGCGAATGGGCTGACCGCTTCCTCGTGCTTTACGGTATCGCCGCTCCCGATTCGCAGCGCATCGCCTTCTATCGC
CTTCTTGACGAGTTCTTCTGAGCGGGACTCTGGGGTTCGAAATGACCGACCAAGCGACGCCCAACCTGCCATCACGA
GATTTCGATTCCACCGCCGCCTTCTATGAAAGGTTGGGCTTCGGAATCGTTTTCCGGGACGCCGGCTGGATGATCCT
CCAGCGCGGGGATCTCATGCTGGAGTTCTTCGCCCACCCCAACTTGTTTATTGCAGCTTATAATGGTTACAAATAAA
GCAATAGCATCACAAATTTCACAAATAAAGCATTTTTTTCACTGCATTCTAGTTGTGGTTTGTCCAAACTCATCAAT
GTATCTTATCATGTCTGTATACCGTCGACCTCTAGCTAGAGCTTGGCGTAATCATGGTCATAGCTGTTTCCTGTGTG
AAATTGTTATCCGCTCACAATTCCACACAACATACGAGCCGGAAGCATAAAGTGTAAAGCCTGGGGTGCCTAATGAG
TGAGCTAACTCACATTAATTGCGTTGCGCTCACTGCCCGCTTTCCAGTCGGGAAACCTGTCGTGCCAGCTGCATTAA
TGAATCGGCCAACGCGCGGGGAGAGGCGGTTTGCGTATTGGGCGCTCTTCCGCTTCCTCGCTCACTGACTCGCTGCG
CTCGGTCGTTCGGCTGCGGCGAGCGGTATCAGCTCACTCAAAGGCGGTAATACGGTTATCCACAGAATCAGGGGATA
ACGCAGGAAAGAACATGTGAGCAAAAGGCCAGCAAAAGGCCAGGAACCGTAAAAAGGCCGCGTTGCTGGCGTTTTTC
CATAGGCTCCGCCCCCCTGACGAGCATCACAAAAATCGACGCTCAAGTCAGAGGTGGCGAAACCCGACAGGACTATA
AAGATACCAGGCGTTTCCCCCTGGAAGCTCCCTCGTGCGCTCTCCTGTTCCGACCCTGCCGCTTACCGGATACCTGT
CCGCCTTTCTCCCTTCGGGAAGCGTGGCGCTTTCTCATAGCTCACGCTGTAGGTATCTCAGTTCGGTGTAGGTCGTT
CGCTCCAAGCTGGGCTGTGTGCACGAACCCCCCGTTCAGCCCGACCGCTGCGCCTTATCCGGTAACTATCGTCTTGA
GTCCAACCCGGTAAGACACGACTTATCGCCACTGGCAGCAGCCACTGGTAACAGGATTAGCAGAGCGAGGTATGTAG
GCGGTGCTACAGAGTTCTTGAAGTGGTGGCCTAACTACGGCTACACTAGAAGAACAGTATTTGGTATCTGCGCTCTG
CTGAAGCCAGTTACCTTCGGAAAAAGAGTTGGTAGCTCTTGATCCGGCAAACAAACCACCGCTGGTAGCGGTGGTTT
TTTTGTTTGCAAGCAGCAGATTACGCGCAGAAAAAAAGGATCTCAAGAAGATCCTTTGATCTTTTCTACGGGGTCTG
ACGCTCAGTGGAACGAAAACTCACGTTAAGGGATTTTGGTCATGAGATTATCAAAAAGGATCTTCACCTAGATCCTT
TTAAATTAAAAATGAAGTTTTAAATCAATCTAAAGTATATATGAGTAAACTTGGTCTGACAGTTACCAATGCTTAAT
CAGTGAGGCACCTATCTCAGCGATCTGTCTATTTCGTTCATCCATAGTTGCCTGACTCCCCGTCGTGTAGATAACTA
CGATACGGGAGGGCTTACCATCTGGCCCCAGTGCTGCAATGATACCGCGAGACCCACGCTCACCGGCTCCAGATTTA
TCAGCAATAAACCAGCCAGCCGGAAGGGCCGAGCGCAGAAGTGGTCCTGCAACTTTATCCGCCTCCATCCAGTCTAT
TAATTGTTGCCGGGAAGCTAGAGTAAGTAGTTCGCCAGTTAATAGTTTGCGCAACGTTGTTGCCATTGCTACAGGCA
TCGTGGTGTCACGCTCGTCGTTTGGTATGGCTTCATTCAGCTCCGGTTCCCAACGATCAAGGCGAGTTACATGATCC
CCCATGTTGTGCAAAAAAGCGGTTAGCTCCTTCGGTCCTCCGATCGTTGTCAGAAGTAAGTTGGCCGCAGTGTTATC
ACTCATGGTTATGGCAGCACTGCATAATTCTCTTACTGTCATGCCATCCGTAAGATGCTTTTCTGTGACTGGTGAGT
ACTCAACCAAGTCATTCTGAGAATAGTGTATGCGGCGACCGAGTTGCTCTTGCCCGGCGTCAATACGGGATAATACC
GCGCCACATAGCAGAACTTTAAAAGTGCTCATCATTGGAAAACGTTCTTCGGGGCGAAAACTCTCAAGGATCTTACC
GCTGTTGAGATCCAGTTCGATGTAACCCACTCGTGCACCCAACTGATCTTCAGCATCTTTTACTTTCACCAGCGTTT
CTGGGTGAGCAAAAACAGGAAGGCAAAATGCCGCAAAAAAGGGAATAAGGGCGACACGGAAATGTTGAATACTCATA

CTCTTCCTTTTTCAATATTATTGAAGCATTTATCAGGGTTATTGTCTCATGAGCGGATACATATTTGAATGTATTTA
GAAAAATAAACAAATAGGGGTTCCGCGCACATTTCCCCGAAAAGTGCCACCTGACGTC

>CRX K50Q in CRX_OHu18946C_pcDNA3.1(+)-N-HA

GACGGATCGGGAGATCTCCCGATCCCCTATGGTGCACTCTCAGTACAATCTGCTCTGATGCCGCATAGTTAAGCCAG
TATCTGCTCCCTGCTTGTGTGTTGGAGGTCGCTGAGTAGTGCGCGAGCAAAATTTAAGCTACAACAAGGCAAGGCTT
GACCGACAATTGCATGAAGAATCTGCTTAGGGTTAGGCGTTTTGCGCTGCTTCGCGATGTACGGGCCAGATATACGC
GTTGACATTGATTATTGACTAGTTATTAATAGTAATCAATTACGGGGTCATTAGTTCATAGCCCATATATGGAGTTC
CGCGTTACATAACTTACGGTAAATGGCCCGCCTGGCTGACCGCCCAACGACCCCCGCCCATTGACGTCAATAATGAC
GTATGTTCCCATAGTAACGCCAATAGGGACTTTCCATTGACGTCAATGGGTGGAGTATTTACGGTAAACTGCCCACT
TGGCAGTACATCAAGTGTATCATATGCCAAGTACGCCCCCTATTGACGTCAATGACGGTAAATGGCCCGCCTGGCAT
TATGCCCAGTACATGACCTTATGGGACTTTCCTACTTGGCAGTACATCTACGTATTAGTCATCGCTATTACCATGGT
GATGCGGTTTTGGCAGTACATCAATGGGCGTGGATAGCGGTTTGACTCACGGGGATTTCCAAGTCTCCACCCCATTG
ACGTCAATGGGAGTTTGTTTTGGCACCAAAATCAACGGGACTTTCCAAAATGTCGTAACAACTCCGCCCCATTGACG
CAAATGGGCGGTAGGCGTGTACGGTGGGAGGTCTATATAAGCAGAGCTCTCTGGCTAACTAGAGAACCCACTGCTTA
CTGGCTTATCGAAATTAATACGACTCACTATAGGGAGACCCAAGCTGGCTAGCGTTTAAACTTAAGCTTGCCACCAT
GTACCCATACGATGTTCCAGATTACGCTGGCTCTGGAATGGCGTATATGAACCCGGGGCCCCACTATTCTGTCAACG
CCTTGGCCCTAAGTGGCCCCAGTGTGGATCTGATGCACCAGGCTGTGCCCTACCCAAGCGCCCCCAGGAAGCAGCGG
CGGGAGCGCACCACCTTCACCCGGAGCCAACTGGAGGAGCTGGAGGCACTGTTTGCCAAGACCCAGTACCCAGACGT
CTATGCCCGTGAGGAGGTGGCTCTGAAGATCAATCTGCCTGAGTCCAGGGTTCAGGTTTGGTTCCAGAACCGGAGGG
CTAAATGCAGGCAGCAGCGACAGCAGCAGAAACAGCAGCAGCAGCCCCCAGGGGGCCAGGCCAAGGCCCGGCCTGCC
AAGAGGAAGGCGGGCACGTCCCCAAGACCCTCCACAGATGTGTGTCCAGACCCTCTGGGCATCTCAGATTCCTACAG
TCCCCCTCTGCCCGGCCCCTCAGGCTCCCCAACCACGGCAGTGGCCACTGTGTCCATCTGGAGCCCAGCCTCAGAGT
CCCCTTTGCCTGAGGCGCAGCGGGCTGGGCTGGTGGCCTCAGGGCCGTCTCTGACCTCCGCCCCCTATGCCATGACC
TACGCCCCGGCCTCCGCTTTCTGCTCTTCCCCCTCCGCCTATGGGTCTCCGAGCTCCTATTTCAGCGGCCTAGACCC
CTACCTTTCTCCCATGGTGCCCCAGCTAGGGGGCCCGGCTCTTAGCCCCCTCTCTGGCCCCTCCGTGGGACCTTCCC
TGGCCCAGTCCCCCACCTCCCTATCAGGCCAGAGCTATGGCGCCTACAGCCCCGTGGATAGCTTGGAATTCAAGGAC
CCCACGGGCACCTGGAAATTCACCTACAATCCCATGGACCCTCTGGACTACAAGGATCAGAGTGCCTGGAAGTTTCA
GATCTTGTGATAAACCCGCTGATCAGCCTCGACTGTGCCTTCTAGTTGCCAGCCATCTGTTGTTTGCCCCTCCCCCG
TGCCTTCCTTGACCCTGGAAGGTGCCACTCCCACTGTCCTTTCCTAATAAAATGAGGAAATTGCATCGCATTGTCTG
AGTAGGTGTCATTCTATTCTGGGGGGTGGGGTGGGGCAGGACAGCAAGGGGGAGGATTGGGAAGACAATAGCAGGCA
TGCTGGGGATGCGGTGGGCTCTATGGCTTCTGAGGCGGAAAGAACCAGCTGGGGCTCTAGGGGGTATCCCCACGCGC
CCTGTAGCGGCGCATTAAGCGCGGCGGGTGTGGTGGTTACGCGCAGCGTGACCGCTACACTTGCCAGCGCCCTAGCG
CCCGCTCCTTTCGCTTTCTTCCCTTCCTTTCTCGCCACGTTCGCCGGCTTTCCCCGTCAAGCTCTAAATCGGGGGCT
CCCTTTAGGGTTCCGATTTAGTGCTTTACGGCACCTCGACCCCAAAAAACTTGATTAGGGTGATGGTTCACGTAGTG
GGCCATCGCCCTGATAGACGGTTTTTCGCCCTTTGACGTTGGAGTCCACGTTCTTTAATAGTGGACTCTTGTTCCAA
ACTGGAACAACACTCAACCCTATCTCGGTCTATTCTTTTGATTTATAAGGGATTTTGCCGATTTCGGCCTATTGGTT
AAAAAATGAGCTGATTTAACAAAAATTTAACGCGAATTAATTCTGTGGAATGTGTGTCAGTTAGGGTGTGGAAAGTC
CCCAGGCTCCCCAGCAGGCAGAAGTATGCAAAGCATGCATCTCAATTAGTCAGCAACCAGGTGTGGAAAGTCCCCAG
GCTCCCCAGCAGGCAGAAGTATGCAAAGCATGCATCTCAATTAGTCAGCAACCATAGTCCCGCCCCTAACTCCGCCC
ATCCCGCCCCTAACTCCGCCCAGTTCCGCCCATTCTCCGCCCCATGGCTGACTAATTTTTTTTATTTATGCAGAGGC
CGAGGCCGCCTCTGCCTCTGAGCTATTCCAGAAGTAGTGAGGAGGCTTTTTTGGAGGCCTAGGCTTTTGCAAAAAGC
TCCCGGGAGCTTGTATATCCATTTTCGGATCTGATCAAGAGACAGGATGAGGATCGTTTCGCATGATTGAACAAGAT
GGATTGCACGCAGGTTCTCCGGCCGCTTGGGTGGAGAGGCTATTCGGCTATGACTGGGCACAACAGACAATCGGCTG
CTCTGATGCCGCCGTGTTCCGGCTGTCAGCGCAGGGGCGCCCGGTTCTTTTTGTCAAGACCGACCTGTCCGGTGCCC
TGAATGAACTGCAGGACGAGGCAGCGCGGCTATCGTGGCTGGCCACGACGGGCGTTCCTTGCGCAGCTGTGCTCGAC

GTTGTCACTGAAGCGGGAAGGGACTGGCTGCTATTGGGCGAAGTGCCGGGGCAGGATCTCCTGTCATCTCACCTTGC
TCCTGCCGAGAAAGTATCCATCATGGCTGATGCAATGCGGCGGCTGCATACGCTTGATCCGGCTACCTGCCCATTCG
ACCACCAAGCGAAACATCGCATCGAGCGAGCACGTACTCGGATGGAAGCCGGTCTTGTCGATCAGGATGATCTGGAC
GAAGAGCATCAGGGGCTCGCGCCAGCCGAACTGTTCGCCAGGCTCAAGGCGCGCATGCCCGACGGCGAGGATCTCGT
CGTGACCCATGGCGATGCCTGCTTGCCGAATATCATGGTGGAAAATGGCCGCTTTTCTGGATTCATCGACTGTGGCC
GGCTGGGTGTGGCGGACCGCTATCAGGACATAGCGTTGGCTACCCGTGATATTGCTGAAGAGCTTGGCGGCGAATGG
GCTGACCGCTTCCTCGTGCTTTACGGTATCGCCGCTCCCGATTCGCAGCGCATCGCCTTCTATCGCCTTCTTGACGA
GTTCTTCTGAGCGGGACTCTGGGGTTCGAAATGACCGACCAAGCGACGCCCAACCTGCCATCACGAGATTTCGATTC
CACCGCCGCCTTCTATGAAAGGTTGGGCTTCGGAATCGTTTTCCGGGACGCCGGCTGGATGATCCTCCAGCGCGGGG
ATCTCATGCTGGAGTTCTTCGCCCACCCCAACTTGTTTATTGCAGCTTATAATGGTTACAAATAAAGCAATAGCATC
ACAAATTTCACAAATAAAGCATTTTTTTCACTGCATTCTAGTTGTGGTTTGTCCAAACTCATCAATGTATCTTATCA
TGTCTGTATACCGTCGACCTCTAGCTAGAGCTTGGCGTAATCATGGTCATAGCTGTTTCCTGTGTGAAATTGTTATC
CGCTCACAATTCCACACAACATACGAGCCGGAAGCATAAAGTGTAAAGCCTGGGGTGCCTAATGAGTGAGCTAACTC
ACATTAATTGCGTTGCGCTCACTGCCCGCTTTCCAGTCGGGAAACCTGTCGTGCCAGCTGCATTAATGAATCGGCCA
ACGCGCGGGGAGAGGCGGTTTGCGTATTGGGCGCTCTTCCGCTTCCTCGCTCACTGACTCGCTGCGCTCGGTCGTTC
GGCTGCGGCGAGCGGTATCAGCTCACTCAAAGGCGGTAATACGGTTATCCACAGAATCAGGGGATAACGCAGGAAAG
AACATGTGAGCAAAAGGCCAGCAAAAGGCCAGGAACCGTAAAAAGGCCGCGTTGCTGGCGTTTTTCCATAGGCTCCG
CCCCCCTGACGAGCATCACAAAAATCGACGCTCAAGTCAGAGGTGGCGAAACCCGACAGGACTATAAAGATACCAGG
CGTTTCCCCCTGGAAGCTCCCTCGTGCGCTCTCCTGTTCCGACCCTGCCGCTTACCGGATACCTGTCCGCCTTTCTC
CCTTCGGGAAGCGTGGCGCTTTCTCATAGCTCACGCTGTAGGTATCTCAGTTCGGTGTAGGTCGTTCGCTCCAAGCT
GGGCTGTGTGCACGAACCCCCCGTTCAGCCCGACCGCTGCGCCTTATCCGGTAACTATCGTCTTGAGTCCAACCCGG
TAAGACACGACTTATCGCCACTGGCAGCAGCCACTGGTAACAGGATTAGCAGAGCGAGGTATGTAGGCGGTGCTACA
GAGTTCTTGAAGTGGTGGCCTAACTACGGCTACACTAGAAGAACAGTATTTGGTATCTGCGCTCTGCTGAAGCCAGT
TACCTTCGGAAAAAGAGTTGGTAGCTCTTGATCCGGCAAACAAACCACCGCTGGTAGCGGTGGTTTTTTTGTTTGCA
AGCAGCAGATTACGCGCAGAAAAAAAGGATCTCAAGAAGATCCTTTGATCTTTTCTACGGGGTCTGACGCTCAGTGG
AACGAAAACTCACGTTAAGGGATTTTGGTCATGAGATTATCAAAAAGGATCTTCACCTAGATCCTTTTAAATTAAAA
ATGAAGTTTTAAATCAATCTAAAGTATATATGAGTAAACTTGGTCTGACAGTTACCAATGCTTAATCAGTGAGGCAC
CTATCTCAGCGATCTGTCTATTTCGTTCATCCATAGTTGCCTGACTCCCCGTCGTGTAGATAACTACGATACGGGAG
GGCTTACCATCTGGCCCCAGTGCTGCAATGATACCGCGAGACCCACGCTCACCGGCTCCAGATTTATCAGCAATAAA
CCAGCCAGCCGGAAGGGCCGAGCGCAGAAGTGGTCCTGCAACTTTATCCGCCTCCATCCAGTCTATTAATTGTTGCC
GGGAAGCTAGAGTAAGTAGTTCGCCAGTTAATAGTTTGCGCAACGTTGTTGCCATTGCTACAGGCATCGTGGTGTCA
CGCTCGTCGTTTGGTATGGCTTCATTCAGCTCCGGTTCCCAACGATCAAGGCGAGTTACATGATCCCCCATGTTGTG
CAAAAAAGCGGTTAGCTCCTTCGGTCCTCCGATCGTTGTCAGAAGTAAGTTGGCCGCAGTGTTATCACTCATGGTTA
TGGCAGCACTGCATAATTCTCTTACTGTCATGCCATCCGTAAGATGCTTTTCTGTGACTGGTGAGTACTCAACCAAG
TCATTCTGAGAATAGTGTATGCGGCGACCGAGTTGCTCTTGCCCGGCGTCAATACGGGATAATACCGCGCCACATAG
CAGAACTTTAAAAGTGCTCATCATTGGAAAACGTTCTTCGGGGCGAAAACTCTCAAGGATCTTACCGCTGTTGAGAT
CCAGTTCGATGTAACCCACTCGTGCACCCAACTGATCTTCAGCATCTTTTACTTTCACCAGCGTTTCTGGGTGAGCA
AAAACAGGAAGGCAAAATGCCGCAAAAAAGGGAATAAGGGCGACACGGAAATGTTGAATACTCATACTCTTCCTTTT
TCAATATTATTGAAGCATTTATCAGGGTTATTGTCTCATGAGCGGATACATATTTGAATGTATTTAGAAAAATAAAC
AAATAGGGGTTCCGCGCACATTTCCCCGAAAAGTGCCACCTGACGTC

>ARX P26L in Arx_OMu17806D_pcDNA3.1+/C-(K)-DYK

GACGGATCGGGAGATCTCCCGATCCCCTATGGTGCACTCTCAGTACAATCTGCTCTGATGCCGCATAGTTAAGCCAG
TATCTGCTCCCTGCTTGTGTGTTGGAGGTCGCTGAGTAGTGCGCGAGCAAAATTTAAGCTACAACAAGGCAAGGCTT
GACCGACAATTGCATGAAGAATCTGCTTAGGGTTAGGCGTTTTGCGCTGCTTCGCGATGTACGGGCCAGATATACGC
GTTGACATTGATTATTGACTAGTTATTAATAGTAATCAATTACGGGGTCATTAGTTCATAGCCCATATATGGAGTTC
CGCGTTACATAACTTACGGTAAATGGCCCGCCTGGCTGACCGCCCAACGACCCCCGCCCATTGACGTCAATAATGAC
GTATGTTCCCATAGTAACGCCAATAGGGACTTTCCATTGACGTCAATGGGTGGAGTATTTACGGTAAACTGCCCACT
TGGCAGTACATCAAGTGTATCATATGCCAAGTACGCCCCCTATTGACGTCAATGACGGTAAATGGCCCGCCTGGCAT
TATGCCCAGTACATGACCTTATGGGACTTTCCTACTTGGCAGTACATCTACGTATTAGTCATCGCTATTACCATGGT
GATGCGGTTTTGGCAGTACATCAATGGGCGTGGATAGCGGTTTGACTCACGGGGATTTCCAAGTCTCCACCCCATTG
ACGTCAATGGGAGTTTGTTTTGGCACCAAAATCAACGGGACTTTCCAAAATGTCGTAACAACTCCGCCCCATTGACG
CAAATGGGCGGTAGGCGTGTACGGTGGGAGGTCTATATAAGCAGAGCTCTCTGGCTAACTAGAGAACCCACTGCTTA
CTGGCTTATCGAAATTAATACGACTCACTATAGGGAGACCCAAGCTGGCTAGCGTTTAAACTTAAGCTTGGTACCGA
GCTCGGATCCGCCACCATGAGCAATCAGTACCAGGAAGAGGGCTGCTCTGAGAGGCCGGAGTGCAAGAGTAAATCTC
CAACTTTGCTCTCCTCCTACTGCATCGACAGCATCCTGGGCCGGAGAAGCCCATGCAAAATGAGGCTTCTGGGAGCC
GCGCAGAGCTTGCCTGCCCCGCTGGCCAGCCGCGCTGACCAGGAAAAAGCCATGCAAGGCTCCCCCAAGAGCAGCAG
CGCCCCGTTCGAGGCTGAGCTGCACCTGCCACCCAAGCTGCGGCGCCTCTACGGCCCGGGCGGGGGCCGCCTCCTCC
AGGGCGCGGCGGCGGCGGCTGCAGCGGCGGCGGCGGCCGCAGCGGCGGCCACGGCCACTGGCACCGCAGGTCCCCGT
GGAGAGGTTCCTCCACCGCCACCACCAGCTGCACGGCCAGGGGAGCGTCAGGACAGCGCAGGGGCCGTGGCTGCCGC
CGCTGCCGCCGCTGCCTGGGACACGCTCAAGATCAGCCAGGCGCCGCAGGTGAGCATCAGTCGTAGCAAGTCGTACC
GCGAGAACGGGGCGCCCTTCGTGCCCCCGCCGCCAGCCCTGGACGAGCTGAGTGGCCCGGGGGGCGTCGCGCACCCC
GAGGAGCGCCTCAGCGCGGCCAGCGGACCCGGCAGTGCCCCCGCGGCGGGTGGTGGCACGGGCGCCGAGGACGACGA
GGAGGAGCTGTTGGAGGATGAGGAAGATGAAGAGGAGGAAGAGGAGCTGCTGGAGGACGACGACGAGGAGCTGCTGG
AGGACGATGCCCGCGCGCTGCTCAAGGAGCCCCGGCGCTGCTCGGTGGCCACCACCGGCACAGTGGCCGCCGCCGCC
GCCGCTGCTGCCGCCGCCGTGGCCACCGAGGGCGGGGAGCTGTCGCCCAAGGAAGAGCTGTTGCTGCACCCGGAGGA
CGCCGAGGGCAAGGATGGTGAGGACAGCGTGTGCCTGTCGGCCGGCAGCGACTCGGAGGAGGGGCTGCTGAAGCGCA
AACAGAGGCGCTACCGCACCACGTTCACCAGTTACCAGCTGGAGGAACTGGAGCGGGCTTTCCAGAAGACGCACTAC
CTTGACGTCTTCACCAGGGAGGAGCTGGCCATGAGGCTGGACCTGACAGAGGCCCGAGTCCAGGTGTGGTTCCAGAA
CCGTCGGGCCAAGTGGCGCAAGCGGGAGAAGGCTGGCGCACAGACCCACCCCCCGGGGCTGCCCTTTCCAGGGCCGC
TCTCGGCTACCCACCCTCTCAGCCCCTACCTGGACGCCAGCCCCTTTCCCCCGCATCACCCTGCGCTTGATTCGGCC
TGGACCGCCGCGGCGGCGGCGGCTGCAGCTGCCTTCCCGAGCCTACCTCCGCCCCCAGGCTCTGCTAGCCTGCCACC
CAGCGGGGCGCCGCTGGGTCTGAGCACTTTTCTAGGAGCAGCGGTGTTCCGCCACCCAGCCTTCATCAGCCCAGCAT
TTGGCAGGCTCTTTTCCACTATGGCCCCCCTGACCAGCGCGTCGACTGCGGCTGCGCTCCTGAGACAGCCCACACCC
GCGGTGGAGGGCGCAGTGGCCTCGGGCGCGCTGGCCGACCCGGCCACCGCCGCGGCAGACAGACGCGCGTCAAGCAT
AGCCGCGCTGAGGCTCAAGGCTAAGGAGCACGCCGCGCAGCTCACGCAGCTCAACATCCTGCCGGGCACCAGCACGG
GGAAGGAGGTGTGCGATTACAAGGATGACGACGATAAGTGATAAACCCGCTGATCAGCCTCGACTGTGCCTTCTAGT
TGCCAGCCATCTGTTGTTTGCCCCTCCCCCGTGCCTTCCTTGACCCTGGAAGGTGCCACTCCCACTGTCCTTTCCTA
ATAAAATGAGGAAATTGCATCGCATTGTCTGAGTAGGTGTCATTCTATTCTGGGGGGTGGGGTGGGGCAGGACAGCA
AGGGGGAGGATTGGGAAGACAATAGCAGGCATGCTGGGGATGCGGTGGGCTCTATGGCTTCTGAGGCGGAAAGAACC
AGCTGGGGCTCTAGGGGGTATCCCCACGCGCCCTGTAGCGGCGCATTAAGCGCGGCGGGTGTGGTGGTTACGCGCAG
CGTGACCGCTACACTTGCCAGCGCCCTAGCGCCCGCTCCTTTCGCTTTCTTCCCTTCCTTTCTCGCCACGTTCGCCG
GCTTTCCCCGTCAAGCTCTAAATCGGGGGCTCCCTTTAGGGTTCCGATTTAGTGCTTTACGGCACCTCGACCCCAAA
AAACTTGATTAGGGTGATGGTTCACGTAGTGGGCCATCGCCCTGATAGACGGTTTTTCGCCCTTTGACGTTGGAGTC

CACGTTCTTTAATAGTGGACTCTTGTTCCAAACTGGAACAACACTCAACCCTATCTCGGTCTATTCTTTTGATTTAT
AAGGGATTTTGCCGATTTCGGCCTATTGGTTAAAAAATGAGCTGATTTAACAAAAATTTAACGCGAATTAATTCTGT
GGAATGTGTGTCAGTTAGGGTGTGGAAAGTCCCCAGGCTCCCCAGCAGGCAGAAGTATGCAAAGCATGCATCTCAAT
TAGTCAGCAACCAGGTGTGGAAAGTCCCCAGGCTCCCCAGCAGGCAGAAGTATGCAAAGCATGCATCTCAATTAGTC
AGCAACCATAGTCCCGCCCCTAACTCCGCCCATCCCGCCCCTAACTCCGCCCAGTTCCGCCCATTCTCCGCCCCATG
GCTGACTAATTTTTTTTATTTATGCAGAGGCCGAGGCCGCCTCTGCCTCTGAGCTATTCCAGAAGTAGTGAGGAGGC
TTTTTTGGAGGCCTAGGCTTTTGCAAAAAGCTCCCGGGAGCTTGTATATCCATTTTCGGATCTGATCAAGAGACAGG
ATGAGGATCGTTTCGCATGATTGAACAAGATGGATTGCACGCAGGTTCTCCGGCCGCTTGGGTGGAGAGGCTATTCG
GCTATGACTGGGCACAACAGACAATCGGCTGCTCTGATGCCGCCGTGTTCCGGCTGTCAGCGCAGGGGCGCCCGGTT
CTTTTTGTCAAGACCGACCTGTCCGGTGCCCTGAATGAACTGCAGGACGAGGCAGCGCGGCTATCGTGGCTGGCCAC
GACGGGCGTTCCTTGCGCAGCTGTGCTCGACGTTGTCACTGAAGCGGGAAGGGACTGGCTGCTATTGGGCGAAGTGC
CGGGGCAGGATCTCCTGTCATCTCACCTTGCTCCTGCCGAGAAAGTATCCATCATGGCTGATGCAATGCGGCGGCTG
CATACGCTTGATCCGGCTACCTGCCCATTCGACCACCAAGCGAAACATCGCATCGAGCGAGCACGTACTCGGATGGA
AGCCGGTCTTGTCGATCAGGATGATCTGGACGAAGAGCATCAGGGGCTCGCGCCAGCCGAACTGTTCGCCAGGCTCA
AGGCGCGCATGCCCGACGGCGAGGATCTCGTCGTGACCCATGGCGATGCCTGCTTGCCGAATATCATGGTGGAAAAT
GGCCGCTTTTCTGGATTCATCGACTGTGGCCGGCTGGGTGTGGCGGACCGCTATCAGGACATAGCGTTGGCTACCCG
TGATATTGCTGAAGAGCTTGGCGGCGAATGGGCTGACCGCTTCCTCGTGCTTTACGGTATCGCCGCTCCCGATTCGC
AGCGCATCGCCTTCTATCGCCTTCTTGACGAGTTCTTCTGAGCGGGACTCTGGGGTTCGAAATGACCGACCAAGCGA
CGCCCAACCTGCCATCACGAGATTTCGATTCCACCGCCGCCTTCTATGAAAGGTTGGGCTTCGGAATCGTTTTCCGG
GACGCCGGCTGGATGATCCTCCAGCGCGGGGATCTCATGCTGGAGTTCTTCGCCCACCCCAACTTGTTTATTGCAGC
TTATAATGGTTACAAATAAAGCAATAGCATCACAAATTTCACAAATAAAGCATTTTTTTCACTGCATTCTAGTTGTG
GTTTGTCCAAACTCATCAATGTATCTTATCATGTCTGTATACCGTCGACCTCTAGCTAGAGCTTGGCGTAATCATGG
TCATAGCTGTTTCCTGTGTGAAATTGTTATCCGCTCACAATTCCACACAACATACGAGCCGGAAGCATAAAGTGTAA
AGCCTGGGGTGCCTAATGAGTGAGCTAACTCACATTAATTGCGTTGCGCTCACTGCCCGCTTTCCAGTCGGGAAACC
TGTCGTGCCAGCTGCATTAATGAATCGGCCAACGCGCGGGGAGAGGCGGTTTGCGTATTGGGCGCTCTTCCGCTTCC
TCGCTCACTGACTCGCTGCGCTCGGTCGTTCGGCTGCGGCGAGCGGTATCAGCTCACTCAAAGGCGGTAATACGGTT
ATCCACAGAATCAGGGGATAACGCAGGAAAGAACATGTGAGCAAAAGGCCAGCAAAAGGCCAGGAACCGTAAAAAGG
CCGCGTTGCTGGCGTTTTTCCATAGGCTCCGCCCCCCTGACGAGCATCACAAAAATCGACGCTCAAGTCAGAGGTGG
CGAAACCCGACAGGACTATAAAGATACCAGGCGTTTCCCCCTGGAAGCTCCCTCGTGCGCTCTCCTGTTCCGACCCT
GCCGCTTACCGGATACCTGTCCGCCTTTCTCCCTTCGGGAAGCGTGGCGCTTTCTCATAGCTCACGCTGTAGGTATC
TCAGTTCGGTGTAGGTCGTTCGCTCCAAGCTGGGCTGTGTGCACGAACCCCCCGTTCAGCCCGACCGCTGCGCCTTA
TCCGGTAACTATCGTCTTGAGTCCAACCCGGTAAGACACGACTTATCGCCACTGGCAGCAGCCACTGGTAACAGGAT
TAGCAGAGCGAGGTATGTAGGCGGTGCTACAGAGTTCTTGAAGTGGTGGCCTAACTACGGCTACACTAGAAGAACAG
TATTTGGTATCTGCGCTCTGCTGAAGCCAGTTACCTTCGGAAAAAGAGTTGGTAGCTCTTGATCCGGCAAACAAACC
ACCGCTGGTAGCGGTGGTTTTTTTGTTTGCAAGCAGCAGATTACGCGCAGAAAAAAAGGATCTCAAGAAGATCCTTT
GATCTTTTCTACGGGGTCTGACGCTCAGTGGAACGAAAACTCACGTTAAGGGATTTTGGTCATGAGATTATCAAAAA
GGATCTTCACCTAGATCCTTTTAAATTAAAAATGAAGTTTTAAATCAATCTAAAGTATATATGAGTAAACTTGGTCT
GACAGTTACCAATGCTTAATCAGTGAGGCACCTATCTCAGCGATCTGTCTATTTCGTTCATCCATAGTTGCCTGACT
CCCCGTCGTGTAGATAACTACGATACGGGAGGGCTTACCATCTGGCCCCAGTGCTGCAATGATACCGCGAGACCCAC
GCTCACCGGCTCCAGATTTATCAGCAATAAACCAGCCAGCCGGAAGGGCCGAGCGCAGAAGTGGTCCTGCAACTTTA
TCCGCCTCCATCCAGTCTATTAATTGTTGCCGGGAAGCTAGAGTAAGTAGTTCGCCAGTTAATAGTTTGCGCAACGT
TGTTGCCATTGCTACAGGCATCGTGGTGTCACGCTCGTCGTTTGGTATGGCTTCATTCAGCTCCGGTTCCCAACGAT
CAAGGCGAGTTACATGATCCCCCATGTTGTGCAAAAAAGCGGTTAGCTCCTTCGGTCCTCCGATCGTTGTCAGAAGT

AAGTTGGCCGCAGTGTTATCACTCATGGTTATGGCAGCACTGCATAATTCTCTTACTGTCATGCCATCCGTAAGATG
CTTTTCTGTGACTGGTGAGTACTCAACCAAGTCATTCTGAGAATAGTGTATGCGGCGACCGAGTTGCTCTTGCCCGG
CGTCAATACGGGATAATACCGCGCCACATAGCAGAACTTTAAAAGTGCTCATCATTGGAAAACGTTCTTCGGGGCGA
AAACTCTCAAGGATCTTACCGCTGTTGAGATCCAGTTCGATGTAACCCACTCGTGCACCCAACTGATCTTCAGCATC
TTTTACTTTCACCAGCGTTTCTGGGTGAGCAAAAACAGGAAGGCAAAATGCCGCAAAAAAGGGAATAAGGGCGACAC
GGAAATGTTGAATACTCATACTCTTCCTTTTTCAATATTATTGAAGCATTTATCAGGGTTATTGTCTCATGAGCGGA
TACATATTTGAATGTATTTAGAAAAATAAACAAATAGGGGTTCCGCGCACATTTCCCCGAAAAGTGCCACCTGACGT
C

>ARX R44Q in Arx_OMu17806D_pcDNA3.1+/C-(K)-DYK

GACGGATCGGGAGATCTCCCGATCCCCTATGGTGCACTCTCAGTACAATCTGCTCTGATGCCGCATAGTTAAGCCAG
TATCTGCTCCCTGCTTGTGTGTTGGAGGTCGCTGAGTAGTGCGCGAGCAAAATTTAAGCTACAACAAGGCAAGGCTT
GACCGACAATTGCATGAAGAATCTGCTTAGGGTTAGGCGTTTTGCGCTGCTTCGCGATGTACGGGCCAGATATACGC
GTTGACATTGATTATTGACTAGTTATTAATAGTAATCAATTACGGGGTCATTAGTTCATAGCCCATATATGGAGTTC
CGCGTTACATAACTTACGGTAAATGGCCCGCCTGGCTGACCGCCCAACGACCCCCGCCCATTGACGTCAATAATGAC
GTATGTTCCCATAGTAACGCCAATAGGGACTTTCCATTGACGTCAATGGGTGGAGTATTTACGGTAAACTGCCCACT
TGGCAGTACATCAAGTGTATCATATGCCAAGTACGCCCCCTATTGACGTCAATGACGGTAAATGGCCCGCCTGGCAT
TATGCCCAGTACATGACCTTATGGGACTTTCCTACTTGGCAGTACATCTACGTATTAGTCATCGCTATTACCATGGT
GATGCGGTTTTGGCAGTACATCAATGGGCGTGGATAGCGGTTTGACTCACGGGGATTTCCAAGTCTCCACCCCATTG
ACGTCAATGGGAGTTTGTTTTGGCACCAAAATCAACGGGACTTTCCAAAATGTCGTAACAACTCCGCCCCATTGACG
CAAATGGGCGGTAGGCGTGTACGGTGGGAGGTCTATATAAGCAGAGCTCTCTGGCTAACTAGAGAACCCACTGCTTA
CTGGCTTATCGAAATTAATACGACTCACTATAGGGAGACCCAAGCTGGCTAGCGTTTAAACTTAAGCTTGGTACCGA
GCTCGGATCCGCCACCATGAGCAATCAGTACCAGGAAGAGGGCTGCTCTGAGAGGCCGGAGTGCAAGAGTAAATCTC
CAACTTTGCTCTCCTCCTACTGCATCGACAGCATCCTGGGCCGGAGAAGCCCATGCAAAATGAGGCTTCTGGGAGCC
GCGCAGAGCTTGCCTGCCCCGCTGGCCAGCCGCGCTGACCAGGAAAAAGCCATGCAAGGCTCCCCCAAGAGCAGCAG
CGCCCCGTTCGAGGCTGAGCTGCACCTGCCACCCAAGCTGCGGCGCCTCTACGGCCCGGGCGGGGGCCGCCTCCTCC
AGGGCGCGGCGGCGGCGGCTGCAGCGGCGGCGGCGGCCGCAGCGGCGGCCACGGCCACTGGCACCGCAGGTCCCCGT
GGAGAGGTTCCTCCACCGCCACCACCAGCTGCACGGCCAGGGGAGCGTCAGGACAGCGCAGGGGCCGTGGCTGCCGC
CGCTGCCGCCGCTGCCTGGGACACGCTCAAGATCAGCCAGGCGCCGCAGGTGAGCATCAGTCGTAGCAAGTCGTACC
GCGAGAACGGGGCGCCCTTCGTGCCCCCGCCGCCAGCCCTGGACGAGCTGAGTGGCCCGGGGGGCGTCGCGCACCCC
GAGGAGCGCCTCAGCGCGGCCAGCGGACCCGGCAGTGCCCCCGCGGCGGGTGGTGGCACGGGCGCCGAGGACGACGA
GGAGGAGCTGTTGGAGGATGAGGAAGATGAAGAGGAGGAAGAGGAGCTGCTGGAGGACGACGACGAGGAGCTGCTGG
AGGACGATGCCCGCGCGCTGCTCAAGGAGCCCCGGCGCTGCTCGGTGGCCACCACCGGCACAGTGGCCGCCGCCGCC
GCCGCTGCTGCCGCCGCCGTGGCCACCGAGGGCGGGGAGCTGTCGCCCAAGGAAGAGCTGTTGCTGCACCCGGAGGA
CGCCGAGGGCAAGGATGGTGAGGACAGCGTGTGCCTGTCGGCCGGCAGCGACTCGGAGGAGGGGCTGCTGAAGCGCA
AACAGAGGCGCTACCGCACCACGTTCACCAGTTACCAGCTGGAGGAACTGGAGCGGGCTTTCCAGAAGACGCACTAC
CCTGACGTCTTCACCAGGGAGGAGCTGGCCATGAGGCTGGACCTGACAGAGGCCCAAGTCCAGGTGTGGTTCCAGAA
CCGTCGGGCCAAGTGGCGCAAGCGGGAGAAGGCTGGCGCACAGACCCACCCCCCGGGGCTGCCCTTTCCAGGGCCGC
TCTCGGCTACCCACCCTCTCAGCCCCTACCTGGACGCCAGCCCCTTTCCCCCGCATCACCCTGCGCTTGATTCGGCC
TGGACCGCCGCGGCGGCGGCGGCTGCAGCTGCCTTCCCGAGCCTACCTCCGCCCCCAGGCTCTGCTAGCCTGCCACC
CAGCGGGGCGCCGCTGGGTCTGAGCACTTTTCTAGGAGCAGCGGTGTTCCGCCACCCAGCCTTCATCAGCCCAGCAT
TTGGCAGGCTCTTTTCCACTATGGCCCCCCTGACCAGCGCGTCGACTGCGGCTGCGCTCCTGAGACAGCCCACACCC
GCGGTGGAGGGCGCAGTGGCCTCGGGCGCGCTGGCCGACCCGGCCACCGCCGCGGCAGACAGACGCGCGTCAAGCAT
AGCCGCGCTGAGGCTCAAGGCTAAGGAGCACGCCGCGCAGCTCACGCAGCTCAACATCCTGCCGGGCACCAGCACGG
GGAAGGAGGTGTGCGATTACAAGGATGACGACGATAAGTGATAAACCCGCTGATCAGCCTCGACTGTGCCTTCTAGT
TGCCAGCCATCTGTTGTTTGCCCCTCCCCCGTGCCTTCCTTGACCCTGGAAGGTGCCACTCCCACTGTCCTTTCCTA
ATAAAATGAGGAAATTGCATCGCATTGTCTGAGTAGGTGTCATTCTATTCTGGGGGGTGGGGTGGGGCAGGACAGCA
AGGGGGAGGATTGGGAAGACAATAGCAGGCATGCTGGGGATGCGGTGGGCTCTATGGCTTCTGAGGCGGAAAGAACC
AGCTGGGGCTCTAGGGGGTATCCCCACGCGCCCTGTAGCGGCGCATTAAGCGCGGCGGGTGTGGTGGTTACGCGCAG
CGTGACCGCTACACTTGCCAGCGCCCTAGCGCCCGCTCCTTTCGCTTTCTTCCCTTCCTTTCTCGCCACGTTCGCCG
GCTTTCCCCGTCAAGCTCTAAATCGGGGGCTCCCTTTAGGGTTCCGATTTAGTGCTTTACGGCACCTCGACCCCAAA
AAACTTGATTAGGGTGATGGTTCACGTAGTGGGCCATCGCCCTGATAGACGGTTTTTCGCCCTTTGACGTTGGAGTC

CACGTTCTTTAATAGTGGACTCTTGTTCCAAACTGGAACAACACTCAACCCTATCTCGGTCTATTCTTTTGATTTAT
AAGGGATTTTGCCGATTTCGGCCTATTGGTTAAAAAATGAGCTGATTTAACAAAAATTTAACGCGAATTAATTCTGT
GGAATGTGTGTCAGTTAGGGTGTGGAAAGTCCCCAGGCTCCCCAGCAGGCAGAAGTATGCAAAGCATGCATCTCAAT
TAGTCAGCAACCAGGTGTGGAAAGTCCCCAGGCTCCCCAGCAGGCAGAAGTATGCAAAGCATGCATCTCAATTAGTC
AGCAACCATAGTCCCGCCCCTAACTCCGCCCATCCCGCCCCTAACTCCGCCCAGTTCCGCCCATTCTCCGCCCCATG
GCTGACTAATTTTTTTTATTTATGCAGAGGCCGAGGCCGCCTCTGCCTCTGAGCTATTCCAGAAGTAGTGAGGAGGC
TTTTTTGGAGGCCTAGGCTTTTGCAAAAAGCTCCCGGGAGCTTGTATATCCATTTTCGGATCTGATCAAGAGACAGG
ATGAGGATCGTTTCGCATGATTGAACAAGATGGATTGCACGCAGGTTCTCCGGCCGCTTGGGTGGAGAGGCTATTCG
GCTATGACTGGGCACAACAGACAATCGGCTGCTCTGATGCCGCCGTGTTCCGGCTGTCAGCGCAGGGGCGCCCGGTT
CTTTTTGTCAAGACCGACCTGTCCGGTGCCCTGAATGAACTGCAGGACGAGGCAGCGCGGCTATCGTGGCTGGCCAC
GACGGGCGTTCCTTGCGCAGCTGTGCTCGACGTTGTCACTGAAGCGGGAAGGGACTGGCTGCTATTGGGCGAAGTGC
CGGGGCAGGATCTCCTGTCATCTCACCTTGCTCCTGCCGAGAAAGTATCCATCATGGCTGATGCAATGCGGCGGCTG
CATACGCTTGATCCGGCTACCTGCCCATTCGACCACCAAGCGAAACATCGCATCGAGCGAGCACGTACTCGGATGGA
AGCCGGTCTTGTCGATCAGGATGATCTGGACGAAGAGCATCAGGGGCTCGCGCCAGCCGAACTGTTCGCCAGGCTCA
AGGCGCGCATGCCCGACGGCGAGGATCTCGTCGTGACCCATGGCGATGCCTGCTTGCCGAATATCATGGTGGAAAAT
GGCCGCTTTTCTGGATTCATCGACTGTGGCCGGCTGGGTGTGGCGGACCGCTATCAGGACATAGCGTTGGCTACCCG
TGATATTGCTGAAGAGCTTGGCGGCGAATGGGCTGACCGCTTCCTCGTGCTTTACGGTATCGCCGCTCCCGATTCGC
AGCGCATCGCCTTCTATCGCCTTCTTGACGAGTTCTTCTGAGCGGGACTCTGGGGTTCGAAATGACCGACCAAGCGA
CGCCCAACCTGCCATCACGAGATTTCGATTCCACCGCCGCCTTCTATGAAAGGTTGGGCTTCGGAATCGTTTTCCGG
GACGCCGGCTGGATGATCCTCCAGCGCGGGGATCTCATGCTGGAGTTCTTCGCCCACCCCAACTTGTTTATTGCAGC
TTATAATGGTTACAAATAAAGCAATAGCATCACAAATTTCACAAATAAAGCATTTTTTTCACTGCATTCTAGTTGTG
GTTTGTCCAAACTCATCAATGTATCTTATCATGTCTGTATACCGTCGACCTCTAGCTAGAGCTTGGCGTAATCATGG
TCATAGCTGTTTCCTGTGTGAAATTGTTATCCGCTCACAATTCCACACAACATACGAGCCGGAAGCATAAAGTGTAA
AGCCTGGGGTGCCTAATGAGTGAGCTAACTCACATTAATTGCGTTGCGCTCACTGCCCGCTTTCCAGTCGGGAAACC
TGTCGTGCCAGCTGCATTAATGAATCGGCCAACGCGCGGGGAGAGGCGGTTTGCGTATTGGGCGCTCTTCCGCTTCC
TCGCTCACTGACTCGCTGCGCTCGGTCGTTCGGCTGCGGCGAGCGGTATCAGCTCACTCAAAGGCGGTAATACGGTT
ATCCACAGAATCAGGGGATAACGCAGGAAAGAACATGTGAGCAAAAGGCCAGCAAAAGGCCAGGAACCGTAAAAAGG
CCGCGTTGCTGGCGTTTTTCCATAGGCTCCGCCCCCCTGACGAGCATCACAAAAATCGACGCTCAAGTCAGAGGTGG
CGAAACCCGACAGGACTATAAAGATACCAGGCGTTTCCCCCTGGAAGCTCCCTCGTGCGCTCTCCTGTTCCGACCCT
GCCGCTTACCGGATACCTGTCCGCCTTTCTCCCTTCGGGAAGCGTGGCGCTTTCTCATAGCTCACGCTGTAGGTATC
TCAGTTCGGTGTAGGTCGTTCGCTCCAAGCTGGGCTGTGTGCACGAACCCCCCGTTCAGCCCGACCGCTGCGCCTTA
TCCGGTAACTATCGTCTTGAGTCCAACCCGGTAAGACACGACTTATCGCCACTGGCAGCAGCCACTGGTAACAGGAT
TAGCAGAGCGAGGTATGTAGGCGGTGCTACAGAGTTCTTGAAGTGGTGGCCTAACTACGGCTACACTAGAAGAACAG
TATTTGGTATCTGCGCTCTGCTGAAGCCAGTTACCTTCGGAAAAAGAGTTGGTAGCTCTTGATCCGGCAAACAAACC
ACCGCTGGTAGCGGTGGTTTTTTTGTTTGCAAGCAGCAGATTACGCGCAGAAAAAAAGGATCTCAAGAAGATCCTTT
GATCTTTTCTACGGGGTCTGACGCTCAGTGGAACGAAAACTCACGTTAAGGGATTTTGGTCATGAGATTATCAAAAA
GGATCTTCACCTAGATCCTTTTAAATTAAAAATGAAGTTTTAAATCAATCTAAAGTATATATGAGTAAACTTGGTCT
GACAGTTACCAATGCTTAATCAGTGAGGCACCTATCTCAGCGATCTGTCTATTTCGTTCATCCATAGTTGCCTGACT
CCCCGTCGTGTAGATAACTACGATACGGGAGGGCTTACCATCTGGCCCCAGTGCTGCAATGATACCGCGAGACCCAC
GCTCACCGGCTCCAGATTTATCAGCAATAAACCAGCCAGCCGGAAGGGCCGAGCGCAGAAGTGGTCCTGCAACTTTA
TCCGCCTCCATCCAGTCTATTAATTGTTGCCGGGAAGCTAGAGTAAGTAGTTCGCCAGTTAATAGTTTGCGCAACGT
TGTTGCCATTGCTACAGGCATCGTGGTGTCACGCTCGTCGTTTGGTATGGCTTCATTCAGCTCCGGTTCCCAACGAT
CAAGGCGAGTTACATGATCCCCCATGTTGTGCAAAAAAGCGGTTAGCTCCTTCGGTCCTCCGATCGTTGTCAGAAGT

AAGTTGGCCGCAGTGTTATCACTCATGGTTATGGCAGCACTGCATAATTCTCTTACTGTCATGCCATCCGTAAGATG
CTTTTCTGTGACTGGTGAGTACTCAACCAAGTCATTCTGAGAATAGTGTATGCGGCGACCGAGTTGCTCTTGCCCGG
CGTCAATACGGGATAATACCGCGCCACATAGCAGAACTTTAAAAGTGCTCATCATTGGAAAACGTTCTTCGGGGCGA
AAACTCTCAAGGATCTTACCGCTGTTGAGATCCAGTTCGATGTAACCCACTCGTGCACCCAACTGATCTTCAGCATC
TTTTACTTTCACCAGCGTTTCTGGGTGAGCAAAAACAGGAAGGCAAAATGCCGCAAAAAAGGGAATAAGGGCGACAC
GGAAATGTTGAATACTCATACTCTTCCTTTTTCAATATTATTGAAGCATTTATCAGGGTTATTGTCTCATGAGCGGA
TACATATTTGAATGTATTTAGAAAAATAAACAAATAGGGGTTCCGCGCACATTTCCCCGAAAAGTGCCACCTGACGT
C

>PHOX2B R44Q in pUC57

TCGCGCGTTTCGGTGATGACGGTGAAAACCTCTGACACATGCAGCTCCCGGAGACGGTCACAGCTTGTCTGTAAGCG
GATGCCGGGAGCAGACAAGCCCGTCAGGGCGCGTCAGCGGGTGTTGGCGGGTGTCGGGGCTGGCTTAACTATGCGGC
ATCAGAGCAGATTGTACTGAGAGTGCACCATATGCGGTGTGAAATACCGCACAGATGCGTAAGGAGAAAATACCGCA
TCAGGCGCCATTCGCCATTCAGGCTGCGCAACTGTTGGGAAGGGCGATCGGTGCGGGCCTCTTCGCTATTACGCCAG
CTGGCGAAAGGGGGATGTGCTGCAAGGCGATTAAGTTGGGTAACGCCAGGGTTTTCCCAGTCACGACGTTGTAAAAC
GACGGCCAGTGAATTCGAGCTCGGTACCTCGCGAATGCATCTAGATCATATGAACGAGAAGCGCAAGCAGCGGCGCA
TCCGCACCACTTTCACCAGTGCCCAGCTCAAAGAGCTGGAAAGGGTCTTCGCGGAGACTCACTACCCCGACATCTAC
ACTCGGGAGGAGCTGGCCCTGAAGATCGACCTCACAGAGGCGCAAGTCCAGGTGTGGTTCCAGAACCGCCGCGCCAA
GTTTCGCAAGCAGGAGCGCTAACTCGAGATCGGATCCCGGGCCCGTCGACTGCAGAGGCCTGCATGCAAGCTTGGCG
TAATCATGGTCATAGCTGTTTCCTGTGTGAAATTGTTATCCGCTCACAATTCCACACAACATACGAGCCGGAAGCAT
AAAGTGTAAAGCCTGGGGTGCCTAATGAGTGAGCTAACTCACATTAATTGCGTTGCGCTCACTGCCCGCTTTCCAGT
CGGGAAACCTGTCGTGCCAGCTGCATTAATGAATCGGCCAACGCGCGGGGAGAGGCGGTTTGCGTATTGGGCGCTCT
TCCGCTTCCTCGCTCACTGACTCGCTGCGCTCGGTCGTTCGGCTGCGGCGAGCGGTATCAGCTCACTCAAAGGCGGT
AATACGGTTATCCACAGAATCAGGGGATAACGCAGGAAAGAACATGTGAGCAAAAGGCCAGCAAAAGGCCAGGAACC
GTAAAAAGGCCGCGTTGCTGGCGTTTTTCCATAGGCTCCGCCCCCCTGACGAGCATCACAAAAATCGACGCTCAAGT
CAGAGGTGGCGAAACCCGACAGGACTATAAAGATACCAGGCGTTTCCCCCTGGAAGCTCCCTCGTGCGCTCTCCTGT
TCCGACCCTGCCGCTTACCGGATACCTGTCCGCCTTTCTCCCTTCGGGAAGCGTGGCGCTTTCTCATAGCTCACGCT
GTAGGTATCTCAGTTCGGTGTAGGTCGTTCGCTCCAAGCTGGGCTGTGTGCACGAACCCCCCGTTCAGCCCGACCGC
TGCGCCTTATCCGGTAACTATCGTCTTGAGTCCAACCCGGTAAGACACGACTTATCGCCACTGGCAGCAGCCACTGG
TAACAGGATTAGCAGAGCGAGGTATGTAGGCGGTGCTACAGAGTTCTTGAAGTGGTGGCCTAACTACGGCTACACTA
GAAGAACAGTATTTGGTATCTGCGCTCTGCTGAAGCCAGTTACCTTCGGAAAAAGAGTTGGTAGCTCTTGATCCGGC
AAACAAACCACCGCTGGTAGCGGTGGTTTTTTTGTTTGCAAGCAGCAGATTACGCGCAGAAAAAAAGGATCTCAAGA
AGATCCTTTGATCTTTTCTACGGGGTCTGACGCTCAGTGGAACGAAAACTCACGTTAAGGGATTTTGGTCATGAGAT
TATCAAAAAGGATCTTCACCTAGATCCTTTTAAATTAAAAATGAAGTTTTAAATCAATCTAAAGTATATATGAGTAA
ACTTGGTCTGACAGTTACCAATGCTTAATCAGTGAGGCACCTATCTCAGCGATCTGTCTATTTCGTTCATCCATAGT
TGCCTGACTCCCCGTCGTGTAGATAACTACGATACGGGAGGGCTTACCATCTGGCCCCAGTGCTGCAATGATACCGC
GAGACCCACGCTCACCGGCTCCAGATTTATCAGCAATAAACCAGCCAGCCGGAAGGGCCGAGCGCAGAAGTGGTCCT
GCAACTTTATCCGCCTCCATCCAGTCTATTAATTGTTGCCGGGAAGCTAGAGTAAGTAGTTCGCCAGTTAATAGTTT
GCGCAACGTTGTTGCCATTGCTACAGGCATCGTGGTGTCACGCTCGTCGTTTGGTATGGCTTCATTCAGCTCCGGTT
CCCAACGATCAAGGCGAGTTACATGATCCCCCATGTTGTGCAAAAAAGCGGTTAGCTCCTTCGGTCCTCCGATCGTT
GTCAGAAGTAAGTTGGCCGCAGTGTTATCACTCATGGTTATGGCAGCACTGCATAATTCTCTTACTGTCATGCCATC
CGTAAGATGCTTTTCTGTGACTGGTGAGTACTCAACCAAGTCATTCTGAGAATAGTGTATGCGGCGACCGAGTTGCT
CTTGCCCGGCGTCAATACGGGATAATACCGCGCCACATAGCAGAACTTTAAAAGTGCTCATCATTGGAAAACGTTCT
TCGGGGCGAAAACTCTCAAGGATCTTACCGCTGTTGAGATCCAGTTCGATGTAACCCACTCGTGCACCCAACTGATC
TTCAGCATCTTTTACTTTCACCAGCGTTTCTGGGTGAGCAAAAACAGGAAGGCAAAATGCCGCAAAAAAGGGAATAA
GGGCGACACGGAAATGTTGAATACTCATACTCTTCCTTTTTCAATATTATTGAAGCATTTATCAGGGTTATTGTCTC
ATGAGCGGATACATATTTGAATGTATTTAGAAAAATAAACAAATAGGGGTTCCGCGCACATTTCCCCGAAAAGTGCC
ACCTGACGTCTAAGAAACCATTATTATCATGACATTAACCTATAAAAATAGGCGTATCACGAGGCCCTTTCGTC

>PHOX2B R44G in pUC57

TCGCGCGTTTCGGTGATGACGGTGAAAACCTCTGACACATGCAGCTCCCGGAGACGGTCACAGCTTGTCTGTAAGCG
GATGCCGGGAGCAGACAAGCCCGTCAGGGCGCGTCAGCGGGTGTTGGCGGGTGTCGGGGCTGGCTTAACTATGCGGC
ATCAGAGCAGATTGTACTGAGAGTGCACCATATGCGGTGTGAAATACCGCACAGATGCGTAAGGAGAAAATACCGCA
TCAGGCGCCATTCGCCATTCAGGCTGCGCAACTGTTGGGAAGGGCGATCGGTGCGGGCCTCTTCGCTATTACGCCAG
CTGGCGAAAGGGGGATGTGCTGCAAGGCGATTAAGTTGGGTAACGCCAGGGTTTTCCCAGTCACGACGTTGTAAAAC
GACGGCCAGTGAATTCGAGCTCGGTACCTCGCGAATGCATCTAGATCATATGAACGAGAAGCGCAAGCAGCGGCGCA
TCCGCACCACTTTCACCAGTGCCCAGCTCAAAGAGCTGGAAAGGGTCTTCGCGGAGACTCACTACCCCGACATCTAC
ACTCGGGAGGAGCTGGCCCTGAAGATCGACCTCACAGAGGCGGGAGTCCAGGTGTGGTTCCAGAACCGCCGCGCCAA
GTTTCGCAAGCAGGAGCGCTAACTCGAGATCGGATCCCGGGCCCGTCGACTGCAGAGGCCTGCATGCAAGCTTGGCG
TAATCATGGTCATAGCTGTTTCCTGTGTGAAATTGTTATCCGCTCACAATTCCACACAACATACGAGCCGGAAGCAT
AAAGTGTAAAGCCTGGGGTGCCTAATGAGTGAGCTAACTCACATTAATTGCGTTGCGCTCACTGCCCGCTTTCCAGT
CGGGAAACCTGTCGTGCCAGCTGCATTAATGAATCGGCCAACGCGCGGGGAGAGGCGGTTTGCGTATTGGGCGCTCT
TCCGCTTCCTCGCTCACTGACTCGCTGCGCTCGGTCGTTCGGCTGCGGCGAGCGGTATCAGCTCACTCAAAGGCGGT
AATACGGTTATCCACAGAATCAGGGGATAACGCAGGAAAGAACATGTGAGCAAAAGGCCAGCAAAAGGCCAGGAACC
GTAAAAAGGCCGCGTTGCTGGCGTTTTTCCATAGGCTCCGCCCCCCTGACGAGCATCACAAAAATCGACGCTCAAGT
CAGAGGTGGCGAAACCCGACAGGACTATAAAGATACCAGGCGTTTCCCCCTGGAAGCTCCCTCGTGCGCTCTCCTGT
TCCGACCCTGCCGCTTACCGGATACCTGTCCGCCTTTCTCCCTTCGGGAAGCGTGGCGCTTTCTCATAGCTCACGCT
GTAGGTATCTCAGTTCGGTGTAGGTCGTTCGCTCCAAGCTGGGCTGTGTGCACGAACCCCCCGTTCAGCCCGACCGC
TGCGCCTTATCCGGTAACTATCGTCTTGAGTCCAACCCGGTAAGACACGACTTATCGCCACTGGCAGCAGCCACTGG
TAACAGGATTAGCAGAGCGAGGTATGTAGGCGGTGCTACAGAGTTCTTGAAGTGGTGGCCTAACTACGGCTACACTA
GAAGAACAGTATTTGGTATCTGCGCTCTGCTGAAGCCAGTTACCTTCGGAAAAAGAGTTGGTAGCTCTTGATCCGGC
AAACAAACCACCGCTGGTAGCGGTGGTTTTTTTGTTTGCAAGCAGCAGATTACGCGCAGAAAAAAAGGATCTCAAGA
AGATCCTTTGATCTTTTCTACGGGGTCTGACGCTCAGTGGAACGAAAACTCACGTTAAGGGATTTTGGTCATGAGAT
TATCAAAAAGGATCTTCACCTAGATCCTTTTAAATTAAAAATGAAGTTTTAAATCAATCTAAAGTATATATGAGTAA
ACTTGGTCTGACAGTTACCAATGCTTAATCAGTGAGGCACCTATCTCAGCGATCTGTCTATTTCGTTCATCCATAGT
TGCCTGACTCCCCGTCGTGTAGATAACTACGATACGGGAGGGCTTACCATCTGGCCCCAGTGCTGCAATGATACCGC
GAGACCCACGCTCACCGGCTCCAGATTTATCAGCAATAAACCAGCCAGCCGGAAGGGCCGAGCGCAGAAGTGGTCCT
GCAACTTTATCCGCCTCCATCCAGTCTATTAATTGTTGCCGGGAAGCTAGAGTAAGTAGTTCGCCAGTTAATAGTTT
GCGCAACGTTGTTGCCATTGCTACAGGCATCGTGGTGTCACGCTCGTCGTTTGGTATGGCTTCATTCAGCTCCGGTT
CCCAACGATCAAGGCGAGTTACATGATCCCCCATGTTGTGCAAAAAAGCGGTTAGCTCCTTCGGTCCTCCGATCGTT
GTCAGAAGTAAGTTGGCCGCAGTGTTATCACTCATGGTTATGGCAGCACTGCATAATTCTCTTACTGTCATGCCATC
CGTAAGATGCTTTTCTGTGACTGGTGAGTACTCAACCAAGTCATTCTGAGAATAGTGTATGCGGCGACCGAGTTGCT
CTTGCCCGGCGTCAATACGGGATAATACCGCGCCACATAGCAGAACTTTAAAAGTGCTCATCATTGGAAAACGTTCT
TCGGGGCGAAAACTCTCAAGGATCTTACCGCTGTTGAGATCCAGTTCGATGTAACCCACTCGTGCACCCAACTGATC
TTCAGCATCTTTTACTTTCACCAGCGTTTCTGGGTGAGCAAAAACAGGAAGGCAAAATGCCGCAAAAAAGGGAATAA
GGGCGACACGGAAATGTTGAATACTCATACTCTTCCTTTTTCAATATTATTGAAGCATTTATCAGGGTTATTGTCTC
ATGAGCGGATACATATTTGAATGTATTTAGAAAAATAAACAAATAGGGGTTCCGCGCACATTTCCCCGAAAAGTGCC
ACCTGACGTCTAAGAAACCATTATTATCATGACATTAACCTATAAAAATAGGCGTATCACGAGGCCCTTTCGTC

>PHOX2B Q46K in PHOX2B_OHu26329C_pcDNA3.1(+)-N-HA

GACGGATCGGGAGATCTCCCGATCCCCTATGGTGCACTCTCAGTACAATCTGCTCTGATGCCGCATAGTTAAGCCAG
TATCTGCTCCCTGCTTGTGTGTTGGAGGTCGCTGAGTAGTGCGCGAGCAAAATTTAAGCTACAACAAGGCAAGGCTT
GACCGACAATTGCATGAAGAATCTGCTTAGGGTTAGGCGTTTTGCGCTGCTTCGCGATGTACGGGCCAGATATACGC
GTTGACATTGATTATTGACTAGTTATTAATAGTAATCAATTACGGGGTCATTAGTTCATAGCCCATATATGGAGTTC
CGCGTTACATAACTTACGGTAAATGGCCCGCCTGGCTGACCGCCCAACGACCCCCGCCCATTGACGTCAATAATGAC
GTATGTTCCCATAGTAACGCCAATAGGGACTTTCCATTGACGTCAATGGGTGGAGTATTTACGGTAAACTGCCCACT
TGGCAGTACATCAAGTGTATCATATGCCAAGTACGCCCCCTATTGACGTCAATGACGGTAAATGGCCCGCCTGGCAT
TATGCCCAGTACATGACCTTATGGGACTTTCCTACTTGGCAGTACATCTACGTATTAGTCATCGCTATTACCATGGT
GATGCGGTTTTGGCAGTACATCAATGGGCGTGGATAGCGGTTTGACTCACGGGGATTTCCAAGTCTCCACCCCATTG
ACGTCAATGGGAGTTTGTTTTGGCACCAAAATCAACGGGACTTTCCAAAATGTCGTAACAACTCCGCCCCATTGACG
CAAATGGGCGGTAGGCGTGTACGGTGGGAGGTCTATATAAGCAGAGCTCTCTGGCTAACTAGAGAACCCACTGCTTA
CTGGCTTATCGAAATTAATACGACTCACTATAGGGAGACCCAAGCTGGCTAGCGTTTAAACTTAAGCTTGCCACCAT
GTACCCATACGATGTTCCAGATTACGCTGGCTCTGGATATAAAATGGAATATTCTTACCTCAATTCCTCTGCCTACG
AGTCCTGTATGGCTGGGATGGACACCTCGAGCCTGGCTTCAGCCTATGCTGACTTCAGTTCCTGCAGCCAGGCCAGT
GGCTTCCAGTATAACCCGATAAGGACCACTTTTGGGGCCACGTCCGGCTGCCCTTCCCTCACGCCGGGATCCTGCAG
CCTGGGCACCCTCAGGGACCACCAGAGCAGTCCGTACGCCGCAGTTCCTTACAAACTCTTCACGGACCACGGCGGCC
TCAACGAGAAGCGCAAGCAGCGGCGCATCCGCACCACTTTCACCAGTGCCCAGCTCAAAGAGCTGGAAAGGGTCTTC
GCGGAGACTCACTACCCCGACATCTACACTCGGGAGGAGCTGGCCCTGAAGATCGACCTCACAGAGGCGCGAGTCAA
GGTGTGGTTCCAGAACCGCCGCGCCAAGTTTCGCAAGCAGGAGCGCGCAGCGGCAGCCGCAGCGGCCGCGGCCAAGA
ACGGCTCCTCGGGCAAAAAGTCTGACTCTTCCAGGGACGACGAGAGCAAAGAGGCCAAGAGCACTGACCCGGACAGC
ACTGGGGGCCCAGGTCCCAATCCCAACCCCACCCCCAGCTGCGGGGCGAATGGAGGCGGCGGCGGCGGGCCCAGCCC
GGCTGGAGCTCCGGGGGCGGCGGGGCCCGGGGGCCCGGGAGGCGAACCCGGCAAGGGCGGCGCAGCAGCAGCGGCGG
CGGCCGCGGCAGCGGCGGCGGCGGCAGCGGCAGCGGCGGCAGCTGGAGGCCTGGCTGCGGCTGGGGGCCCTGGACAA
GGCTGGGCTCCCGGCCCCGGCCCCATCACCTCCATCCCGGATTCGCTTGGGGGTCCCTTCGCCAGCGTCCTATCTTC
GCTCCAAAGACCCAACGGTGCCAAAGCCGCCTTAGTGAAGAGCAGTATGTTCTGAGGTACCGAGCTCGGATCCGAAT
TCTGCAGATATCCAGCACAGTGGCGGCCGCTCGAGTCTAGAGGGCCCGTTTAAACCCGCTGATCAGCCTCGACTGTG
CCTTCTAGTTGCCAGCCATCTGTTGTTTGCCCCTCCCCCGTGCCTTCCTTGACCCTGGAAGGTGCCACTCCCACTGT
CCTTTCCTAATAAAATGAGGAAATTGCATCGCATTGTCTGAGTAGGTGTCATTCTATTCTGGGGGGTGGGGTGGGGC
AGGACAGCAAGGGGGAGGATTGGGAAGACAATAGCAGGCATGCTGGGGATGCGGTGGGCTCTATGGCTTCTGAGGCG
GAAAGAACCAGCTGGGGCTCTAGGGGGTATCCCCACGCGCCCTGTAGCGGCGCATTAAGCGCGGCGGGTGTGGTGGT
TACGCGCAGCGTGACCGCTACACTTGCCAGCGCCCTAGCGCCCGCTCCTTTCGCTTTCTTCCCTTCCTTTCTCGCCA
CGTTCGCCGGCTTTCCCCGTCAAGCTCTAAATCGGGGGCTCCCTTTAGGGTTCCGATTTAGTGCTTTACGGCACCTC
GACCCCAAAAAACTTGATTAGGGTGATGGTTCACGTAGTGGGCCATCGCCCTGATAGACGGTTTTTCGCCCTTTGAC
GTTGGAGTCCACGTTCTTTAATAGTGGACTCTTGTTCCAAACTGGAACAACACTCAACCCTATCTCGGTCTATTCTT
TTGATTTATAAGGGATTTTGCCGATTTCGGCCTATTGGTTAAAAAATGAGCTGATTTAACAAAAATTTAACGCGAAT
TAATTCTGTGGAATGTGTGTCAGTTAGGGTGTGGAAAGTCCCCAGGCTCCCCAGCAGGCAGAAGTATGCAAAGCATG
CATCTCAATTAGTCAGCAACCAGGTGTGGAAAGTCCCCAGGCTCCCCAGCAGGCAGAAGTATGCAAAGCATGCATCT
CAATTAGTCAGCAACCATAGTCCCGCCCCTAACTCCGCCCATCCCGCCCCTAACTCCGCCCAGTTCCGCCCATTCTC
CGCCCCATGGCTGACTAATTTTTTTTATTTATGCAGAGGCCGAGGCCGCCTCTGCCTCTGAGCTATTCCAGAAGTAG
TGAGGAGGCTTTTTTGGAGGCCTAGGCTTTTGCAAAAAGCTCCCGGGAGCTTGTATATCCATTTTCGGATCTGATCA
AGAGACAGGATGAGGATCGTTTCGCATGATTGAACAAGATGGATTGCACGCAGGTTCTCCGGCCGCTTGGGTGGAGA
GGCTATTCGGCTATGACTGGGCACAACAGACAATCGGCTGCTCTGATGCCGCCGTGTTCCGGCTGTCAGCGCAGGGG

CGCCCGGTTCTTTTTGTCAAGACCGACCTGTCCGGTGCCCTGAATGAACTGCAGGACGAGGCAGCGCGGCTATCGTG
GCTGGCCACGACGGGCGTTCCTTGCGCAGCTGTGCTCGACGTTGTCACTGAAGCGGGAAGGGACTGGCTGCTATTGG
GCGAAGTGCCGGGGCAGGATCTCCTGTCATCTCACCTTGCTCCTGCCGAGAAAGTATCCATCATGGCTGATGCAATG
CGGCGGCTGCATACGCTTGATCCGGCTACCTGCCCATTCGACCACCAAGCGAAACATCGCATCGAGCGAGCACGTAC
TCGGATGGAAGCCGGTCTTGTCGATCAGGATGATCTGGACGAAGAGCATCAGGGGCTCGCGCCAGCCGAACTGTTCG
CCAGGCTCAAGGCGCGCATGCCCGACGGCGAGGATCTCGTCGTGACCCATGGCGATGCCTGCTTGCCGAATATCATG
GTGGAAAATGGCCGCTTTTCTGGATTCATCGACTGTGGCCGGCTGGGTGTGGCGGACCGCTATCAGGACATAGCGTT
GGCTACCCGTGATATTGCTGAAGAGCTTGGCGGCGAATGGGCTGACCGCTTCCTCGTGCTTTACGGTATCGCCGCTC
CCGATTCGCAGCGCATCGCCTTCTATCGCCTTCTTGACGAGTTCTTCTGAGCGGGACTCTGGGGTTCGAAATGACCG
ACCAAGCGACGCCCAACCTGCCATCACGAGATTTCGATTCCACCGCCGCCTTCTATGAAAGGTTGGGCTTCGGAATC
GTTTTCCGGGACGCCGGCTGGATGATCCTCCAGCGCGGGGATCTCATGCTGGAGTTCTTCGCCCACCCCAACTTGTT
TATTGCAGCTTATAATGGTTACAAATAAAGCAATAGCATCACAAATTTCACAAATAAAGCATTTTTTTCACTGCATT
CTAGTTGTGGTTTGTCCAAACTCATCAATGTATCTTATCATGTCTGTATACCGTCGACCTCTAGCTAGAGCTTGGCG
TAATCATGGTCATAGCTGTTTCCTGTGTGAAATTGTTATCCGCTCACAATTCCACACAACATACGAGCCGGAAGCAT
AAAGTGTAAAGCCTGGGGTGCCTAATGAGTGAGCTAACTCACATTAATTGCGTTGCGCTCACTGCCCGCTTTCCAGT
CGGGAAACCTGTCGTGCCAGCTGCATTAATGAATCGGCCAACGCGCGGGGAGAGGCGGTTTGCGTATTGGGCGCTCT
TCCGCTTCCTCGCTCACTGACTCGCTGCGCTCGGTCGTTCGGCTGCGGCGAGCGGTATCAGCTCACTCAAAGGCGGT
AATACGGTTATCCACAGAATCAGGGGATAACGCAGGAAAGAACATGTGAGCAAAAGGCCAGCAAAAGGCCAGGAACC
GTAAAAAGGCCGCGTTGCTGGCGTTTTTCCATAGGCTCCGCCCCCCTGACGAGCATCACAAAAATCGACGCTCAAGT
CAGAGGTGGCGAAACCCGACAGGACTATAAAGATACCAGGCGTTTCCCCCTGGAAGCTCCCTCGTGCGCTCTCCTGT
TCCGACCCTGCCGCTTACCGGATACCTGTCCGCCTTTCTCCCTTCGGGAAGCGTGGCGCTTTCTCATAGCTCACGCT
GTAGGTATCTCAGTTCGGTGTAGGTCGTTCGCTCCAAGCTGGGCTGTGTGCACGAACCCCCCGTTCAGCCCGACCGC
TGCGCCTTATCCGGTAACTATCGTCTTGAGTCCAACCCGGTAAGACACGACTTATCGCCACTGGCAGCAGCCACTGG
TAACAGGATTAGCAGAGCGAGGTATGTAGGCGGTGCTACAGAGTTCTTGAAGTGGTGGCCTAACTACGGCTACACTA
GAAGAACAGTATTTGGTATCTGCGCTCTGCTGAAGCCAGTTACCTTCGGAAAAAGAGTTGGTAGCTCTTGATCCGGC
AAACAAACCACCGCTGGTAGCGGTGGTTTTTTTGTTTGCAAGCAGCAGATTACGCGCAGAAAAAAAGGATCTCAAGA
AGATCCTTTGATCTTTTCTACGGGGTCTGACGCTCAGTGGAACGAAAACTCACGTTAAGGGATTTTGGTCATGAGAT
TATCAAAAAGGATCTTCACCTAGATCCTTTTAAATTAAAAATGAAGTTTTAAATCAATCTAAAGTATATATGAGTAA
ACTTGGTCTGACAGTTACCAATGCTTAATCAGTGAGGCACCTATCTCAGCGATCTGTCTATTTCGTTCATCCATAGT
TGCCTGACTCCCCGTCGTGTAGATAACTACGATACGGGAGGGCTTACCATCTGGCCCCAGTGCTGCAATGATACCGC
GAGACCCACGCTCACCGGCTCCAGATTTATCAGCAATAAACCAGCCAGCCGGAAGGGCCGAGCGCAGAAGTGGTCCT
GCAACTTTATCCGCCTCCATCCAGTCTATTAATTGTTGCCGGGAAGCTAGAGTAAGTAGTTCGCCAGTTAATAGTTT
GCGCAACGTTGTTGCCATTGCTACAGGCATCGTGGTGTCACGCTCGTCGTTTGGTATGGCTTCATTCAGCTCCGGTT
CCCAACGATCAAGGCGAGTTACATGATCCCCCATGTTGTGCAAAAAAGCGGTTAGCTCCTTCGGTCCTCCGATCGTT
GTCAGAAGTAAGTTGGCCGCAGTGTTATCACTCATGGTTATGGCAGCACTGCATAATTCTCTTACTGTCATGCCATC
CGTAAGATGCTTTTCTGTGACTGGTGAGTACTCAACCAAGTCATTCTGAGAATAGTGTATGCGGCGACCGAGTTGCT
CTTGCCCGGCGTCAATACGGGATAATACCGCGCCACATAGCAGAACTTTAAAAGTGCTCATCATTGGAAAACGTTCT
TCGGGGCGAAAACTCTCAAGGATCTTACCGCTGTTGAGATCCAGTTCGATGTAACCCACTCGTGCACCCAACTGATC
TTCAGCATCTTTTACTTTCACCAGCGTTTCTGGGTGAGCAAAAACAGGAAGGCAAAATGCCGCAAAAAAGGGAATAA
GGGCGACACGGAAATGTTGAATACTCATACTCTTCCTTTTTCAATATTATTGAAGCATTTATCAGGGTTATTGTCTC
ATGAGCGGATACATATTTGAATGTATTTAGAAAAATAAACAAATAGGGGTTCCGCGCACATTTCCCCGAAAAGTGCC
ACCTGACGTC

>PHOX2B R3L in pUC57

TCGCGCGTTTCGGTGATGACGGTGAAAACCTCTGACACATGCAGCTCCCGGAGACGGTCACAGCTTGTCTGTAAGCG
GATGCCGGGAGCAGACAAGCCCGTCAGGGCGCGTCAGCGGGTGTTGGCGGGTGTCGGGGCTGGCTTAACTATGCGGC
ATCAGAGCAGATTGTACTGAGAGTGCACCATATGCGGTGTGAAATACCGCACAGATGCGTAAGGAGAAAATACCGCA
TCAGGCGCCATTCGCCATTCAGGCTGCGCAACTGTTGGGAAGGGCGATCGGTGCGGGCCTCTTCGCTATTACGCCAG
CTGGCGAAAGGGGGATGTGCTGCAAGGCGATTAAGTTGGGTAACGCCAGGGTTTTCCCAGTCACGACGTTGTAAAAC
GACGGCCAGTGAATTCGAGCTCGGTACCTCGCGAATGCATCTAGATCATATGAACGAGAAGCGCAAGCAGCGGCTCA
TCCGCACCACTTTCACCAGTGCCCAGCTCAAAGAGCTGGAAAGGGTCTTCGCGGAGACTCACTACCCCGACATCTAC
ACTCGGGAGGAGCTGGCCCTGAAGATCGACCTCACAGAGGCGCGAGTCCAGGTGTGGTTCCAGAACCGCCGCGCCAA
GTTTCGCAAGCAGGAGCGCTAACTCGAGATCGGATCCCGGGCCCGTCGACTGCAGAGGCCTGCATGCAAGCTTGGCG
TAATCATGGTCATAGCTGTTTCCTGTGTGAAATTGTTATCCGCTCACAATTCCACACAACATACGAGCCGGAAGCAT
AAAGTGTAAAGCCTGGGGTGCCTAATGAGTGAGCTAACTCACATTAATTGCGTTGCGCTCACTGCCCGCTTTCCAGT
CGGGAAACCTGTCGTGCCAGCTGCATTAATGAATCGGCCAACGCGCGGGGAGAGGCGGTTTGCGTATTGGGCGCTCT
TCCGCTTCCTCGCTCACTGACTCGCTGCGCTCGGTCGTTCGGCTGCGGCGAGCGGTATCAGCTCACTCAAAGGCGGT
AATACGGTTATCCACAGAATCAGGGGATAACGCAGGAAAGAACATGTGAGCAAAAGGCCAGCAAAAGGCCAGGAACC
GTAAAAAGGCCGCGTTGCTGGCGTTTTTCCATAGGCTCCGCCCCCCTGACGAGCATCACAAAAATCGACGCTCAAGT
CAGAGGTGGCGAAACCCGACAGGACTATAAAGATACCAGGCGTTTCCCCCTGGAAGCTCCCTCGTGCGCTCTCCTGT
TCCGACCCTGCCGCTTACCGGATACCTGTCCGCCTTTCTCCCTTCGGGAAGCGTGGCGCTTTCTCATAGCTCACGCT
GTAGGTATCTCAGTTCGGTGTAGGTCGTTCGCTCCAAGCTGGGCTGTGTGCACGAACCCCCCGTTCAGCCCGACCGC
TGCGCCTTATCCGGTAACTATCGTCTTGAGTCCAACCCGGTAAGACACGACTTATCGCCACTGGCAGCAGCCACTGG
TAACAGGATTAGCAGAGCGAGGTATGTAGGCGGTGCTACAGAGTTCTTGAAGTGGTGGCCTAACTACGGCTACACTA
GAAGAACAGTATTTGGTATCTGCGCTCTGCTGAAGCCAGTTACCTTCGGAAAAAGAGTTGGTAGCTCTTGATCCGGC
AAACAAACCACCGCTGGTAGCGGTGGTTTTTTTGTTTGCAAGCAGCAGATTACGCGCAGAAAAAAAGGATCTCAAGA
AGATCCTTTGATCTTTTCTACGGGGTCTGACGCTCAGTGGAACGAAAACTCACGTTAAGGGATTTTGGTCATGAGAT
TATCAAAAAGGATCTTCACCTAGATCCTTTTAAATTAAAAATGAAGTTTTAAATCAATCTAAAGTATATATGAGTAA
ACTTGGTCTGACAGTTACCAATGCTTAATCAGTGAGGCACCTATCTCAGCGATCTGTCTATTTCGTTCATCCATAGT
TGCCTGACTCCCCGTCGTGTAGATAACTACGATACGGGAGGGCTTACCATCTGGCCCCAGTGCTGCAATGATACCGC
GAGACCCACGCTCACCGGCTCCAGATTTATCAGCAATAAACCAGCCAGCCGGAAGGGCCGAGCGCAGAAGTGGTCCT
GCAACTTTATCCGCCTCCATCCAGTCTATTAATTGTTGCCGGGAAGCTAGAGTAAGTAGTTCGCCAGTTAATAGTTT
GCGCAACGTTGTTGCCATTGCTACAGGCATCGTGGTGTCACGCTCGTCGTTTGGTATGGCTTCATTCAGCTCCGGTT
CCCAACGATCAAGGCGAGTTACATGATCCCCCATGTTGTGCAAAAAAGCGGTTAGCTCCTTCGGTCCTCCGATCGTT
GTCAGAAGTAAGTTGGCCGCAGTGTTATCACTCATGGTTATGGCAGCACTGCATAATTCTCTTACTGTCATGCCATC
CGTAAGATGCTTTTCTGTGACTGGTGAGTACTCAACCAAGTCATTCTGAGAATAGTGTATGCGGCGACCGAGTTGCT
CTTGCCCGGCGTCAATACGGGATAATACCGCGCCACATAGCAGAACTTTAAAAGTGCTCATCATTGGAAAACGTTCT
TCGGGGCGAAAACTCTCAAGGATCTTACCGCTGTTGAGATCCAGTTCGATGTAACCCACTCGTGCACCCAACTGATC
TTCAGCATCTTTTACTTTCACCAGCGTTTCTGGGTGAGCAAAAACAGGAAGGCAAAATGCCGCAAAAAAGGGAATAA
GGGCGACACGGAAATGTTGAATACTCATACTCTTCCTTTTTCAATATTATTGAAGCATTTATCAGGGTTATTGTCTC
ATGAGCGGATACATATTTGAATGTATTTAGAAAAATAAACAAATAGGGGTTCCGCGCACATTTCCCCGAAAAGTGCC
ACCTGACGTCTAAGAAACCATTATTATCATGACATTAACCTATAAAAATAGGCGTATCACGAGGCCCTTTCGTC

>CRX R3W in CRX_OHu18946C_pcDNA3.1(+)-N-HA

GACGGATCGGGAGATCTCCCGATCCCCTATGGTGCACTCTCAGTACAATCTGCTCTGATGCCGCATAGTTAAGCCAG
TATCTGCTCCCTGCTTGTGTGTTGGAGGTCGCTGAGTAGTGCGCGAGCAAAATTTAAGCTACAACAAGGCAAGGCTT
GACCGACAATTGCATGAAGAATCTGCTTAGGGTTAGGCGTTTTGCGCTGCTTCGCGATGTACGGGCCAGATATACGC
GTTGACATTGATTATTGACTAGTTATTAATAGTAATCAATTACGGGGTCATTAGTTCATAGCCCATATATGGAGTTC
CGCGTTACATAACTTACGGTAAATGGCCCGCCTGGCTGACCGCCCAACGACCCCCGCCCATTGACGTCAATAATGAC
GTATGTTCCCATAGTAACGCCAATAGGGACTTTCCATTGACGTCAATGGGTGGAGTATTTACGGTAAACTGCCCACT
TGGCAGTACATCAAGTGTATCATATGCCAAGTACGCCCCCTATTGACGTCAATGACGGTAAATGGCCCGCCTGGCAT
TATGCCCAGTACATGACCTTATGGGACTTTCCTACTTGGCAGTACATCTACGTATTAGTCATCGCTATTACCATGGT
GATGCGGTTTTGGCAGTACATCAATGGGCGTGGATAGCGGTTTGACTCACGGGGATTTCCAAGTCTCCACCCCATTG
ACGTCAATGGGAGTTTGTTTTGGCACCAAAATCAACGGGACTTTCCAAAATGTCGTAACAACTCCGCCCCATTGACG
CAAATGGGCGGTAGGCGTGTACGGTGGGAGGTCTATATAAGCAGAGCTCTCTGGCTAACTAGAGAACCCACTGCTTA
CTGGCTTATCGAAATTAATACGACTCACTATAGGGAGACCCAAGCTGGCTAGCGTTTAAACTTAAGCTTGCCACCAT
GTACCCATACGATGTTCCAGATTACGCTGGCTCTGGAATGGCGTATATGAACCCGGGGCCCCACTATTCTGTCAACG
CCTTGGCCCTAAGTGGCCCCAGTGTGGATCTGATGCACCAGGCTGTGCCCTACCCAAGCGCCCCCAGGAAGCAGCGG
TGGGAGCGCACCACCTTCACCCGGAGCCAACTGGAGGAGCTGGAGGCACTGTTTGCCAAGACCCAGTACCCAGACGT
CTATGCCCGTGAGGAGGTGGCTCTGAAGATCAATCTGCCTGAGTCCAGGGTTCAGGTTTGGTTCAAGAACCGGAGGG
CTAAATGCAGGCAGCAGCGACAGCAGCAGAAACAGCAGCAGCAGCCCCCAGGGGGCCAGGCCAAGGCCCGGCCTGCC
AAGAGGAAGGCGGGCACGTCCCCAAGACCCTCCACAGATGTGTGTCCAGACCCTCTGGGCATCTCAGATTCCTACAG
TCCCCCTCTGCCCGGCCCCTCAGGCTCCCCAACCACGGCAGTGGCCACTGTGTCCATCTGGAGCCCAGCCTCAGAGT
CCCCTTTGCCTGAGGCGCAGCGGGCTGGGCTGGTGGCCTCAGGGCCGTCTCTGACCTCCGCCCCCTATGCCATGACC
TACGCCCCGGCCTCCGCTTTCTGCTCTTCCCCCTCCGCCTATGGGTCTCCGAGCTCCTATTTCAGCGGCCTAGACCC
CTACCTTTCTCCCATGGTGCCCCAGCTAGGGGGCCCGGCTCTTAGCCCCCTCTCTGGCCCCTCCGTGGGACCTTCCC
TGGCCCAGTCCCCCACCTCCCTATCAGGCCAGAGCTATGGCGCCTACAGCCCCGTGGATAGCTTGGAATTCAAGGAC
CCCACGGGCACCTGGAAATTCACCTACAATCCCATGGACCCTCTGGACTACAAGGATCAGAGTGCCTGGAAGTTTCA
GATCTTGTGATAAACCCGCTGATCAGCCTCGACTGTGCCTTCTAGTTGCCAGCCATCTGTTGTTTGCCCCTCCCCCG
TGCCTTCCTTGACCCTGGAAGGTGCCACTCCCACTGTCCTTTCCTAATAAAATGAGGAAATTGCATCGCATTGTCTG
AGTAGGTGTCATTCTATTCTGGGGGGTGGGGTGGGGCAGGACAGCAAGGGGGAGGATTGGGAAGACAATAGCAGGCA
TGCTGGGGATGCGGTGGGCTCTATGGCTTCTGAGGCGGAAAGAACCAGCTGGGGCTCTAGGGGGTATCCCCACGCGC
CCTGTAGCGGCGCATTAAGCGCGGCGGGTGTGGTGGTTACGCGCAGCGTGACCGCTACACTTGCCAGCGCCCTAGCG
CCCGCTCCTTTCGCTTTCTTCCCTTCCTTTCTCGCCACGTTCGCCGGCTTTCCCCGTCAAGCTCTAAATCGGGGGCT
CCCTTTAGGGTTCCGATTTAGTGCTTTACGGCACCTCGACCCCAAAAAACTTGATTAGGGTGATGGTTCACGTAGTG
GGCCATCGCCCTGATAGACGGTTTTTCGCCCTTTGACGTTGGAGTCCACGTTCTTTAATAGTGGACTCTTGTTCCAA
ACTGGAACAACACTCAACCCTATCTCGGTCTATTCTTTTGATTTATAAGGGATTTTGCCGATTTCGGCCTATTGGTT
AAAAAATGAGCTGATTTAACAAAAATTTAACGCGAATTAATTCTGTGGAATGTGTGTCAGTTAGGGTGTGGAAAGTC
CCCAGGCTCCCCAGCAGGCAGAAGTATGCAAAGCATGCATCTCAATTAGTCAGCAACCAGGTGTGGAAAGTCCCCAG
GCTCCCCAGCAGGCAGAAGTATGCAAAGCATGCATCTCAATTAGTCAGCAACCATAGTCCCGCCCCTAACTCCGCCC
ATCCCGCCCCTAACTCCGCCCAGTTCCGCCCATTCTCCGCCCCATGGCTGACTAATTTTTTTTATTTATGCAGAGGC
CGAGGCCGCCTCTGCCTCTGAGCTATTCCAGAAGTAGTGAGGAGGCTTTTTTGGAGGCCTAGGCTTTTGCAAAAAGC
TCCCGGGAGCTTGTATATCCATTTTCGGATCTGATCAAGAGACAGGATGAGGATCGTTTCGCATGATTGAACAAGAT
GGATTGCACGCAGGTTCTCCGGCCGCTTGGGTGGAGAGGCTATTCGGCTATGACTGGGCACAACAGACAATCGGCTG
CTCTGATGCCGCCGTGTTCCGGCTGTCAGCGCAGGGGCGCCCGGTTCTTTTTGTCAAGACCGACCTGTCCGGTGCCC
TGAATGAACTGCAGGACGAGGCAGCGCGGCTATCGTGGCTGGCCACGACGGGCGTTCCTTGCGCAGCTGTGCTCGAC

GTTGTCACTGAAGCGGGAAGGGACTGGCTGCTATTGGGCGAAGTGCCGGGGCAGGATCTCCTGTCATCTCACCTTGC
TCCTGCCGAGAAAGTATCCATCATGGCTGATGCAATGCGGCGGCTGCATACGCTTGATCCGGCTACCTGCCCATTCG
ACCACCAAGCGAAACATCGCATCGAGCGAGCACGTACTCGGATGGAAGCCGGTCTTGTCGATCAGGATGATCTGGAC
GAAGAGCATCAGGGGCTCGCGCCAGCCGAACTGTTCGCCAGGCTCAAGGCGCGCATGCCCGACGGCGAGGATCTCGT
CGTGACCCATGGCGATGCCTGCTTGCCGAATATCATGGTGGAAAATGGCCGCTTTTCTGGATTCATCGACTGTGGCC
GGCTGGGTGTGGCGGACCGCTATCAGGACATAGCGTTGGCTACCCGTGATATTGCTGAAGAGCTTGGCGGCGAATGG
GCTGACCGCTTCCTCGTGCTTTACGGTATCGCCGCTCCCGATTCGCAGCGCATCGCCTTCTATCGCCTTCTTGACGA
GTTCTTCTGAGCGGGACTCTGGGGTTCGAAATGACCGACCAAGCGACGCCCAACCTGCCATCACGAGATTTCGATTC
CACCGCCGCCTTCTATGAAAGGTTGGGCTTCGGAATCGTTTTCCGGGACGCCGGCTGGATGATCCTCCAGCGCGGGG
ATCTCATGCTGGAGTTCTTCGCCCACCCCAACTTGTTTATTGCAGCTTATAATGGTTACAAATAAAGCAATAGCATC
ACAAATTTCACAAATAAAGCATTTTTTTCACTGCATTCTAGTTGTGGTTTGTCCAAACTCATCAATGTATCTTATCA
TGTCTGTATACCGTCGACCTCTAGCTAGAGCTTGGCGTAATCATGGTCATAGCTGTTTCCTGTGTGAAATTGTTATC
CGCTCACAATTCCACACAACATACGAGCCGGAAGCATAAAGTGTAAAGCCTGGGGTGCCTAATGAGTGAGCTAACTC
ACATTAATTGCGTTGCGCTCACTGCCCGCTTTCCAGTCGGGAAACCTGTCGTGCCAGCTGCATTAATGAATCGGCCA
ACGCGCGGGGAGAGGCGGTTTGCGTATTGGGCGCTCTTCCGCTTCCTCGCTCACTGACTCGCTGCGCTCGGTCGTTC
GGCTGCGGCGAGCGGTATCAGCTCACTCAAAGGCGGTAATACGGTTATCCACAGAATCAGGGGATAACGCAGGAAAG
AACATGTGAGCAAAAGGCCAGCAAAAGGCCAGGAACCGTAAAAAGGCCGCGTTGCTGGCGTTTTTCCATAGGCTCCG
CCCCCCTGACGAGCATCACAAAAATCGACGCTCAAGTCAGAGGTGGCGAAACCCGACAGGACTATAAAGATACCAGG
CGTTTCCCCCTGGAAGCTCCCTCGTGCGCTCTCCTGTTCCGACCCTGCCGCTTACCGGATACCTGTCCGCCTTTCTC
CCTTCGGGAAGCGTGGCGCTTTCTCATAGCTCACGCTGTAGGTATCTCAGTTCGGTGTAGGTCGTTCGCTCCAAGCT
GGGCTGTGTGCACGAACCCCCCGTTCAGCCCGACCGCTGCGCCTTATCCGGTAACTATCGTCTTGAGTCCAACCCGG
TAAGACACGACTTATCGCCACTGGCAGCAGCCACTGGTAACAGGATTAGCAGAGCGAGGTATGTAGGCGGTGCTACA
GAGTTCTTGAAGTGGTGGCCTAACTACGGCTACACTAGAAGAACAGTATTTGGTATCTGCGCTCTGCTGAAGCCAGT
TACCTTCGGAAAAAGAGTTGGTAGCTCTTGATCCGGCAAACAAACCACCGCTGGTAGCGGTGGTTTTTTTGTTTGCA
AGCAGCAGATTACGCGCAGAAAAAAAGGATCTCAAGAAGATCCTTTGATCTTTTCTACGGGGTCTGACGCTCAGTGG
AACGAAAACTCACGTTAAGGGATTTTGGTCATGAGATTATCAAAAAGGATCTTCACCTAGATCCTTTTAAATTAAAA
ATGAAGTTTTAAATCAATCTAAAGTATATATGAGTAAACTTGGTCTGACAGTTACCAATGCTTAATCAGTGAGGCAC
CTATCTCAGCGATCTGTCTATTTCGTTCATCCATAGTTGCCTGACTCCCCGTCGTGTAGATAACTACGATACGGGAG
GGCTTACCATCTGGCCCCAGTGCTGCAATGATACCGCGAGACCCACGCTCACCGGCTCCAGATTTATCAGCAATAAA
CCAGCCAGCCGGAAGGGCCGAGCGCAGAAGTGGTCCTGCAACTTTATCCGCCTCCATCCAGTCTATTAATTGTTGCC
GGGAAGCTAGAGTAAGTAGTTCGCCAGTTAATAGTTTGCGCAACGTTGTTGCCATTGCTACAGGCATCGTGGTGTCA
CGCTCGTCGTTTGGTATGGCTTCATTCAGCTCCGGTTCCCAACGATCAAGGCGAGTTACATGATCCCCCATGTTGTG
CAAAAAAGCGGTTAGCTCCTTCGGTCCTCCGATCGTTGTCAGAAGTAAGTTGGCCGCAGTGTTATCACTCATGGTTA
TGGCAGCACTGCATAATTCTCTTACTGTCATGCCATCCGTAAGATGCTTTTCTGTGACTGGTGAGTACTCAACCAAG
TCATTCTGAGAATAGTGTATGCGGCGACCGAGTTGCTCTTGCCCGGCGTCAATACGGGATAATACCGCGCCACATAG
CAGAACTTTAAAAGTGCTCATCATTGGAAAACGTTCTTCGGGGCGAAAACTCTCAAGGATCTTACCGCTGTTGAGAT
CCAGTTCGATGTAACCCACTCGTGCACCCAACTGATCTTCAGCATCTTTTACTTTCACCAGCGTTTCTGGGTGAGCA
AAAACAGGAAGGCAAAATGCCGCAAAAAAGGGAATAAGGGCGACACGGAAATGTTGAATACTCATACTCTTCCTTTT
TCAATATTATTGAAGCATTTATCAGGGTTATTGTCTCATGAGCGGATACATATTTGAATGTATTTAGAAAAATAAAC
AAATAGGGGTTCCGCGCACATTTCCCCGAAAAGTGCCACCTGACGTC

>ALX4 Y25D in ALX4_OHu22706C_pcDNA3.1(+)-N-HA

GACGGATCGGGAGATCTCCCGATCCCCTATGGTGCACTCTCAGTACAATCTGCTCTGATGCCGCATAGTTAAGCCAG
TATCTGCTCCCTGCTTGTGTGTTGGAGGTCGCTGAGTAGTGCGCGAGCAAAATTTAAGCTACAACAAGGCAAGGCTT
GACCGACAATTGCATGAAGAATCTGCTTAGGGTTAGGCGTTTTGCGCTGCTTCGCGATGTACGGGCCAGATATACGC
GTTGACATTGATTATTGACTAGTTATTAATAGTAATCAATTACGGGGTCATTAGTTCATAGCCCATATATGGAGTTC
CGCGTTACATAACTTACGGTAAATGGCCCGCCTGGCTGACCGCCCAACGACCCCCGCCCATTGACGTCAATAATGAC
GTATGTTCCCATAGTAACGCCAATAGGGACTTTCCATTGACGTCAATGGGTGGAGTATTTACGGTAAACTGCCCACT
TGGCAGTACATCAAGTGTATCATATGCCAAGTACGCCCCCTATTGACGTCAATGACGGTAAATGGCCCGCCTGGCAT
TATGCCCAGTACATGACCTTATGGGACTTTCCTACTTGGCAGTACATCTACGTATTAGTCATCGCTATTACCATGGT
GATGCGGTTTTGGCAGTACATCAATGGGCGTGGATAGCGGTTTGACTCACGGGGATTTCCAAGTCTCCACCCCATTG
ACGTCAATGGGAGTTTGTTTTGGCACCAAAATCAACGGGACTTTCCAAAATGTCGTAACAACTCCGCCCCATTGACG
CAAATGGGCGGTAGGCGTGTACGGTGGGAGGTCTATATAAGCAGAGCTCTCTGGCTAACTAGAGAACCCACTGCTTA
CTGGCTTATCGAAATTAATACGACTCACTATAGGGAGACCCAAGCTGGCTAGCGTTTAAACTTAAGCTTGCCACCAT
GTACCCATACGATGTTCCAGATTACGCTGGCTCTGGAAATGCTGAGACTTGCGTCTCTTACTGCGAGTCGCCGGCCG
CTGCCATGGACGCCTACTACAGCCCGGTGTCGCAGAGTCGGGAGGGCTCGTCGCCTTTTAGGGCATTTCCCGGAGGC
GACAAGTTCGGCACAACTTTCCTGTCGGCCGCCGCCAAAGCACAGGGATTCGGGGACGCCAAGAGCCGGGCCCGTTA
CGGCGCTGGGCAGCAGGACCTGGCGACACCCCTGGAGAGTGGAGCTGGGGCGCGGGGCTCCTTTAACAAGTTCCAGC
CCCAGCCGTCGACCCCGCAGCCCCAGCCGCCGCCGCAGCCGCAGCCGCAGCAGCAGCAGCCGCAGCCCCAGCCGCCC
GCGCAACCGCATCTTTACTTGCAGCGAGGCGCCTGCAAGACGCCCCCGGACGGCAGCCTCAAACTCCAGGAAGGCAG
CAGCGGCCACAGCGCGGCCTTGCAGGTTCCCTGCTACGCTAAAGAGAGCTCCCTGGGTGAGCCAGAGTTACCCCCTG
ACTCTGACACTGTGGGGATGGACAGCAGCTACCTGAGTGTCAAGGAGGCTGGGGTGAAGGGGCCCCAGGACCGGGCC
AGCTCAGACCTCCCCAGCCCATTGGAGAAGGCCGACTCAGAGAGCAACAAGGGCAAGAAGCGGCGGAACCGGACCAC
CTTCACCAGCTACCAGCTGGAGGAGCTGGAGAAGGTCTTCCAGAAGACCCACGACCCAGACGTGTATGCGCGGGAAC
AGCTGGCCATGAGGACAGACCTCACTGAGGCCCGCGTGCAGGTCTGGTTCCAGAACCGAAGGGCCAAGTGGAGGAAG
CGGGAGCGTTTTGGGCAGATGCAGCAGGTTCGAACCCACTTCTCCACTGCATATGAGCTGCCCCTCCTCACCCGAGC
TGAGAACTACGCCCAGATTCAGAACCCGTCCTGGCTCGGCAACAACGGGGCTGCCTCACCAGTGCCAGCCTGCGTGG
TCCCCTGCGACCCGGTGCCTGCCTGCATGTCCCCTCATGCCCACCCCCCTGGCTCTGGGGCCAGCAGCGTCACCGAC
TTCCTGAGTGTGTCTGGGGCTGGCAGTCACGTGGGCCAGACGCACATGGGCAGCCTGTTTGGAGCAGCCAGCCTCAG
CCCAGGCCTCAATGGCTACGAGCTCAACGGCGAGCCGGACCGCAAGACCTCGAGCATCGCGGCCCTCCGCATGAAGG
CCAAGGAGCACAGTGCGGCCATTTCCTGGGCCACATGAGGATCCGAATTCTGCAGATATCCAGCACAGTGGCGGCCG
CTCGAGTCTAGAGGGCCCGTTTAAACCCGCTGATCAGCCTCGACTGTGCCTTCTAGTTGCCAGCCATCTGTTGTTTG
CCCCTCCCCCGTGCCTTCCTTGACCCTGGAAGGTGCCACTCCCACTGTCCTTTCCTAATAAAATGAGGAAATTGCAT
CGCATTGTCTGAGTAGGTGTCATTCTATTCTGGGGGGTGGGGTGGGGCAGGACAGCAAGGGGGAGGATTGGGAAGAC
AATAGCAGGCATGCTGGGGATGCGGTGGGCTCTATGGCTTCTGAGGCGGAAAGAACCAGCTGGGGCTCTAGGGGGTA
TCCCCACGCGCCCTGTAGCGGCGCATTAAGCGCGGCGGGTGTGGTGGTTACGCGCAGCGTGACCGCTACACTTGCCA
GCGCCCTAGCGCCCGCTCCTTTCGCTTTCTTCCCTTCCTTTCTCGCCACGTTCGCCGGCTTTCCCCGTCAAGCTCTA
AATCGGGGGCTCCCTTTAGGGTTCCGATTTAGTGCTTTACGGCACCTCGACCCCAAAAAACTTGATTAGGGTGATGG
TTCACGTAGTGGGCCATCGCCCTGATAGACGGTTTTTCGCCCTTTGACGTTGGAGTCCACGTTCTTTAATAGTGGAC
TCTTGTTCCAAACTGGAACAACACTCAACCCTATCTCGGTCTATTCTTTTGATTTATAAGGGATTTTGCCGATTTCG
GCCTATTGGTTAAAAAATGAGCTGATTTAACAAAAATTTAACGCGAATTAATTCTGTGGAATGTGTGTCAGTTAGGG
TGTGGAAAGTCCCCAGGCTCCCCAGCAGGCAGAAGTATGCAAAGCATGCATCTCAATTAGTCAGCAACCAGGTGTGG
AAAGTCCCCAGGCTCCCCAGCAGGCAGAAGTATGCAAAGCATGCATCTCAATTAGTCAGCAACCATAGTCCCGCCCC
TAACTCCGCCCATCCCGCCCCTAACTCCGCCCAGTTCCGCCCATTCTCCGCCCCATGGCTGACTAATTTTTTTTATT

TATGCAGAGGCCGAGGCCGCCTCTGCCTCTGAGCTATTCCAGAAGTAGTGAGGAGGCTTTTTTGGAGGCCTAGGCTT
TTGCAAAAAGCTCCCGGGAGCTTGTATATCCATTTTCGGATCTGATCAAGAGACAGGATGAGGATCGTTTCGCATGA
TTGAACAAGATGGATTGCACGCAGGTTCTCCGGCCGCTTGGGTGGAGAGGCTATTCGGCTATGACTGGGCACAACAG
ACAATCGGCTGCTCTGATGCCGCCGTGTTCCGGCTGTCAGCGCAGGGGCGCCCGGTTCTTTTTGTCAAGACCGACCT
GTCCGGTGCCCTGAATGAACTGCAGGACGAGGCAGCGCGGCTATCGTGGCTGGCCACGACGGGCGTTCCTTGCGCAG
CTGTGCTCGACGTTGTCACTGAAGCGGGAAGGGACTGGCTGCTATTGGGCGAAGTGCCGGGGCAGGATCTCCTGTCA
TCTCACCTTGCTCCTGCCGAGAAAGTATCCATCATGGCTGATGCAATGCGGCGGCTGCATACGCTTGATCCGGCTAC
CTGCCCATTCGACCACCAAGCGAAACATCGCATCGAGCGAGCACGTACTCGGATGGAAGCCGGTCTTGTCGATCAGG
ATGATCTGGACGAAGAGCATCAGGGGCTCGCGCCAGCCGAACTGTTCGCCAGGCTCAAGGCGCGCATGCCCGACGGC
GAGGATCTCGTCGTGACCCATGGCGATGCCTGCTTGCCGAATATCATGGTGGAAAATGGCCGCTTTTCTGGATTCAT
CGACTGTGGCCGGCTGGGTGTGGCGGACCGCTATCAGGACATAGCGTTGGCTACCCGTGATATTGCTGAAGAGCTTG
GCGGCGAATGGGCTGACCGCTTCCTCGTGCTTTACGGTATCGCCGCTCCCGATTCGCAGCGCATCGCCTTCTATCGC
CTTCTTGACGAGTTCTTCTGAGCGGGACTCTGGGGTTCGAAATGACCGACCAAGCGACGCCCAACCTGCCATCACGA
GATTTCGATTCCACCGCCGCCTTCTATGAAAGGTTGGGCTTCGGAATCGTTTTCCGGGACGCCGGCTGGATGATCCT
CCAGCGCGGGGATCTCATGCTGGAGTTCTTCGCCCACCCCAACTTGTTTATTGCAGCTTATAATGGTTACAAATAAA
GCAATAGCATCACAAATTTCACAAATAAAGCATTTTTTTCACTGCATTCTAGTTGTGGTTTGTCCAAACTCATCAAT
GTATCTTATCATGTCTGTATACCGTCGACCTCTAGCTAGAGCTTGGCGTAATCATGGTCATAGCTGTTTCCTGTGTG
AAATTGTTATCCGCTCACAATTCCACACAACATACGAGCCGGAAGCATAAAGTGTAAAGCCTGGGGTGCCTAATGAG
TGAGCTAACTCACATTAATTGCGTTGCGCTCACTGCCCGCTTTCCAGTCGGGAAACCTGTCGTGCCAGCTGCATTAA
TGAATCGGCCAACGCGCGGGGAGAGGCGGTTTGCGTATTGGGCGCTCTTCCGCTTCCTCGCTCACTGACTCGCTGCG
CTCGGTCGTTCGGCTGCGGCGAGCGGTATCAGCTCACTCAAAGGCGGTAATACGGTTATCCACAGAATCAGGGGATA
ACGCAGGAAAGAACATGTGAGCAAAAGGCCAGCAAAAGGCCAGGAACCGTAAAAAGGCCGCGTTGCTGGCGTTTTTC
CATAGGCTCCGCCCCCCTGACGAGCATCACAAAAATCGACGCTCAAGTCAGAGGTGGCGAAACCCGACAGGACTATA
AAGATACCAGGCGTTTCCCCCTGGAAGCTCCCTCGTGCGCTCTCCTGTTCCGACCCTGCCGCTTACCGGATACCTGT
CCGCCTTTCTCCCTTCGGGAAGCGTGGCGCTTTCTCATAGCTCACGCTGTAGGTATCTCAGTTCGGTGTAGGTCGTT
CGCTCCAAGCTGGGCTGTGTGCACGAACCCCCCGTTCAGCCCGACCGCTGCGCCTTATCCGGTAACTATCGTCTTGA
GTCCAACCCGGTAAGACACGACTTATCGCCACTGGCAGCAGCCACTGGTAACAGGATTAGCAGAGCGAGGTATGTAG
GCGGTGCTACAGAGTTCTTGAAGTGGTGGCCTAACTACGGCTACACTAGAAGAACAGTATTTGGTATCTGCGCTCTG
CTGAAGCCAGTTACCTTCGGAAAAAGAGTTGGTAGCTCTTGATCCGGCAAACAAACCACCGCTGGTAGCGGTGGTTT
TTTTGTTTGCAAGCAGCAGATTACGCGCAGAAAAAAAGGATCTCAAGAAGATCCTTTGATCTTTTCTACGGGGTCTG
ACGCTCAGTGGAACGAAAACTCACGTTAAGGGATTTTGGTCATGAGATTATCAAAAAGGATCTTCACCTAGATCCTT
TTAAATTAAAAATGAAGTTTTAAATCAATCTAAAGTATATATGAGTAAACTTGGTCTGACAGTTACCAATGCTTAAT
CAGTGAGGCACCTATCTCAGCGATCTGTCTATTTCGTTCATCCATAGTTGCCTGACTCCCCGTCGTGTAGATAACTA
CGATACGGGAGGGCTTACCATCTGGCCCCAGTGCTGCAATGATACCGCGAGACCCACGCTCACCGGCTCCAGATTTA
TCAGCAATAAACCAGCCAGCCGGAAGGGCCGAGCGCAGAAGTGGTCCTGCAACTTTATCCGCCTCCATCCAGTCTAT
TAATTGTTGCCGGGAAGCTAGAGTAAGTAGTTCGCCAGTTAATAGTTTGCGCAACGTTGTTGCCATTGCTACAGGCA
TCGTGGTGTCACGCTCGTCGTTTGGTATGGCTTCATTCAGCTCCGGTTCCCAACGATCAAGGCGAGTTACATGATCC
CCCATGTTGTGCAAAAAAGCGGTTAGCTCCTTCGGTCCTCCGATCGTTGTCAGAAGTAAGTTGGCCGCAGTGTTATC
ACTCATGGTTATGGCAGCACTGCATAATTCTCTTACTGTCATGCCATCCGTAAGATGCTTTTCTGTGACTGGTGAGT
ACTCAACCAAGTCATTCTGAGAATAGTGTATGCGGCGACCGAGTTGCTCTTGCCCGGCGTCAATACGGGATAATACC
GCGCCACATAGCAGAACTTTAAAAGTGCTCATCATTGGAAAACGTTCTTCGGGGCGAAAACTCTCAAGGATCTTACC
GCTGTTGAGATCCAGTTCGATGTAACCCACTCGTGCACCCAACTGATCTTCAGCATCTTTTACTTTCACCAGCGTTT
CTGGGTGAGCAAAAACAGGAAGGCAAAATGCCGCAAAAAAGGGAATAAGGGCGACACGGAAATGTTGAATACTCATA

CTCTTCCTTTTTCAATATTATTGAAGCATTTATCAGGGTTATTGTCTCATGAGCGGATACATATTTGAATGTATTTA
GAAAAATAAACAAATAGGGGTTCCGCGCACATTTCCCCGAAAAGTGCCACCTGACGTC

>PHOX2B E42Q in pUC57

TCGCGCGTTTCGGTGATGACGGTGAAAACCTCTGACACATGCAGCTCCCGGAGACGGTCACAGCTTGTCTGTAAGCG
GATGCCGGGAGCAGACAAGCCCGTCAGGGCGCGTCAGCGGGTGTTGGCGGGTGTCGGGGCTGGCTTAACTATGCGGC
ATCAGAGCAGATTGTACTGAGAGTGCACCATATGCGGTGTGAAATACCGCACAGATGCGTAAGGAGAAAATACCGCA
TCAGGCGCCATTCGCCATTCAGGCTGCGCAACTGTTGGGAAGGGCGATCGGTGCGGGCCTCTTCGCTATTACGCCAG
CTGGCGAAAGGGGGATGTGCTGCAAGGCGATTAAGTTGGGTAACGCCAGGGTTTTCCCAGTCACGACGTTGTAAAAC
GACGGCCAGTGAATTCGAGCTCGGTACCTCGCGAATGCATCTAGATCATATGAACGAGAAGCGCAAGCAGCGGCGCA
TCCGCACCACTTTCACCAGTGCCCAGCTCAAAGAGCTGGAAAGGGTCTTCGCGGAGACTCACTACCCCGACATCTAC
ACTCGGGAGGAGCTGGCCCTGAAGATCGACCTCACACAGGCGCGAGTCCAGGTGTGGTTCCAGAACCGCCGCGCCAA
GTTTCGCAAGCAGGAGCGCTAACTCGAGATCGGATCCCGGGCCCGTCGACTGCAGAGGCCTGCATGCAAGCTTGGCG
TAATCATGGTCATAGCTGTTTCCTGTGTGAAATTGTTATCCGCTCACAATTCCACACAACATACGAGCCGGAAGCAT
AAAGTGTAAAGCCTGGGGTGCCTAATGAGTGAGCTAACTCACATTAATTGCGTTGCGCTCACTGCCCGCTTTCCAGT
CGGGAAACCTGTCGTGCCAGCTGCATTAATGAATCGGCCAACGCGCGGGGAGAGGCGGTTTGCGTATTGGGCGCTCT
TCCGCTTCCTCGCTCACTGACTCGCTGCGCTCGGTCGTTCGGCTGCGGCGAGCGGTATCAGCTCACTCAAAGGCGGT
AATACGGTTATCCACAGAATCAGGGGATAACGCAGGAAAGAACATGTGAGCAAAAGGCCAGCAAAAGGCCAGGAACC
GTAAAAAGGCCGCGTTGCTGGCGTTTTTCCATAGGCTCCGCCCCCCTGACGAGCATCACAAAAATCGACGCTCAAGT
CAGAGGTGGCGAAACCCGACAGGACTATAAAGATACCAGGCGTTTCCCCCTGGAAGCTCCCTCGTGCGCTCTCCTGT
TCCGACCCTGCCGCTTACCGGATACCTGTCCGCCTTTCTCCCTTCGGGAAGCGTGGCGCTTTCTCATAGCTCACGCT
GTAGGTATCTCAGTTCGGTGTAGGTCGTTCGCTCCAAGCTGGGCTGTGTGCACGAACCCCCCGTTCAGCCCGACCGC
TGCGCCTTATCCGGTAACTATCGTCTTGAGTCCAACCCGGTAAGACACGACTTATCGCCACTGGCAGCAGCCACTGG
TAACAGGATTAGCAGAGCGAGGTATGTAGGCGGTGCTACAGAGTTCTTGAAGTGGTGGCCTAACTACGGCTACACTA
GAAGAACAGTATTTGGTATCTGCGCTCTGCTGAAGCCAGTTACCTTCGGAAAAAGAGTTGGTAGCTCTTGATCCGGC
AAACAAACCACCGCTGGTAGCGGTGGTTTTTTTGTTTGCAAGCAGCAGATTACGCGCAGAAAAAAAGGATCTCAAGA
AGATCCTTTGATCTTTTCTACGGGGTCTGACGCTCAGTGGAACGAAAACTCACGTTAAGGGATTTTGGTCATGAGAT
TATCAAAAAGGATCTTCACCTAGATCCTTTTAAATTAAAAATGAAGTTTTAAATCAATCTAAAGTATATATGAGTAA
ACTTGGTCTGACAGTTACCAATGCTTAATCAGTGAGGCACCTATCTCAGCGATCTGTCTATTTCGTTCATCCATAGT
TGCCTGACTCCCCGTCGTGTAGATAACTACGATACGGGAGGGCTTACCATCTGGCCCCAGTGCTGCAATGATACCGC
GAGACCCACGCTCACCGGCTCCAGATTTATCAGCAATAAACCAGCCAGCCGGAAGGGCCGAGCGCAGAAGTGGTCCT
GCAACTTTATCCGCCTCCATCCAGTCTATTAATTGTTGCCGGGAAGCTAGAGTAAGTAGTTCGCCAGTTAATAGTTT
GCGCAACGTTGTTGCCATTGCTACAGGCATCGTGGTGTCACGCTCGTCGTTTGGTATGGCTTCATTCAGCTCCGGTT
CCCAACGATCAAGGCGAGTTACATGATCCCCCATGTTGTGCAAAAAAGCGGTTAGCTCCTTCGGTCCTCCGATCGTT
GTCAGAAGTAAGTTGGCCGCAGTGTTATCACTCATGGTTATGGCAGCACTGCATAATTCTCTTACTGTCATGCCATC
CGTAAGATGCTTTTCTGTGACTGGTGAGTACTCAACCAAGTCATTCTGAGAATAGTGTATGCGGCGACCGAGTTGCT
CTTGCCCGGCGTCAATACGGGATAATACCGCGCCACATAGCAGAACTTTAAAAGTGCTCATCATTGGAAAACGTTCT
TCGGGGCGAAAACTCTCAAGGATCTTACCGCTGTTGAGATCCAGTTCGATGTAACCCACTCGTGCACCCAACTGATC
TTCAGCATCTTTTACTTTCACCAGCGTTTCTGGGTGAGCAAAAACAGGAAGGCAAAATGCCGCAAAAAAGGGAATAA
GGGCGACACGGAAATGTTGAATACTCATACTCTTCCTTTTTCAATATTATTGAAGCATTTATCAGGGTTATTGTCTC
ATGAGCGGATACATATTTGAATGTATTTAGAAAAATAAACAAATAGGGGTTCCGCGCACATTTCCCCGAAAAGTGCC
ACCTGACGTCTAAGAAACCATTATTATCATGACATTAACCTATAAAAATAGGCGTATCACGAGGCCCTTTCGTC

>PHOX2B A43E in pUC57

TCGCGCGTTTCGGTGATGACGGTGAAAACCTCTGACACATGCAGCTCCCGGAGACGGTCACAGCTTGTCTGTAAGCG
GATGCCGGGAGCAGACAAGCCCGTCAGGGCGCGTCAGCGGGTGTTGGCGGGTGTCGGGGCTGGCTTAACTATGCGGC
ATCAGAGCAGATTGTACTGAGAGTGCACCATATGCGGTGTGAAATACCGCACAGATGCGTAAGGAGAAAATACCGCA
TCAGGCGCCATTCGCCATTCAGGCTGCGCAACTGTTGGGAAGGGCGATCGGTGCGGGCCTCTTCGCTATTACGCCAG
CTGGCGAAAGGGGGATGTGCTGCAAGGCGATTAAGTTGGGTAACGCCAGGGTTTTCCCAGTCACGACGTTGTAAAAC
GACGGCCAGTGAATTCGAGCTCGGTACCTCGCGAATGCATCTAGATCATATGAACGAGAAGCGCAAGCAGCGGCGCA
TCCGCACCACTTTCACCAGTGCCCAGCTCAAAGAGCTGGAAAGGGTCTTCGCGGAGACTCACTACCCCGACATCTAC
ACTCGGGAGGAGCTGGCCCTGAAGATCGACCTCACAGAGGAGCGAGTCCAGGTGTGGTTCCAGAACCGCCGCGCCAA
GTTTCGCAAGCAGGAGCGCTAACTCGAGATCGGATCCCGGGCCCGTCGACTGCAGAGGCCTGCATGCAAGCTTGGCG
TAATCATGGTCATAGCTGTTTCCTGTGTGAAATTGTTATCCGCTCACAATTCCACACAACATACGAGCCGGAAGCAT
AAAGTGTAAAGCCTGGGGTGCCTAATGAGTGAGCTAACTCACATTAATTGCGTTGCGCTCACTGCCCGCTTTCCAGT
CGGGAAACCTGTCGTGCCAGCTGCATTAATGAATCGGCCAACGCGCGGGGAGAGGCGGTTTGCGTATTGGGCGCTCT
TCCGCTTCCTCGCTCACTGACTCGCTGCGCTCGGTCGTTCGGCTGCGGCGAGCGGTATCAGCTCACTCAAAGGCGGT
AATACGGTTATCCACAGAATCAGGGGATAACGCAGGAAAGAACATGTGAGCAAAAGGCCAGCAAAAGGCCAGGAACC
GTAAAAAGGCCGCGTTGCTGGCGTTTTTCCATAGGCTCCGCCCCCCTGACGAGCATCACAAAAATCGACGCTCAAGT
CAGAGGTGGCGAAACCCGACAGGACTATAAAGATACCAGGCGTTTCCCCCTGGAAGCTCCCTCGTGCGCTCTCCTGT
TCCGACCCTGCCGCTTACCGGATACCTGTCCGCCTTTCTCCCTTCGGGAAGCGTGGCGCTTTCTCATAGCTCACGCT
GTAGGTATCTCAGTTCGGTGTAGGTCGTTCGCTCCAAGCTGGGCTGTGTGCACGAACCCCCCGTTCAGCCCGACCGC
TGCGCCTTATCCGGTAACTATCGTCTTGAGTCCAACCCGGTAAGACACGACTTATCGCCACTGGCAGCAGCCACTGG
TAACAGGATTAGCAGAGCGAGGTATGTAGGCGGTGCTACAGAGTTCTTGAAGTGGTGGCCTAACTACGGCTACACTA
GAAGAACAGTATTTGGTATCTGCGCTCTGCTGAAGCCAGTTACCTTCGGAAAAAGAGTTGGTAGCTCTTGATCCGGC
AAACAAACCACCGCTGGTAGCGGTGGTTTTTTTGTTTGCAAGCAGCAGATTACGCGCAGAAAAAAAGGATCTCAAGA
AGATCCTTTGATCTTTTCTACGGGGTCTGACGCTCAGTGGAACGAAAACTCACGTTAAGGGATTTTGGTCATGAGAT
TATCAAAAAGGATCTTCACCTAGATCCTTTTAAATTAAAAATGAAGTTTTAAATCAATCTAAAGTATATATGAGTAA
ACTTGGTCTGACAGTTACCAATGCTTAATCAGTGAGGCACCTATCTCAGCGATCTGTCTATTTCGTTCATCCATAGT
TGCCTGACTCCCCGTCGTGTAGATAACTACGATACGGGAGGGCTTACCATCTGGCCCCAGTGCTGCAATGATACCGC
GAGACCCACGCTCACCGGCTCCAGATTTATCAGCAATAAACCAGCCAGCCGGAAGGGCCGAGCGCAGAAGTGGTCCT
GCAACTTTATCCGCCTCCATCCAGTCTATTAATTGTTGCCGGGAAGCTAGAGTAAGTAGTTCGCCAGTTAATAGTTT
GCGCAACGTTGTTGCCATTGCTACAGGCATCGTGGTGTCACGCTCGTCGTTTGGTATGGCTTCATTCAGCTCCGGTT
CCCAACGATCAAGGCGAGTTACATGATCCCCCATGTTGTGCAAAAAAGCGGTTAGCTCCTTCGGTCCTCCGATCGTT
GTCAGAAGTAAGTTGGCCGCAGTGTTATCACTCATGGTTATGGCAGCACTGCATAATTCTCTTACTGTCATGCCATC
CGTAAGATGCTTTTCTGTGACTGGTGAGTACTCAACCAAGTCATTCTGAGAATAGTGTATGCGGCGACCGAGTTGCT
CTTGCCCGGCGTCAATACGGGATAATACCGCGCCACATAGCAGAACTTTAAAAGTGCTCATCATTGGAAAACGTTCT
TCGGGGCGAAAACTCTCAAGGATCTTACCGCTGTTGAGATCCAGTTCGATGTAACCCACTCGTGCACCCAACTGATC
TTCAGCATCTTTTACTTTCACCAGCGTTTCTGGGTGAGCAAAAACAGGAAGGCAAAATGCCGCAAAAAAGGGAATAA
GGGCGACACGGAAATGTTGAATACTCATACTCTTCCTTTTTCAATATTATTGAAGCATTTATCAGGGTTATTGTCTC
ATGAGCGGATACATATTTGAATGTATTTAGAAAAATAAACAAATAGGGGTTCCGCGCACATTTCCCCGAAAAGTGCC
ACCTGACGTCTAAGAAACCATTATTATCATGACATTAACCTATAAAAATAGGCGTATCACGAGGCCCTTTCGTC

>ARX A43V in Arx_OMu17806D_pcDNA3.1+/C-(K)-DYK

GACGGATCGGGAGATCTCCCGATCCCCTATGGTGCACTCTCAGTACAATCTGCTCTGATGCCGCATAGTTAAGCCAG
TATCTGCTCCCTGCTTGTGTGTTGGAGGTCGCTGAGTAGTGCGCGAGCAAAATTTAAGCTACAACAAGGCAAGGCTT
GACCGACAATTGCATGAAGAATCTGCTTAGGGTTAGGCGTTTTGCGCTGCTTCGCGATGTACGGGCCAGATATACGC
GTTGACATTGATTATTGACTAGTTATTAATAGTAATCAATTACGGGGTCATTAGTTCATAGCCCATATATGGAGTTC
CGCGTTACATAACTTACGGTAAATGGCCCGCCTGGCTGACCGCCCAACGACCCCCGCCCATTGACGTCAATAATGAC
GTATGTTCCCATAGTAACGCCAATAGGGACTTTCCATTGACGTCAATGGGTGGAGTATTTACGGTAAACTGCCCACT
TGGCAGTACATCAAGTGTATCATATGCCAAGTACGCCCCCTATTGACGTCAATGACGGTAAATGGCCCGCCTGGCAT
TATGCCCAGTACATGACCTTATGGGACTTTCCTACTTGGCAGTACATCTACGTATTAGTCATCGCTATTACCATGGT
GATGCGGTTTTGGCAGTACATCAATGGGCGTGGATAGCGGTTTGACTCACGGGGATTTCCAAGTCTCCACCCCATTG
ACGTCAATGGGAGTTTGTTTTGGCACCAAAATCAACGGGACTTTCCAAAATGTCGTAACAACTCCGCCCCATTGACG
CAAATGGGCGGTAGGCGTGTACGGTGGGAGGTCTATATAAGCAGAGCTCTCTGGCTAACTAGAGAACCCACTGCTTA
CTGGCTTATCGAAATTAATACGACTCACTATAGGGAGACCCAAGCTGGCTAGCGTTTAAACTTAAGCTTGGTACCGA
GCTCGGATCCGCCACCATGAGCAATCAGTACCAGGAAGAGGGCTGCTCTGAGAGGCCGGAGTGCAAGAGTAAATCTC
CAACTTTGCTCTCCTCCTACTGCATCGACAGCATCCTGGGCCGGAGAAGCCCATGCAAAATGAGGCTTCTGGGAGCC
GCGCAGAGCTTGCCTGCCCCGCTGGCCAGCCGCGCTGACCAGGAAAAAGCCATGCAAGGCTCCCCCAAGAGCAGCAG
CGCCCCGTTCGAGGCTGAGCTGCACCTGCCACCCAAGCTGCGGCGCCTCTACGGCCCGGGCGGGGGCCGCCTCCTCC
AGGGCGCGGCGGCGGCGGCTGCAGCGGCGGCGGCGGCCGCAGCGGCGGCCACGGCCACTGGCACCGCAGGTCCCCGT
GGAGAGGTTCCTCCACCGCCACCACCAGCTGCACGGCCAGGGGAGCGTCAGGACAGCGCAGGGGCCGTGGCTGCCGC
CGCTGCCGCCGCTGCCTGGGACACGCTCAAGATCAGCCAGGCGCCGCAGGTGAGCATCAGTCGTAGCAAGTCGTACC
GCGAGAACGGGGCGCCCTTCGTGCCCCCGCCGCCAGCCCTGGACGAGCTGAGTGGCCCGGGGGGCGTCGCGCACCCC
GAGGAGCGCCTCAGCGCGGCCAGCGGACCCGGCAGTGCCCCCGCGGCGGGTGGTGGCACGGGCGCCGAGGACGACGA
GGAGGAGCTGTTGGAGGATGAGGAAGATGAAGAGGAGGAAGAGGAGCTGCTGGAGGACGACGACGAGGAGCTGCTGG
AGGACGATGCCCGCGCGCTGCTCAAGGAGCCCCGGCGCTGCTCGGTGGCCACCACCGGCACAGTGGCCGCCGCCGCC
GCCGCTGCTGCCGCCGCCGTGGCCACCGAGGGCGGGGAGCTGTCGCCCAAGGAAGAGCTGTTGCTGCACCCGGAGGA
CGCCGAGGGCAAGGATGGTGAGGACAGCGTGTGCCTGTCGGCCGGCAGCGACTCGGAGGAGGGGCTGCTGAAGCGCA
AACAGAGGCGCTACCGCACCACGTTCACCAGTTACCAGCTGGAGGAACTGGAGCGGGCTTTCCAGAAGACGCACTAC
CCTGACGTCTTCACCAGGGAAGAGCTGGCCATGAGGCTGGACCTGACAGAGGTCCGAGTCCAGGTGTGGTTCCAGAA
CCGTCGGGCCAAGTGGCGCAAGCGGGAGAAGGCTGGCGCACAGACCCACCCCCCGGGGCTGCCCTTTCCAGGGCCGC
TCTCGGCTACCCACCCTCTCAGCCCCTACCTGGACGCCAGCCCCTTTCCCCCGCATCACCCTGCGCTTGATTCGGCC
TGGACCGCCGCGGCGGCGGCGGCTGCAGCTGCCTTCCCGAGCCTACCTCCGCCCCCAGGCTCTGCTAGCCTGCCACC
CAGCGGGGCGCCGCTGGGTCTGAGCACTTTTCTAGGAGCAGCGGTGTTCCGCCACCCAGCCTTCATCAGCCCAGCAT
TTGGCAGGCTCTTTTCCACTATGGCCCCCCTGACCAGCGCGTCGACTGCGGCTGCGCTCCTGAGACAGCCCACACCC
GCGGTGGAGGGCGCAGTGGCCTCGGGCGCGCTGGCCGACCCGGCCACCGCCGCGGCAGACAGACGCGCGTCAAGCAT
AGCCGCGCTGAGGCTCAAGGCTAAGGAGCACGCCGCGCAGCTCACGCAGCTCAACATCCTGCCGGGCACCAGCACGG
GGAAGGAGGTGTGCGATTACAAGGATGACGACGATAAGTGATAAACCCGCTGATCAGCCTCGACTGTGCCTTCTAGT
TGCCAGCCATCTGTTGTTTGCCCCTCCCCCGTGCCTTCCTTGACCCTGGAAGGTGCCACTCCCACTGTCCTTTCCTA
ATAAAATGAGGAAATTGCATCGCATTGTCTGAGTAGGTGTCATTCTATTCTGGGGGGTGGGGTGGGGCAGGACAGCA
AGGGGGAGGATTGGGAAGACAATAGCAGGCATGCTGGGGATGCGGTGGGCTCTATGGCTTCTGAGGCGGAAAGAACC
AGCTGGGGCTCTAGGGGGTATCCCCACGCGCCCTGTAGCGGCGCATTAAGCGCGGCGGGTGTGGTGGTTACGCGCAG
CGTGACCGCTACACTTGCCAGCGCCCTAGCGCCCGCTCCTTTCGCTTTCTTCCCTTCCTTTCTCGCCACGTTCGCCG
GCTTTCCCCGTCAAGCTCTAAATCGGGGGCTCCCTTTAGGGTTCCGATTTAGTGCTTTACGGCACCTCGACCCCAAA
AAACTTGATTAGGGTGATGGTTCACGTAGTGGGCCATCGCCCTGATAGACGGTTTTTCGCCCTTTGACGTTGGAGTC

CACGTTCTTTAATAGTGGACTCTTGTTCCAAACTGGAACAACACTCAACCCTATCTCGGTCTATTCTTTTGATTTAT
AAGGGATTTTGCCGATTTCGGCCTATTGGTTAAAAAATGAGCTGATTTAACAAAAATTTAACGCGAATTAATTCTGT
GGAATGTGTGTCAGTTAGGGTGTGGAAAGTCCCCAGGCTCCCCAGCAGGCAGAAGTATGCAAAGCATGCATCTCAAT
TAGTCAGCAACCAGGTGTGGAAAGTCCCCAGGCTCCCCAGCAGGCAGAAGTATGCAAAGCATGCATCTCAATTAGTC
AGCAACCATAGTCCCGCCCCTAACTCCGCCCATCCCGCCCCTAACTCCGCCCAGTTCCGCCCATTCTCCGCCCCATG
GCTGACTAATTTTTTTTATTTATGCAGAGGCCGAGGCCGCCTCTGCCTCTGAGCTATTCCAGAAGTAGTGAGGAGGC
TTTTTTGGAGGCCTAGGCTTTTGCAAAAAGCTCCCGGGAGCTTGTATATCCATTTTCGGATCTGATCAAGAGACAGG
ATGAGGATCGTTTCGCATGATTGAACAAGATGGATTGCACGCAGGTTCTCCGGCCGCTTGGGTGGAGAGGCTATTCG
GCTATGACTGGGCACAACAGACAATCGGCTGCTCTGATGCCGCCGTGTTCCGGCTGTCAGCGCAGGGGCGCCCGGTT
CTTTTTGTCAAGACCGACCTGTCCGGTGCCCTGAATGAACTGCAGGACGAGGCAGCGCGGCTATCGTGGCTGGCCAC
GACGGGCGTTCCTTGCGCAGCTGTGCTCGACGTTGTCACTGAAGCGGGAAGGGACTGGCTGCTATTGGGCGAAGTGC
CGGGGCAGGATCTCCTGTCATCTCACCTTGCTCCTGCCGAGAAAGTATCCATCATGGCTGATGCAATGCGGCGGCTG
CATACGCTTGATCCGGCTACCTGCCCATTCGACCACCAAGCGAAACATCGCATCGAGCGAGCACGTACTCGGATGGA
AGCCGGTCTTGTCGATCAGGATGATCTGGACGAAGAGCATCAGGGGCTCGCGCCAGCCGAACTGTTCGCCAGGCTCA
AGGCGCGCATGCCCGACGGCGAGGATCTCGTCGTGACCCATGGCGATGCCTGCTTGCCGAATATCATGGTGGAAAAT
GGCCGCTTTTCTGGATTCATCGACTGTGGCCGGCTGGGTGTGGCGGACCGCTATCAGGACATAGCGTTGGCTACCCG
TGATATTGCTGAAGAGCTTGGCGGCGAATGGGCTGACCGCTTCCTCGTGCTTTACGGTATCGCCGCTCCCGATTCGC
AGCGCATCGCCTTCTATCGCCTTCTTGACGAGTTCTTCTGAGCGGGACTCTGGGGTTCGAAATGACCGACCAAGCGA
CGCCCAACCTGCCATCACGAGATTTCGATTCCACCGCCGCCTTCTATGAAAGGTTGGGCTTCGGAATCGTTTTCCGG
GACGCCGGCTGGATGATCCTCCAGCGCGGGGATCTCATGCTGGAGTTCTTCGCCCACCCCAACTTGTTTATTGCAGC
TTATAATGGTTACAAATAAAGCAATAGCATCACAAATTTCACAAATAAAGCATTTTTTTCACTGCATTCTAGTTGTG
GTTTGTCCAAACTCATCAATGTATCTTATCATGTCTGTATACCGTCGACCTCTAGCTAGAGCTTGGCGTAATCATGG
TCATAGCTGTTTCCTGTGTGAAATTGTTATCCGCTCACAATTCCACACAACATACGAGCCGGAAGCATAAAGTGTAA
AGCCTGGGGTGCCTAATGAGTGAGCTAACTCACATTAATTGCGTTGCGCTCACTGCCCGCTTTCCAGTCGGGAAACC
TGTCGTGCCAGCTGCATTAATGAATCGGCCAACGCGCGGGGAGAGGCGGTTTGCGTATTGGGCGCTCTTCCGCTTCC
TCGCTCACTGACTCGCTGCGCTCGGTCGTTCGGCTGCGGCGAGCGGTATCAGCTCACTCAAAGGCGGTAATACGGTT
ATCCACAGAATCAGGGGATAACGCAGGAAAGAACATGTGAGCAAAAGGCCAGCAAAAGGCCAGGAACCGTAAAAAGG
CCGCGTTGCTGGCGTTTTTCCATAGGCTCCGCCCCCCTGACGAGCATCACAAAAATCGACGCTCAAGTCAGAGGTGG
CGAAACCCGACAGGACTATAAAGATACCAGGCGTTTCCCCCTGGAAGCTCCCTCGTGCGCTCTCCTGTTCCGACCCT
GCCGCTTACCGGATACCTGTCCGCCTTTCTCCCTTCGGGAAGCGTGGCGCTTTCTCATAGCTCACGCTGTAGGTATC
TCAGTTCGGTGTAGGTCGTTCGCTCCAAGCTGGGCTGTGTGCACGAACCCCCCGTTCAGCCCGACCGCTGCGCCTTA
TCCGGTAACTATCGTCTTGAGTCCAACCCGGTAAGACACGACTTATCGCCACTGGCAGCAGCCACTGGTAACAGGAT
TAGCAGAGCGAGGTATGTAGGCGGTGCTACAGAGTTCTTGAAGTGGTGGCCTAACTACGGCTACACTAGAAGAACAG
TATTTGGTATCTGCGCTCTGCTGAAGCCAGTTACCTTCGGAAAAAGAGTTGGTAGCTCTTGATCCGGCAAACAAACC
ACCGCTGGTAGCGGTGGTTTTTTTGTTTGCAAGCAGCAGATTACGCGCAGAAAAAAAGGATCTCAAGAAGATCCTTT
GATCTTTTCTACGGGGTCTGACGCTCAGTGGAACGAAAACTCACGTTAAGGGATTTTGGTCATGAGATTATCAAAAA
GGATCTTCACCTAGATCCTTTTAAATTAAAAATGAAGTTTTAAATCAATCTAAAGTATATATGAGTAAACTTGGTCT
GACAGTTACCAATGCTTAATCAGTGAGGCACCTATCTCAGCGATCTGTCTATTTCGTTCATCCATAGTTGCCTGACT
CCCCGTCGTGTAGATAACTACGATACGGGAGGGCTTACCATCTGGCCCCAGTGCTGCAATGATACCGCGAGACCCAC
GCTCACCGGCTCCAGATTTATCAGCAATAAACCAGCCAGCCGGAAGGGCCGAGCGCAGAAGTGGTCCTGCAACTTTA
TCCGCCTCCATCCAGTCTATTAATTGTTGCCGGGAAGCTAGAGTAAGTAGTTCGCCAGTTAATAGTTTGCGCAACGT
TGTTGCCATTGCTACAGGCATCGTGGTGTCACGCTCGTCGTTTGGTATGGCTTCATTCAGCTCCGGTTCCCAACGAT
CAAGGCGAGTTACATGATCCCCCATGTTGTGCAAAAAAGCGGTTAGCTCCTTCGGTCCTCCGATCGTTGTCAGAAGT

AAGTTGGCCGCAGTGTTATCACTCATGGTTATGGCAGCACTGCATAATTCTCTTACTGTCATGCCATCCGTAAGATG
CTTTTCTGTGACTGGTGAGTACTCAACCAAGTCATTCTGAGAATAGTGTATGCGGCGACCGAGTTGCTCTTGCCCGG
CGTCAATACGGGATAATACCGCGCCACATAGCAGAACTTTAAAAGTGCTCATCATTGGAAAACGTTCTTCGGGGCGA
AAACTCTCAAGGATCTTACCGCTGTTGAGATCCAGTTCGATGTAACCCACTCGTGCACCCAACTGATCTTCAGCATC
TTTTACTTTCACCAGCGTTTCTGGGTGAGCAAAAACAGGAAGGCAAAATGCCGCAAAAAAGGGAATAAGGGCGACAC
GGAAATGTTGAATACTCATACTCTTCCTTTTTCAATATTATTGAAGCATTTATCAGGGTTATTGTCTCATGAGCGGA
TACATATTTGAATGTATTTAGAAAAATAAACAAATAGGGGTTCCGCGCACATTTCCCCGAAAAGTGCCACCTGACGT
C

>pET14P

TTCTTGAAGACGAAAGGGCCTCGTGATACGCCTATTTTTATAGGTTAATGTCATGATAATAATGGTTTCTTAGACGT
CAGGTGGCACTTTTCGGGGAAATGTGCGCGGAACCCCTATTTGTTTATTTTTCTAAATACATTCAAATATGTATCCG
CTCATGAGACAATAACCCTGATAAATGCTTCAATAATATTGAAAAAGGAAGAGTATGAGTATTCAACATTTCCGTGT
CGCCCTTATTCCCTTTTTTGCGGCATTTTGCCTTCCTGTTTTTGCTCACCCAGAAACGCTGGTGAAAGTAAAAGATG
CTGAAGATCAGTTGGGTGCACGAGTGGGTTACATCGAACTGGATCTCAACAGCGGTAAGATCCTTGAGAGTTTTCGC
CCCGAAGAACGTTTTCCAATGATGAGCACTTTTAAAGTTCTGCTATGTGGCGCGGTATTATCCCGTGTTGACGCCGG
GCAAGAGCAACTCGGTCGCCGCATACACTATTCTCAGAATGACTTGGTTGAGTACTCACCAGTCACAGAAAAGCATC
TTACGGATGGCATGACAGTAAGAGAATTATGCAGTGCTGCCATAACCATGAGTGATAACACTGCGGCCAACTTACTT
CTGACAACGATCGGAGGACCGAAGGAGCTAACCGCTTTTTTGCACAACATGGGGGATCATGTAACTCGCCTTGATCG
TTGGGAACCGGAGCTGAATGAAGCCATACCAAACGACGAGCGTGACACCACGATGCCTGCAGCAATGGCAACAACGT
TGCGCAAACTATTAACTGGCGAACTACTTACTCTAGCTTCCCGGCAACAATTAATAGACTGGATGGAGGCGGATAAA
GTTGCAGGACCACTTCTGCGCTCGGCCCTTCCGGCTGGCTGGTTTATTGCTGATAAATCTGGAGCCGGTGAGCGTGG
GTCTCGCGGTATCATTGCAGCACTGGGGCCAGATGGTAAGCCCTCCCGTATCGTAGTTATCTACACGACGGGGAGTC
AGGCAACTATGGATGAACGAAATAGACAGATCGCTGAGATAGGTGCCTCACTGATTAAGCATTGGTAACTGTCAGAC
CAAGTTTACTCATATATACTTTAGATTGATTTAAAACTTCATTTTTAATTTAAAAGGATCTAGGTGAAGATCCTTTT
TGATAATCTCATGACCAAAATCCCTTAACGTGAGTTTTCGTTCCACTGAGCGTCAGACCCCGTAGAAAAGATCAAAG
GATCTTCTTGAGATCCTTTTTTTCTGCGCGTAATCTGCTGCTTGCAAACAAAAAAACCACCGCTACCAGCGGTGGTT
TGTTTGCCGGATCAAGAGCTACCAACTCTTTTTCCGAAGGTAACTGGCTTCAGCAGAGCGCAGATACCAAATACTGT
CCTTCTAGTGTAGCCGTAGTTAGGCCACCACTTCAAGAACTCTGTAGCACCGCCTACATACCTCGCTCTGCTAATCC
TGTTACCAGTGGCTGCTGCCAGTGGCGATAAGTCGTGTCTTACCGGGTTGGACTCAAGACGATAGTTACCGGATAAG
GCGCAGCGGTCGGGCTGAACGGGGGGTTCGTGCACACAGCCCAGCTTGGAGCGAACGACCTACACCGAACTGAGATA
CCTACAGCGTGAGCTATGAGAAAGCGCCACGCTTCCCGAAGGGAGAAAGGCGGACAGGTATCCGGTAAGCGGCAGGG
TCGGAACAGGAGAGCGCACGAGGGAGCTTCCAGGGGGAAACGCCTGGTATCTTTATAGTCCTGTCGGGTTTCGCCAC
CTCTGACTTGAGCGTCGATTTTTGTGATGCTCGTCAGGGGGGCGGAGCCTATGGAAAAACGCCAGCAACGCGGCCTT
TTTACGGTTCCTGGCCTTTTGCTGGCCTTTTGCTCACATGTTCTTTCCTGCGTTATCCCCTGATTCTGTGGATAACC
GTATTACCGCCTTTGAGTGAGCTGATACCGCTCGCCGCAGCCGAACGACCGAGCGCAGCGAGTCAGTGAGCGAGGAA
GCGGAAGAGCGCCTGATGCGGTATTTTCTCCTTACGCATCTGTGCGGTATTTCACACCGCATATATGGTGCACTCTC
AGTACAATCTGCTCTGATGCCGCATAGTTAAGCCAGTATACACTCCGCTATCGCTACGTGACTGGGTCATGGCTGCG
CCCCGACACCCGCCAACACCCGCTGACGCGCCCTGACGGGCTTGTCTGCTCCCGGCATCCGCTTACAGACAAGCTGT
GACCGTCTCCGGGAGCTGCATGTGTCAGAGGTTTTCACCGTCATCACCGAAACGCGCGAGGCAGCTGCGGTAAAGCT
CATCAGCGTGGTCGTGAAGCGATTCACAGATGTCTGCCTGTTCATCCGCGTCCAGCTCGTTGAGTTTCTCCAGAAGC
GTTAATGTCTGGCTTCTGATAAAGCGGGCCATGTTAAGGGCGGTTTTTTCCTGTTTGGTCACTGATGCCTCCGTGTA
AGGGGGATTTCTGTTCATGGGGGTAATGATACCGATGAAACGAGAGAGGATGCTCACGATACGGGTTACTGATGATG
AACATGCCCGGTTACTGGAACGTTGTGAGGGTAAACAACTGGCGGTATGGATGCGGCGGGACCAGAGAAAAATCACT
CAGGGTCAATGCCAGCGCTTCGTTAATACAGATGTAGGTGTTCCACAGGGTAGCCAGCAGCATCCTGCGATGCAGAT
CCGGAACATAATGGTGCAGGGCGCTGACTTCCGCGTTTCCAGACTTTACGAAACACGGAAACCGAAGACCATTCATG
TTGTTGCTCAGGTCGCAGACGTTTTGCAGCAGCAGTCGCTTCACGTTCGCTCGCGTATCGGTGATTCATTCTGCTAA
CCAGTAAGGCAACCCCGCCAGCCTAGCCGGGTCCTCAACGACAGGAGCACGATCATGCGCACCCGTGGCCAGGACCC
AACGCTGCCCGAGATGCGCCGCGTGCGGCTGCTGGAGATGGCGGACGCGATGGATATGTTCTGCCAAGGGTTGGTTT
GCGCATTCACAGTTCTCCGCAAGAATTGATTGGCTCCAATTCTTGGAGTGGTGAATCCGTTAGCGAGGTGCCGCCGG
CTTCCATTCAGGTCGAGGTGGCCCGGCTCCATGCACCGCGACGCAACGCGGGGAGGCAGACAAGGTATAGGGCGGCG
CCTACAATCCATGCCAACCCGTTCCATGTGCTCGCCGAGGCGGCATAAATCGCCGTGACGATCAGCGGTCCAGTGAT

CGAAGTTAGGCTGGTAAGAGCCGCGAGCGATCCTTGAAGCTGTCCCTGATGGTCGTCATCTACCTGCCTGGACAGCA
TGGCCTGCAACGCGGGCATCCCGATGCCGCCGGAAGCGAGAAGAATCATAATGGGGAAGGCCATCCAGCCTCGCGTC
GCGAACGCCAGCAAGACGTAGCCCAGCGCGTCGGCCGCCATGCCGGCGATAATGGCCTGCTTCTCGCCGAAACGTTT
GGTGGCGGGACCAGTGACGAAGGCTTGAGCGAGGGCGTGCAAGATTCCGAATACCGCAAGCGACAGGCCGATCATCG
TCGCGCTCCAGCGAAAGCGGTCCTCGCCGAAAATGACCCAGAGCGCTGCCGGCACCTGTCCTACGAGTTGCATGATA
AAGAAGACAGTCATAAGTGCGGCGACGATAGTCATGCCCCGCGCCCACCGGAAGGAGCTGACTGGGTTGAAGGCTCT
CAAGGGCATCGGTCGACGCTCTCCCTTATGCGACTCCTGCATTAGGAAGCAGCCCAGTAGTAGGTTGAGGCCGTTGA
GCACCGCCGCCGCAAGGAATGGTGCATGCAAGGAGATGGCGCCCAACAGTCCCCCGGCCACGGGGCCTGCCACCATA
CCCACGCCGAAACAAGCGCTCATGAGCCCGAAGTGGCGAGCCCGATCTTCCCCATCGGTGATGTCGGCGATATAGGC
GCCAGCAACCGCACCTGTGGCGCCGGTGATGCCGGCCACGATGCGTCCGGCGTAGAGGATCGAGATCTCGATCCCGC
GAAATTAATACGACTCACTATAGGGAGACCACAACGGTTTCCCTCTAGAAATAATTTTGTTTAACTTTAAGAAGGAG
ATATACCATGGGCAGCAGCCATCATCATCATCATCACAGCAGCGGCCTGGAAGTTCTGTTCCAGGGGCCCGCGGCCG
CACATATGGGATCCCTCGAGTAAGCTGCTAACAAAGCCCGAAAGGAAGCTGAGTTGGCTGCTGCCACCGCTGAGCAA
TAACTAGCATAACCCCTTGGGGCCTCTAAACGGGTCTTGAGGGGTTTTTTGCTGAAAGGAGGAACTATATCCGGATA
TCCACAGGACGGGTGTGGTCGCCATGATCGCGTAGTCGATAGTGGCTCCAAGTAGCGAAGCGAGCAGGACTGGGCGG
CGGCCAAAGCGGTCGGACAGTGCTCCGAGAACGGGTGCGCATAGAAATTGCATCAACGCATATAGCGCTAGCAGCAC
GCCATAGTGACTGGCGATGCTGTCGGAATGGACGATATCCCGCAAGAGGCCCGGCAGTACCGGCATAACCAAGCCTA
TGCCTACAGCATCCAGGGTGACGGTGCCGAGGATGACGATGAGCGCATTGTTAGATTTCATACACGGTGCCTGACTG
CGTTAGCAATTTAACTGTGATAAACTACCGCATTAAAGCTTATCGATGATAAGCTGTCAAACATGAGAA

Oligos

>Q50 P3 site

TGTGTCTTAATTAGATTAACGCACTTGATAGTGCGGGCGTGGCT

>Q50 P4 site

TGTGTCTTAATTAGGATTAACGCACTTGATAGTGCGGGCGTGGCT

>K50 P3 site

TGTGTCTTAATCCGATTAACGCACTTGATAGTGCGGGCGTGGCT

>K50 P4 site

TGTGTCTTAATCCGGATTAACGCACTTGATAGTGCGGGCGTGGCT
